# Identifying opportunities for high resolution pesticide usage data to improve the efficiency of endangered species pesticide risk assessment

**DOI:** 10.1101/2022.09.07.507030

**Authors:** Steffen E. Eikenberry, Gwen Iacona, Erin L. Murphy, Greg Watson, Leah R. Gerber

## Abstract

The US pesticide registration and review process requires regular re-assessment of the risk of pesticide use, including risk to species listed under the Endangered Species Act (ESA). Yet, current methods for assessment were not designed to assess threats that could involve hundreds of pesticides potentially impacting multiple species across a continent. Thus, many pesticides remain on the market without complete review under ESA. A promising approach to address this challenge calls for the use of high-resolution data in the spatial analyses used to determine the likelihood of adverse effect. However, such data are rare, and implementing data collection initiatives are difficult and expensive. Here we examine the extent to which increased data resolution could result in significant gains in efficiency to the pesticide risk assessment process. To do so, we employ a modified value-of-information (VOI) approach. By using data available only in California, we found that high resolution data increased the number of species deemed not likely to be adversely affected by pesticides from less than 5% to nearly 50%. Across the contiguous US, we predicted that 48% of species would be deemed not likely to be adversely affected using high resolution data, compared to 20% without. However, if such data were available in just 11 states, 68% of the available gains in efficiency could be obtained. Therefore, a focused collection or release of high-resolution pesticide usage data could expedite pesticide risk assessment and reduce endangered species’ risk of pesticide impacts.

**Significance:** Regulation of environmental risk must balance avoidance of harm with assessment efficiency. The current pesticide registration and review process in the U.S. is generally considered ineffective in assessing impact on species listed under the Endangered Species Act, due to the vast scale of the problem. We assessed the value of using high resolution pesticide usage data in the risk assessment process to identify cost-effective opportunities to rapidly improve this assessment process and ensure that ESA listed species are protected while accounting for agricultural production. Our results suggest that using existing high-resolution data in California and collection of such information from 11 other states could reduce the risk assessment burden across the contiguous U.S. by one-quarter.

## Introduction

Agricultural pesticide use illustrates a common environmental risk assessment conundrum. To protect crop yield and quality, pesticides prevent or mitigate impact by killing or repelling target pest organisms. Existing regulations require that crop protection benefits be balanced against potential negative environmental impacts by conducting risk assessments as part of the pesticide review process. Such review must reasonably identify the potential impacts of assessed pesticides, but also be feasible to perform in a timely manner. This second criterion has hindered effective pesticide risk assessment in the U.S. (1-3).

In the U.S., pesticide use must meet a “no unreasonable adverse effects on the environment” standard under the Federal Insecticide, Fungicide and Rodenticide Act (FIFRA) (4). The U.S. Endangered Species Act also requires that pesticides do not negatively impact the persistence of threatened and endangered species (ESA) (1). The differing obligations of these legislative acts have made the current pesticide registration and review process time intensive with a lack of transparency and confidence among stakeholders. Hundreds of pesticides remain on the market without complete review under the ESA (2,3,5). Risk of pesticide use is mandated to be assessed at initial pesticide registration and on 15-year review intervals; risk to ESA listed species is intended to be part of the periodic review. The US Environmental Protection Agency Office of Pesticide Programs (EPA OPP) performs these evaluations and, when deemed necessary, engages in consultation with the US Fish and Wildlife and National Marine Fisheries Service (the Services).

In recent biological evaluations for ESA-listed species, EPA has determined that most endangered species may have the potential for exposure and have sent large lists of species for consultation to the Services (6,7). The consultation process under the ESA (Section 7) was not designed for assessment of continent-wide risks to potentially every species. The Services therefore face an overwhelming bottleneck in the risk assessment process, with hundreds of endangered species determinations incomplete. Stakeholders across the public and private sectors agree that improving this process is a national priority to more effectively minimize the risk of pesticide exposure to ESA listed species and to avoid unnecessary impacts on agriculture (2,3).

Improving the spatial data underlying the exposure evaluation could provide an immediate opportunity for expediting the endangered species evaluation process while enabling compliance with ESA (2,3,8,9). Under current EPA methods, initial screening of pesticide risk considers whether listed species ranges intersect regions where exposure is possible. If more than 1% of the range intersects areas where pesticide is potentially used (analysis process detailed in Section 9 of SI), species are deemed likely to be adversely affected (LAA) and a full Section 7 consultation is required. Inadequate resolution of the data used in the EPA screening process is frequently touted as a roadblock to pesticide risk assessment efficiency because at present, most species are sent for consultation (2,6). Our prior analysis showed that substituting high spatial resolution pesticide usage data for a single agent (carbaryl) reduced the number of California plant species requiring a full consultation for that agent by almost half (2). This suggests that the potential value to decision makers of substituting high resolution data can be high and that improving the spatial data underlying the exposure evaluation could provide an immediate opportunity for expediting the endangered species evaluation process and enable compliance with ESA while supporting agricultural production.

High resolution data are not currently available nationwide. Township resolution usage data (approximately 36 square miles) are only available for CA, and obtaining such data elsewhere would be politically difficult and expensive. Therefore, efforts to collect/release these data for other regions of the US should consider the potential for efficiencies in pesticide review with high resolution data.

Value of information (VOI) analyses offer a promising approach to inform data needs for conservation decision support (10-12). VOI approaches facilitate consideration of the implications of collecting new data on the expected outcome of a decision. VOI represents a powerful way to consider how to balance the expected cost of updating the underlying data with the expected benefits to the decision outcomes. Such an approach is particularly helpful in situations where the context is controversial, the options are not clear, the need is great but capacity is limited, and slow decision making hinders optimal outcomes.

We applied a VOI analysis approach to consider how high-resolution pesticide usage data for a wide range of agents (223 individual pesticides, representing >99% of nationally applied insecticide, herbicide, and fungicide by mass) could be expected to influence risk assessment across ESA listed species. We examined the implications of using township resolution pesticide usage data to assess whether 76 ESA listed animal species ranges and 173 ESA listed plant species ranges endemic to CA were likely to be determined not likely to be adversely affected (NLAA) for the pesticides under an EPA biological evaluation. The potential benefit of township resolution usage data derives from its ability to switch a species from LAA to NLAA designation if the species range has <1% overlap with the township resolution usage data but a larger overlap with lower resolution usage data (i.e., county or crop reporting district resolution data).

We then extrapolated our results to a national scale (contiguous U.S.) to identify regions where higher resolution data may improve screening results. We evaluated the screening utility of existing crop reporting district (CRD) and county level national usage estimates across 577 ESA listed animal ranges and 451 ESA listed plant ranges. We then used a regression/classification analysis, trained on the high-resolution California data, to identify regions where sub-county resolution data may be most valuable. We then quantified if and where high-resolution usage data could bring efficiency to the pesticide risk assessment process.

## Results

We found that about half the listed species in California would be deemed likely to be adversely affected by pesticide use if township resolution usage data were used for risk assessment, compared with 5% under CRD resolution data (roughly analogous to EPA’s current approach). Further efficiencies could be achieved if such data were available in specific regions elsewhere in the US.

### Impact of township level usage data in assessing California species risk

Using township resolution pesticide usage data provided an immediate source of risk assessment efficiency for California species. It markedly increased the number of plant and animal species ranges that could potentially be designated NLAA with <1% overlap with pesticide usage. Across 185 pesticides (all for which there was some usage in CA), and all plants (173) and animals (76), about 5%, 17%, and 47% of pesticide/range pairings yielded <1% overlap under CRD, county, and township usage resolution, respectively (Fig. 1, Table S1). These results suggest a disproportionate benefit when usage data is available at township scale versus CRD or county scale; in contrast, the current approach used by OPP has led to LAA determinations for >90% of species nationally in several biological opinions (7). In general, plants were more likely to be NLAA at all usage resolutions, and <1% overlap was most common across herbicides (Fig. 1, additional details in Figs. S20-S32 and Table S1). Results for 30 pesticides are given in Figs. S26-S31, and results for all pesticides can be visualized in detail using an iPython app available via the Dryad repository (see also Fig. S32).

**Figure 1.**
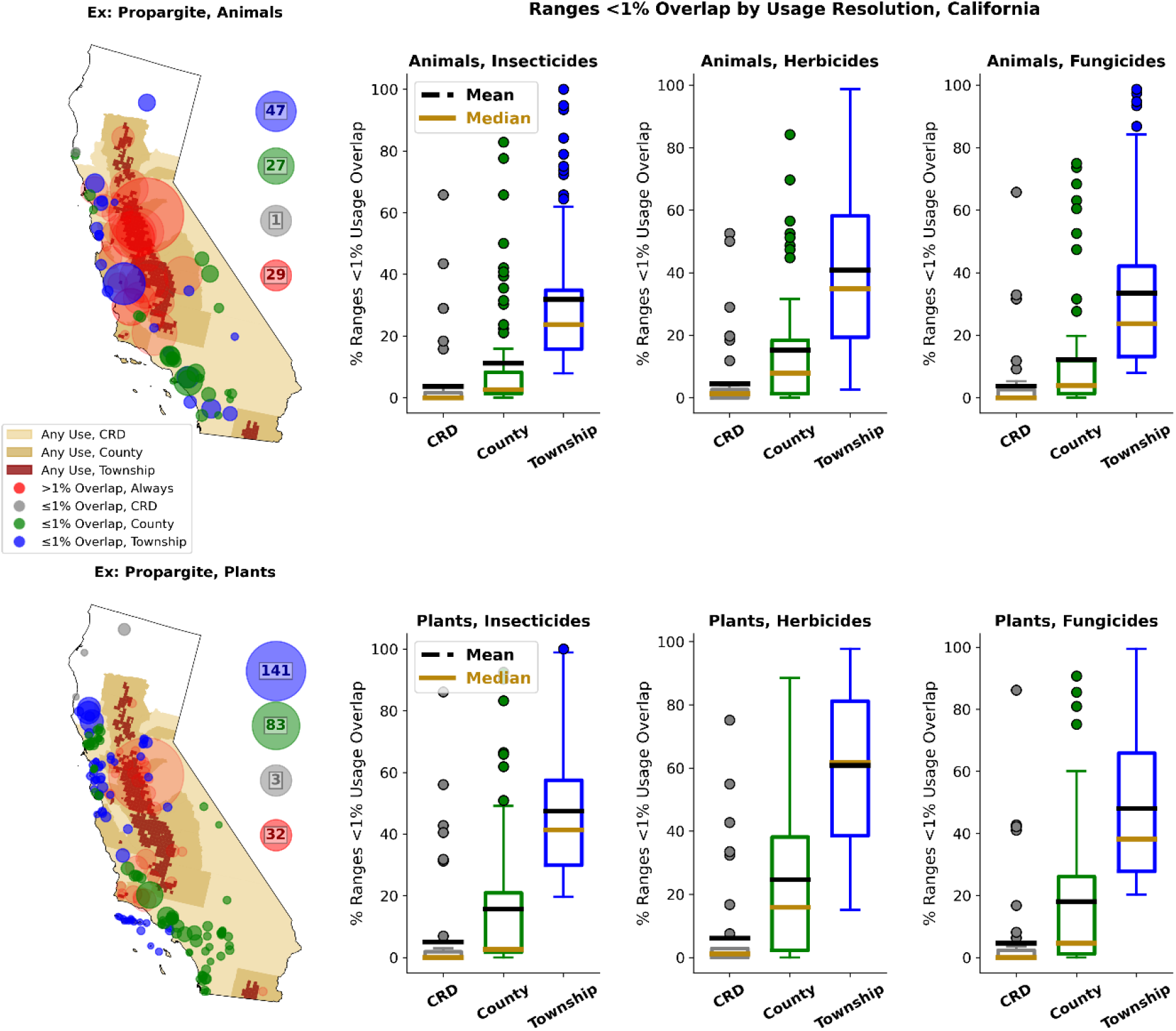
Summary of relationship between pesticide usage resolution and the number of species’ ranges with <1% overlap with any pesticide usage, for 76 animal and 173 plant ranges restricted to California, and across 223 pesticides divided into insecticide, herbicide, and fungicide classes. The maps on the left give example spatial distributions of usage (any) at either CRD, county, and township resolutions, for the insecticide propargite, along with ranges reprsented by bubbles that scale with range area, coded according to whether <1% usage occurs at no usage resolution, at CRD, at county, and at township. Note that if there is <1% overlap at one resolution, this necessarily occurs at all finer resolutions. The circles give the total number of ranges in each overlap category. The six sets of box plots on the right summarize results across all pesticides. In general, there is a relatively small benefit to increasing usage resolution from CRD to county scale, with a larger benefit to township usage data. Note that usage represents the maximum over 2013 through 2017, where this is calculated separately for each base geometric object (township, county, or CRD).

### Impact of higher resolution usage data to assess species risk across the other 47 states

Township resolution data is currently unavailable outside California, so we first explored where existing medium-resolution data might be of value. We found that using county instead of CRD resolution data could yield a modest increase in the number of ranges that are determined to be NLAA. For the lower 48 states outside California, 26% of range/pesticide pairings were found to be NLAA with county usage data compared to roughly 20% with CRD-level data (Fig 2). These gains for county resolution data appeared to depend on large-scale regional usage trends and increases in spatial resolution affected overlap results mainly at the borders between regions of high and little to no usage, such as in California, the southwestern US, southern Appalachia, and Florida (Figs. S49 and S50).

**Figure 2.**
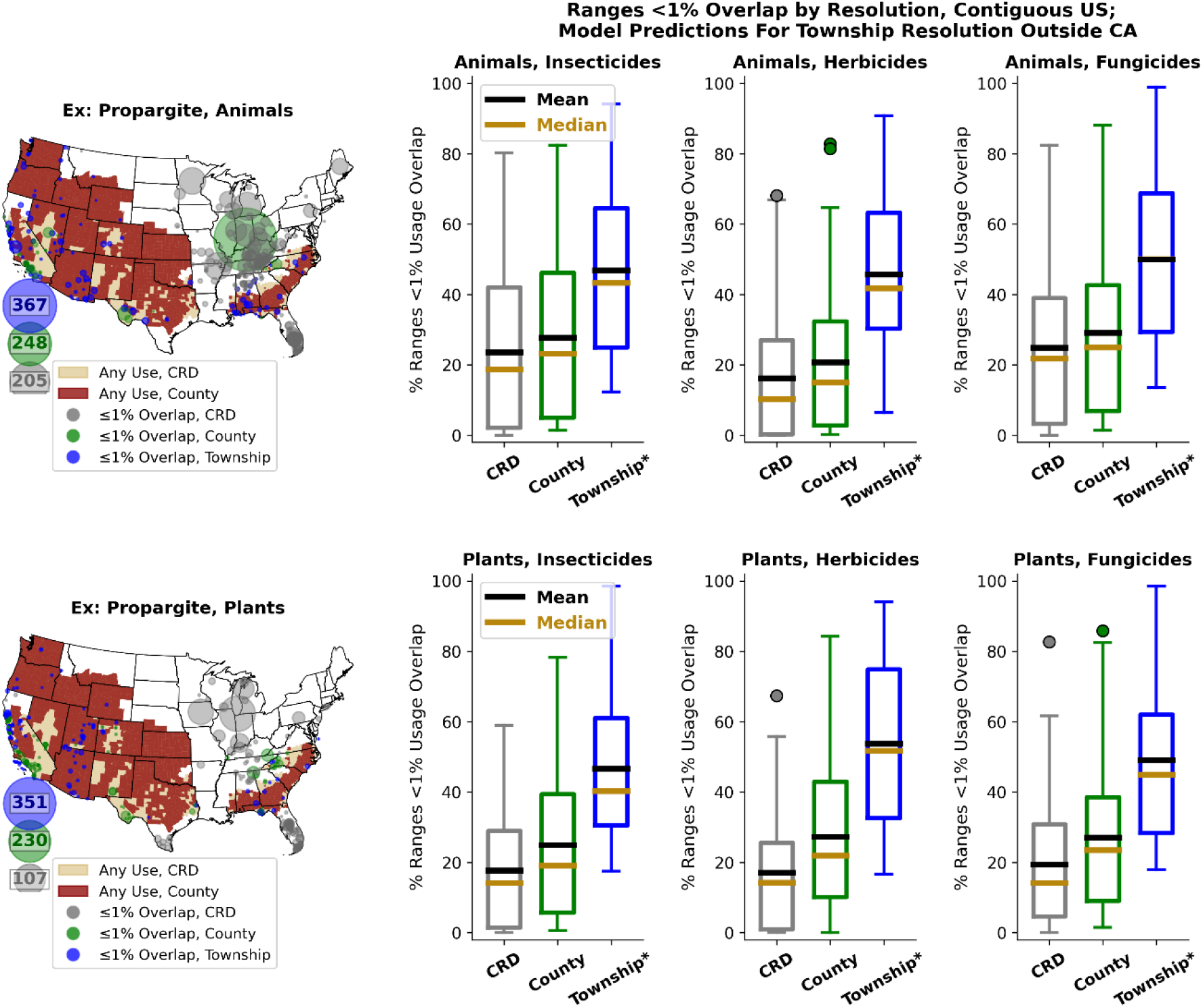
Summary of how usage resolution is predicted to affect the number of species’ ranges with <1% overlap with any pesticide usage, for 577 animal and 451 plant ranges restricted to the contiguous US. Direct overlap analysis is used for CRD and county results, as well as township results for ranges strictly contained within California, while model predictions are used for township resolution outside of California. Results are given for the 184 pesticides for which predictions on township usage outside CA could be made.

However, we were particularly interested in the potential value of collecting or releasing township resolution data in regions where it is not currently available. Thus, we used logistic regression models to extrapolate the impact of township resolution usage data from CA to the rest of the contiguous US. Across the 184 pesticides for which models could be fit, we predicted that, compared to CRD resolution, 28.34% of all range/pesticide pairings would switch to NLAA if township usage data were available (and so 48.48% of range/pesticide pairs would be NLAA in total) (Fig 2, Table 1). These models were fit using CA data with range area, agricultural land use fraction in range (“agricultural fraction”), and mean pesticide exposure at county scale (in kg m^-2^) as the predictors of whether a range was likely to benefit from township data. The final models for animals and plants had mean area under the curve (AUC) ± SDs of 86.12 ± 7.04% and 77.81 ± 8.43%, respectively, which may be regarded as “excellent” and “acceptable” performance, respectively (14). Note the AUC is a measure of classifier performance, with 0.5 representing random chance and 1 a perfect classifier. Figs S33-S36 and Table S2 give further details on model performance. The predicted marginal benefit of township-level data was to designate an additional 22.27% (28.34%) of all species/pesticide pairs to be NLAA over county-level (CRD-level) data (absolute benefit; see Table 1). However, the distribution of effect was geographically heterogeneous (Fig. 3; see also Figs. S37-S47 and iPython app available at (13)).

**Table 1.**
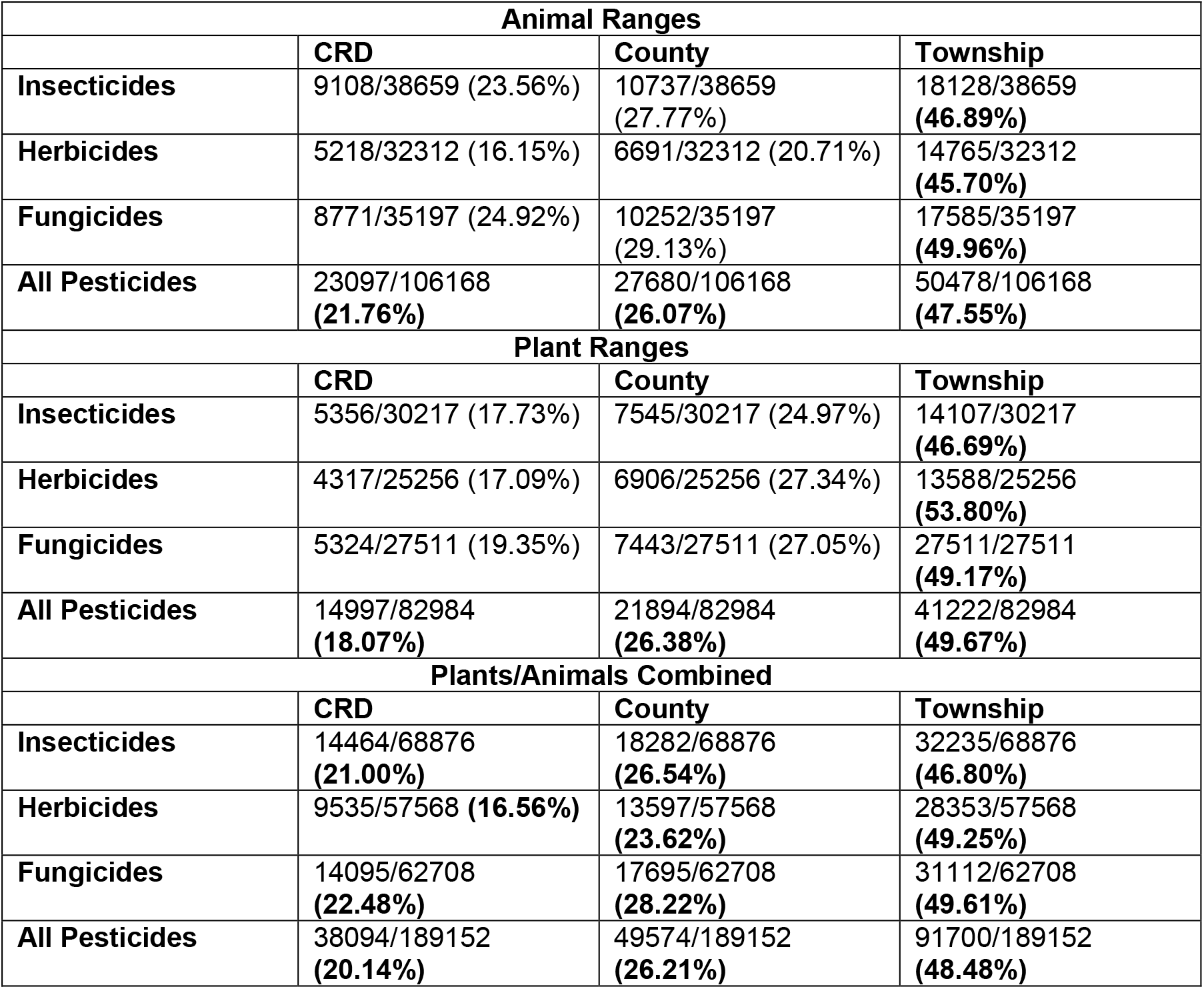
Total number of range/pesticide pairs predicted to be NLAA (<1% range/pesticide overlap) under different pesticide usage resolutions, for the entire contiguous US, and for the 184 pesticides for which township benefit predictions could be made outside of California. Cells give the number of range/pesticide pairs NLAA out of the total, with the percentage in parentheses. Results are disaggregated by pesticide class, and are presented for animals, plants, and animals/plants combined.

| **Animal Ranges** |  |  |  |
| --- | --- | --- | --- |
|  | **CRD** | **County** | **Township** |
| **Insecticides** | 9108/38659 (23.56%) | 10737/38659<br>(27.77%) | 18128/38659<br>**(46.89%)** |
| **Herbicides** | 5218/32312 (16.15%) | 6691/32312 (20.71%) | 14765/32312<br>**(45.70%)** |
| **Fungicides** | 8771/35197 (24.92%) | 10252/35197<br>(29.13%) | 17585/35197<br>**(49.96%)** |
| **All Pesticides** | 23097/106168<br>**(21.76%)** | 27680/106168<br>**(26.07%)** | 50478/106168<br>**(47.55%)** |
| **Plant Ranges** |  |  |  |
|  | **CRD** | **County** | **Township** |
| **Insecticides** | 5356/30217 (17.73%) | 7545/30217 (24.97%) | 14107/30217<br>**(46.69%)** |
| **Herbicides** | 4317/25256 (17.09%) | 6906/25256 (27.34%) | 13588/25256<br>**(53.80%)** |
| **Fungicides** | 5324/27511 (19.35%) | 7443/27511 (27.05%) | 27511/27511<br>**(49.17%)** |
| **All Pesticides** | 14997/82984<br>**(18.07%)** | 21894/82984<br>**(26.38%)** | 41222/82984<br>**(49.67%)** |
| **Plants/Animals Combined** |  |  |  |
|  | **CRD** | **County** | **Township** |
| **Insecticides** | 14464/68876<br>**(21.00%)** | 18282/68876<br>**(26.54%)** | 32235/68876<br>**(46.80%)** |
| **Herbicides** | 9535/57568 **(16.56%)** | 13597/57568<br>**(23.62%)** | 28353/57568<br>**(49.25%)** |
| **Fungicides** | 14095/62708<br>**(22.48%)** | 17695/62708<br>**(28.22%)** | 31112/62708<br>**(49.61%)** |
| **All Pesticides** | 38094/189152<br>**(20.14%)** | 49574/189152<br>**(26.21%)** | 91700/189152<br>**(48.48%)** |

**Figure 3.**
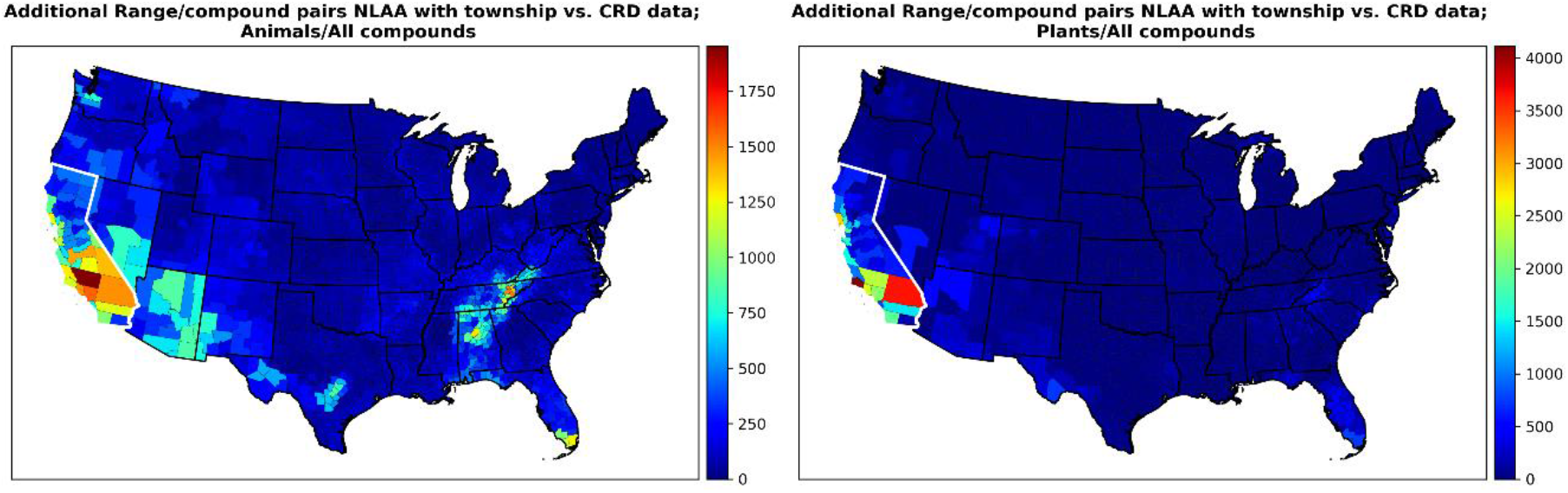
The number of range/pesticide pairs predicted to switch from an LAA to NLAA classification if township-level data is available compared to a baseline where only CRD-level data is available, for each county. The left panel gives results for animals, while the right panel shows results for plants. California is highlighted to emphasize that, for species restricted strictly to CA, predictions are determined using direct overlap calculations, while the classification model described in the text is used outside CA. This model is also used for ranges that overlap CA but are not restricted solely to that state.

### Priority states for high-resolution (township level) usage data

We used two iterative procedures (see Methods for details) to identify 11 individual states (beyond CA) that could be prioritized to achieve the greatest benefits from township resolution data collection. We found that if township usage data were made available for FL, AL, TX, TN, NV, AZ, UT, GA, NC, OR, and WA, in that order of priority, then 65.89% of the additional animal range/pesticide pairs and 72.51% of the additional plant range/pesticide pairs that *could* benefit from township data, in fact *would* benefit. Combined, 68.08% of all range/pesticide pairs that *could* potentially shift to an NLAA classification would do so. That is, outside of CA, 34,404 range/pesticide pairs *could* shift to NLAA if township resolution data were available across the contiguous US., and 23,484 (68.08%) of these shifts *would* be realized with data from just these 11 states. Fig. 4 summarizes the priority states identified by the two iterative methods. The 11 final priority states are highlighted in Fig. S55.

**Figure 4.**
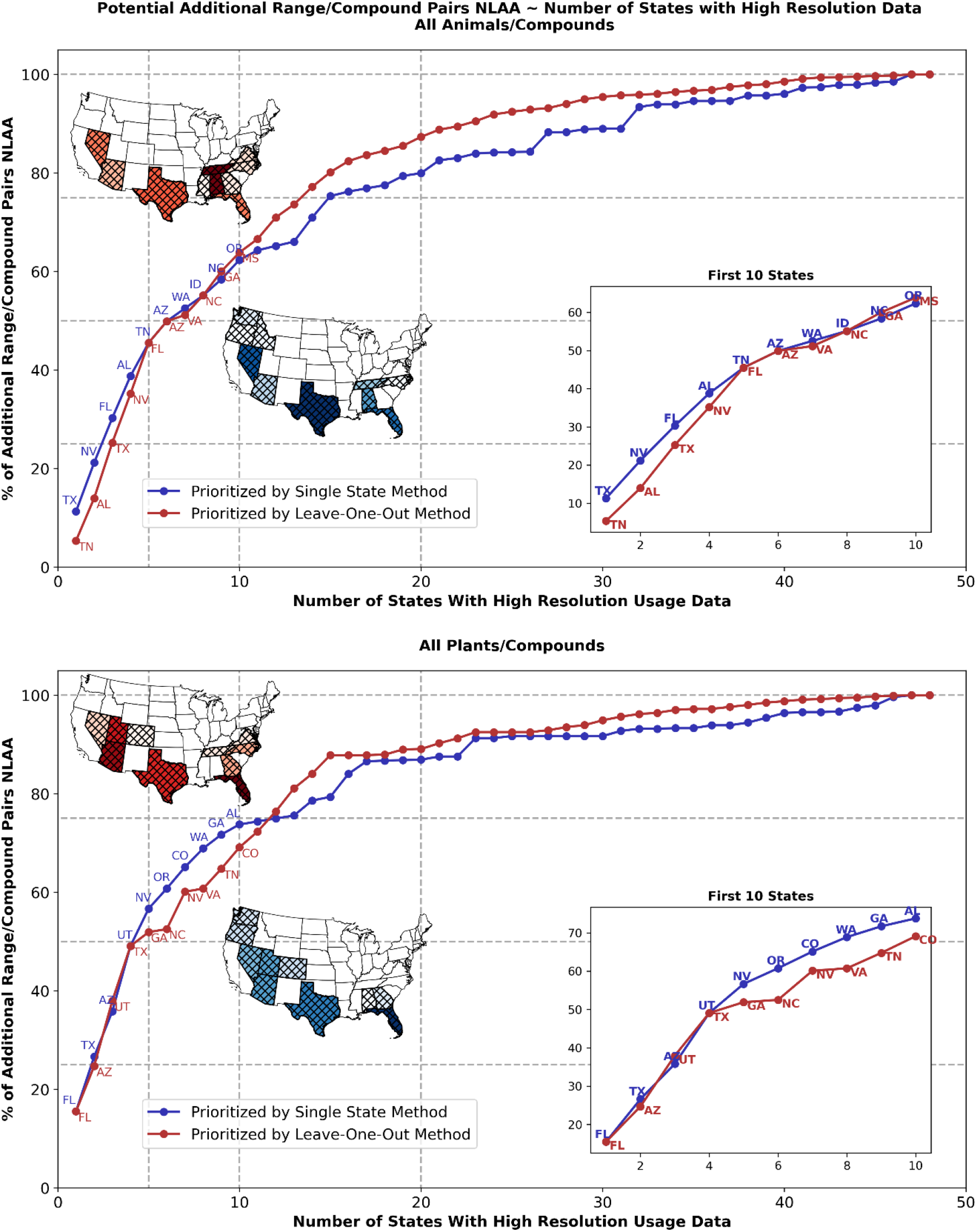
Priority states for township resolution data identified by either a single state or “leave-one-out” analysis. Animals (top panel) and plants (bottom panel) are shown separately. Each panel gives the percentage of potential benefit that can be realized across the contiguous US, in terms of additional species ranges designated NLAA, as a function of the number of states for which data is available. States are added in order of priority identified by the two aforementioned methods. Inset maps and plot indicate the top ten states under each method.

## Discussion

VOI analysis represents a promising approach to address the vast need for bringing efficiency to pesticide risk assessment for endangered species in the U.S. The current approach is likely untenable: 223 individual pesticides on 1,028 endangered species ranges in the contiguous U.S. yields 229,244 range/pesticide pairings that must be evaluated for potential harm. Using current methods for risk assessment, EPA sends most assessments to the Services for a Biological Evaluation as mandated by Section 7, which is onerous and time-consuming, often taking multiple years to complete. The percentage of LAA species determinations in recent OPP biological opinions ranges from 56% (atrazine) to 93% (glyphosate) (6,7). This status quo represents an overwhelming resource demand, particularly for the Services, and leaves pesticides on the market without a full assessment under ESA. Therefore, we examined the potential application of VOI to streamline the pesticide review process. Our results suggest that using township resolution data, aggregated over 184 pesticides applied in the contiguous US, could result in an additional 53,606 range/pesticide pairs (28.3% of all such pairs) being deemed not likely to be adversely affected by the pesticide compared to CRD resolution data, with much of this benefit realizable using existing data in California.

The potential for high resolution data is therefore not a purely hypothetical inquiry. Current screening methods by the EPA are homogeneous across the contiguous U.S. (see SI Section 9) and thus do not take advantage of the existing high-resolution data offered by the California PUR system, which provides a true census of all agricultural (broadly defined) pesticide usage at the township scale (roughly 36 square miles) (3, Section 9 of the SI). Our results suggest that modifying screening methods to exploit the best available data (i.e., PUR data) is an immediate potential avenue for increased efficiency of pesticide evaluation and review. We previously showed that, for a single agent (carbaryl) and terrestrial plants endemic to CA, directly using such data could markedly increase the number of species potentially designated NLAA during the pesticide review process (3). This paper extends that work to 223 pesticides and 76 animal and 173 plant ranges endemic to California and suggests that using township usage data in the pesticide risk assessment process could result in 47% of range/pesticide pairings receiving an NLAA designation, compared to less than 5% when CRD usage data is used (Table S1, Fig 1).

For the contiguous US, we considered 184 pesticides and 577 animal and 451 plant ranges. Between plants and animals, California alone accounts for about 36% of the potential benefit of high-resolution usage data (i.e., township vs. CRD data) across the contiguous US, benefiting 10.5% of all pesticide/range pairs in the US. Extrapolating from CA data, we identified potential avenues for realizing similar gains elsewhere in the country. In total, 28.3% of all pesticide/range pairs are predicted to *benefit* from high resolution data (relative to CRD resolution alone); 18.2% benefit from data outside CA, and most of this is realizable from high resolution data in a minority of US states.

We identified 11 priority states in Southern Appalachia, the Southwest, Pacific Northwest, and Florida (Results; Fig. S55) that account for 68% of the remaining *potential* benefit. That is, of the *absolute* potential benefit of 18.2% of pesticide/range pairs designated NLAA for the entire contiguous US outside CA, an *absolute* benefit of 11.2% of all pairs is realized with these 11 states. According to our analysis, there is virtually no benefit to collecting high resolution data for most states in the Northern Great Plains, Midwest, and Northeast. Indeed, high resolution data for the 30 lowest priority states would collectively yield only about 10% of the potential benefits outside California (absolute benefit of about 2% of all pesticide/range pairs). It should be emphasized that taking full advantage of the high-resolution data already available in CA could be nearly as beneficial as collecting similar data in 11 other priority states.

Perhaps unsurprisingly, the states and regions where township resolution data are predicted to be of most efficacy are largely areas of high concentrations of endangered species, and in particular areas of high range-pesticide overlap. Simply taking the spatial product of range and pesticide density gives a good first-order approximation of where high-resolution data is likely to be of use (SI Section 3; Figs. S17-S20). However, our more rigorous attempt to identify priority states suggests less benefit through much of the Great Plains and Midwest than this more naÏve method implies; the distribution of benefit is also more concentrated in fewer states under our full analysis. It is also important to note that Hawaii was not included in our analysis, which has over 400 endangered species, many of which are endemic, and a wide variety of crops. Only low-resolution data for Hawaii is currently used by EPA methods (SI Section 9), but these basic numbers suggest that high resolution data could be of great value in Hawaii as well.

There are multiple barriers to collecting high resolution usage data. Currently federal regulations only require collection of pesticide records for restricted use pesticides (15). However, most pesticides in the US are registered for ‘general use’ and therefore not subject to record keeping requirements (16). Barriers cited to collection of pesticide use and usage records outside of California have been the costs for creating and managing a reporting system, the lack of existing data management infrastructure, and the absence of stakeholder support driven by privacy concerns on the part of landowners (17). Privacy concerns among stakeholders in agriculture, especially growers, are widespread in general. EPA data privacy regulations protect disclosure of personal data (18), although mistaken sharing of private information in the past has been cited as a cause of distrust in the agricultural community (19).

Recent developments include the announcement of an EPA OPP “workplan” (2) on endangered species assessment, which proposes multiple pilot projects to streamline and improve endangered species risk assessment, determine priority species, and improve engagement with stakeholders, including growers. Insufficient data is identified as a major barrier to efficient ESA assessment; EPA suggests working with pesticide applicators to provide sub-state resolution data on usage to increase the efficiency of ESA screening, and an expanded role for growers to provide information is identified as a priority.

Registrants, operators, state governments, farmers, and other stakeholders all benefit from the efficient, transparent, and accurate evaluation of pesticides undergoing registration and registration review. Our work suggests that, by increasing the efficiency of ESA evaluations, growers and other pesticide users have an incentive to provide sub-state, and ideally sub-county, usage information to EPA, especially in the priority states/regions identified. Major cotton producing states, for example, coincide strongly with the highest priority states identified (20).

Crop-specific usage data should therefore be a priority especially for cotton, vegetables, and field crops like potatoes and wheat that overlap with those states we have identified as benefiting from additional data collection.

While we have focused on the potential utility of pesticide use and usage data, the priority states we have identified should also be priority areas for increasing efforts to delineate endangered species’ ranges and critical habitats at high resolution. Such data would have multiple benefits and could facilitate conservation efforts not specific to a particular pesticide or crop.

In conclusion, our results suggest that nearly 50% of species-pesticide pairings could receive an NLAA designation if township usage data were available across the contiguous US. Furthermore, most of this benefit could be realized by fully taking advantage of the extremely high-resolution data already publicly available in California, and via geographically targeted data collection in a subset of states. By potentially avoiding over 50,000 Section 7 consultations across 184 major pesticides, resources could be focused on assessing the risk to the species/pesticide pairs that are more likely to need it. Thus, the value of collecting or releasing township level data could be tremendous in terms of expediting risk assessment and reducing potential harm to ESA listed species.

## Materials and Methods

### Base geographic data

Within California, township shapefiles obtained from the California Department of Pesticide Regulation Pesticide Use Reporting (DPR PUR) system (21) were aggregated to county and CRD scales for analysis. These geometries were linked to PUR usage data within California. Outside California, county shapefiles (500k:1 resolution) were obtained from the US Census (22), aggregated to CRD scale, and linked to US Geological Survey (USGS) National Pesticide Synthesis Project (NPSP) usage data. County membership in CRDs was obtained from the USDA (23). All geometries were re-projected to the USA Contiguous Albers Equal Area Conic USGS Version projection, to match USDA Cropland Data Layer raster data.

### Pesticide usage data

Pesticide usage estimates for the contiguous US outside California were obtained from the USGS NPSP (24-26). Outside California, this dataset is based on proprietary farm survey pesticide data reported at CRD scale. By-crop usage rates were estimated from USDA Census of Agriculture harvested crop acreage at CRD scale, and county-level estimates were then constructed using the National Land Cover Database 2011 (24,25). Both high and low usage estimates are provided, where the former extrapolates from nearby regions to fill in any missing data; we used high estimates throughout.

The California DPR PUR system directly tracks all pesticide usage in the state for a very broad range of “agricultural” end-uses. Residential and institutional uses are the only major categories omitted. Complete data from 2013 through 2017 was downloaded from the PUR data archives (27). Chemicals in the PUR system are classified by chemical code, more than one of which may map to a single active ingredient. For example, glyphosate is represented by eight chemical names (codes), e.g. pairs glyphosate (2997); glyphosate, diammonium salt (5810); glyphosate, dimethylamine salt (5972), etc. Therefore, to include all versions of an agent and for consistency across datasets, we (manually) mapped 483 unique pesticide names (all pesticides with non-zero usage between 2013 and 2017) from the USGS NPSP to 881 corresponding PUR chemical codes.

We separately processed usage data for 483 pesticides for years 2013 through 2017 to create individual data frames for each pesticide at either township, county, or CRD resolution (with PUR and NPSP datasets processed separately). Yearly usage datasets were filtered to maximum usage within each base polygon (e.g., township or county), and linked to the geographic shapefiles described above; these maximum usage maps were used for all subsequent usage/range overlap analyses. Finally, we classified each agent as either insecticide, herbicide, fungicide, or other, and considered the top 75 of the former three categories, by mass applied in 2017 nationwide (based on the NPSP dataset), for further analysis. We determined that two herbicides listed in the NPSP dataset, namely “metolachlor & metolachlor-s” and “dimethenamid & dimethenamid-p,” were redundant and excluded these from our final analysis and results. The geography of pesticide applications is presented in SI Section 2 (Figs. S12-S16).

### Land use data and species range land category fractions

The USDA Cropland Data Layer (CDL) provides satellite-based raster estimates of land use at 30-meter resolution across the contiguous US (48 states plus the District of Columbia). We downloaded this raster data for 2013 through 2017 from the USDA CropScape system (28). We followed the EPA Revised Method (Section 9 of SI) and remapped 134 unique CDL general land use classes to 13 agricultural classes, referred to as use data layer (UDL) categories. We additionally mapped 23 non-agricultural CDLs to seven “extended” UDL categories, namely developed, forest, barren, grassland or shrubland, water, wetlands, and miscellaneous land.

For each species range, the range polygon was used to mask the CDL and UDL rasters, for each year from 2013 through 2017. The fraction of the range mask that belonged to each category was determined, as was the minimum, mean, and maximum over the five-year period. The fraction of land that was agricultural in general was also recorded.

### Species range data

We used the USFWS Environmental Conservation Online System (ECOS) (29) to individually download shapefiles for all animal and plant ranges included in the ECOS, in May 2021. Range shapefiles were not available for all species in the ECOS. We obtained 708 animal ranges distributed among 625 unique species, and 949 plants ranges across 940 species. Ranges were tagged as being strictly contained within the continuous US, yielding 577 animal ranges for 501 species, and 454 plant ranges for 445 species. Inspection of ranges showed three terrestrial plant ranges (representing three unique species) in California to be likely erroneous, as they extended far into the Pacific Ocean, and these were excluded. Most species have only one associated range, and we use range as our base unit of analysis.

### Species range distributions

We determined the density of endangered species ranges across the county at different geographic resolutions. We iterated separately through each county, CRD, and state to determine the number of ranges that overlapped with each geographic entity. We also determined the number of ranges strictly contained within each county, etc. For ranges strictly contained within California, we also performed this procedure at township scale.

We repeated this procedure separately for plants and animals, and for major taxonomic groups— fishes, clams, reptiles, insects, amphibians, birds, snails, mammals, crustaceans, and arachnids; and flowering plants, ferns, lichens, and conifers—within each. This yields a set of species density maps that show where endangered species ranges are clustered at varying geographic resolutions and by major taxonomic group. Ranges are presented in depth in SI Section 1 and Figs. S1-S11.

### Overlap analysis

We performed a binary overlap analysis between each of the 223 pesticides considered and each species range, where we determined the fraction of the range that overlapped with *any* pesticide usage. We repeated this analysis at each usage resolution—township (in CA only), county, and CRD—using maximum usage over 2013-2017, as described previously. We performed the same analysis for all taxa, and did not distinguish between predator or prey species.

We identified 76 animal ranges and 173 plant ranges endemic to California (if the range extended into the Pacific Ocean but not the boundaries of any other state, it was still included), and performed this overlap analysis for township, county, and CRD usage resolutions. For all ranges, the analysis was also performed at national scale under county and CRD resolution; usage data derived from the PUR was used within California, while data from the USGS NPSP was used elsewhere.

In addition to overlap fraction, we also recorded the minimum, mean, and maximum pesticide exposure in kg m^-2^ at each resolution, for each range. For example, suppose a county was treated with 100 kg of glyphosate, and 30% of this county overlapped with a species range. For minimum exposure, it is possible no application occurred in the overlapping area, and minimum exposure is therefore 0 kg. For the mean, we suppose a uniform distribution of application and expect 30 kg in the overlapping area, while for the max, we assumed that all 100 kg were applied in the overlap. We sum these exposures across all counties, and then divide by the range area to get possible exposure intensities.

We generally divided ranges into those with <1% usage overlap, which could potentially be considered not likely to be adversely affected and excluded from further regulatory analysis, and those with >1% usage overlap. We adopt the EPA terminology and refer to the former as not likely to be adversely affected (NLAA), while the latter are considered likely to adversely affect (LAA). Detailed overlap results within CA are provided in SI Sections 4 and 5 (Figs. S21-S32).

### Regression and classification

We performed logistic regression to see if characteristics of species’ ranges or pesticide exposures at county scale predict whether township-scale usage data is likely to be of benefit in classifying a species range as NLAA. That is, for each pesticide and range, we determined if a range considered LAA (>1% range/usage overlap) under county usage data switched to NLAA under township data. If so, township-level data was beneficial.

As predictors, we used range area, mean agricultural fraction (mean fraction of each range that is agricultural land over 2013 through 2017), mean pesticide exposure (kg m^-2^) at county resolution, or all three. We then determined the optimal probability cutoff for classifying ranges as excluded or non-excluded using an ROC curve and chose the point on the ROC curve closest to the upper left. Note that we also examined other land use classifications and species taxa as predictors, but these were not generally useful. A classifier was constructed for every pesticide/range pair in CA, and results were aggregated separately for plants and animals.

Models were generated for plants and animals separately, for each individual pesticide, and for each predictor alone or all in combination. The area under the curve (AUC) of the ROC curve was our primary metric for model performance. Comparative model performance was evaluated across all pesticides by pooling AUC values for each model specification (set of predictors) and determining the bootstrap distribution of the mean AUC. We determined if a given set of predictors performed better than chance in classifying ranges used a bootstrap re-sampling procedure, wherein observations were repeatedly re-sampled with replacement, yielding a new set of AUC values. If 95% of the bootstrap AUC values were greater than 0.5, the classifier was considered better than chance.

To compare models, a similar bootstrap procedure was performed on the difference between AUC values for the two models. We determined the aggregate performance of a model across species ranges and pesticides by finding the AUC for each pesticide, and again performing bootstrap resampling to yield a bootstrap AUC distribution. Models with all three predictors were superior to other possible specifications for modeling animal range overlaps but did not outperform a model using range area alone as a predictor for plant range overlaps (Figs. S33-S36; SI Section 6). However, for consistency, we used all predictors for both plants and animals for extrapolating to the contiguous US. A detailed breakdown of sensitivities, specificities, and AUCs for all models considered is given in Table S2.

### Prediction of priority areas for high resolution data

We used the results of the classifiers trained on California data to predict where, across the contiguous US, improving spatial usage resolution to township resolution is most likely to be of benefit. We trained the models in California, and then applied them nationwide, for both plants and animals, and for 223 pesticides, if possible. Since not all pesticides used nationally had any use footprint in CA, it was not always possible to perform a prediction. Ranges that were predicted to switch from LAA to NLAA status were tagged, and density maps at county and CRD scale, similar to those for species ranges (described previously), were generated. This was performed for each individual pesticide, with results also aggregated across major taxa (animals vs. plants) and pesticide class (insecticide, herbicide, fungicide). Summary results are provided in SI Section 7 and Figs. S37-S50.

### Identification of priority areas for high resolution data

We used an iterative approach to identify states where the benefit of collecting high resolution data was greatest. Since species’ ranges often cross state lines, rigorously identifying an optimal subset of N states requires evaluating (48 choose N) combinations, or 6.54 billion combinations if N = 10. Therefore, we used the following iterative procedures to approximate the optimal combinations. For each state, we determined, *compared to a baseline where only CRD usage data is available*: 1) The number of additional ranges predicted to switch to NLAA if township-level data was available for that state alone, plus CA, given that CA data is already available. 2) The number of additional ranges predicted to switch to NLAA if this was the only state for which township usage data was not available. This “leave-one-out” analysis identifies states that contain many ranges that overlap with other states, and that would benefit from township data.

Between the two method’s top ten states, fifteen states were identified as important with respect to the impact of township resolution data for both plants and animals (AL, AZ, CO, FL, GA, ID, MS, NC, NV, OR, TN, TX, UT, VA, and WA; see Fig 4 and Figs S51-S55). By then applying the leave-one-out method to this 15-state subgroup, and combining results across plants and animals, we arrived at a final group of 11 priority states. Additional details are provided in SI Section 8.

### Python code and processed data

All data manipulation and analyses were performed in Python. We provide processed datasets (pesticide usage datasets and animal and plant range shapefiles) and Python code that replicates all figures in the paper as Supplementary Materials.

## Acknowledgments

We thank L. Duzy, A. Frank-Reeves, Y.W. Li, and P. Ashfield for discussions that helped inform this work. Bayer CropScience, Award:FP00016353 provided funding.

