## Supplementary Information Appendix for "Identifying opportunities for high resolution pesticide usage data to improve the efficiency of endangered species pesticide risk assessment"

### 1. Characterization of plant and animal ranges

#### Ranges restricted to California only

We initially identified 76 animal ranges and 176 plant ranges with range geometries restricted solely to California. However, three terrestrial plant ranges (*Arctostaphylos hookeri var. ravenii, Clarkia franciscana,* and *Erysimum menziesii*) that extended into the Pacific Ocean were considered erroneous, and were excluded from the analysis, leaving 173 plant ranges. Figs. S1 and S2 give maps of the range polygons and range density at township scale, respectively. Figs. S3 and S4 depict range densities disaggregated by major taxa for animals and plants, respectively.

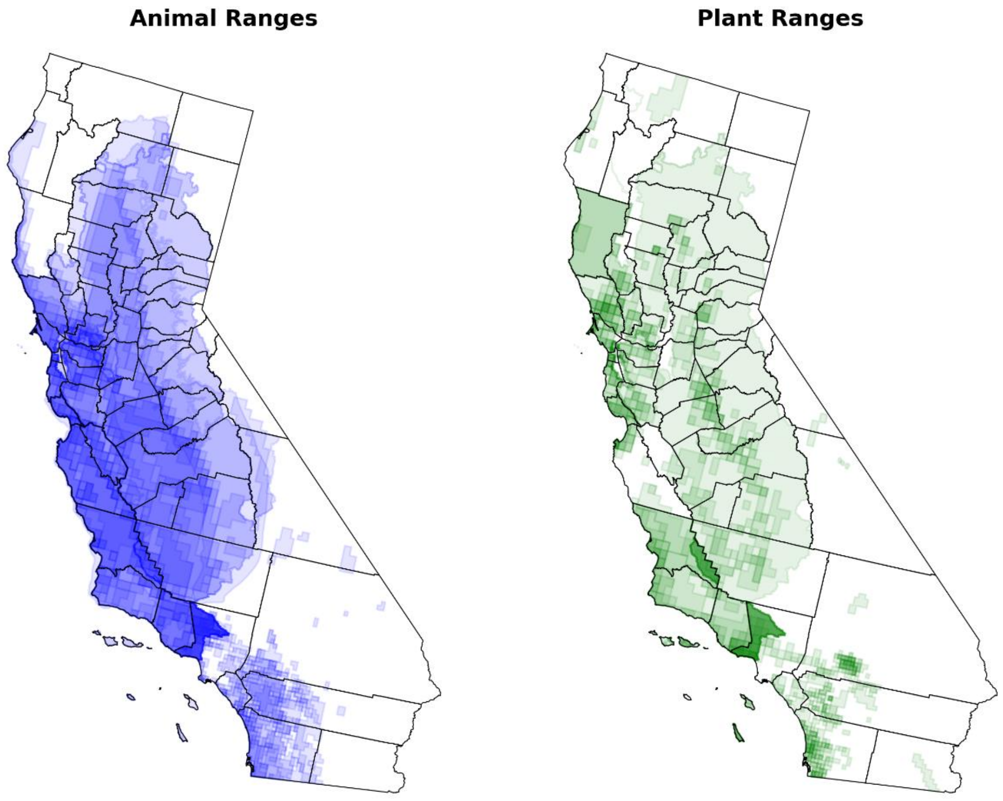

**Fig S1.** Animal (left) and plant (right) range geometries restricted to the state of California.

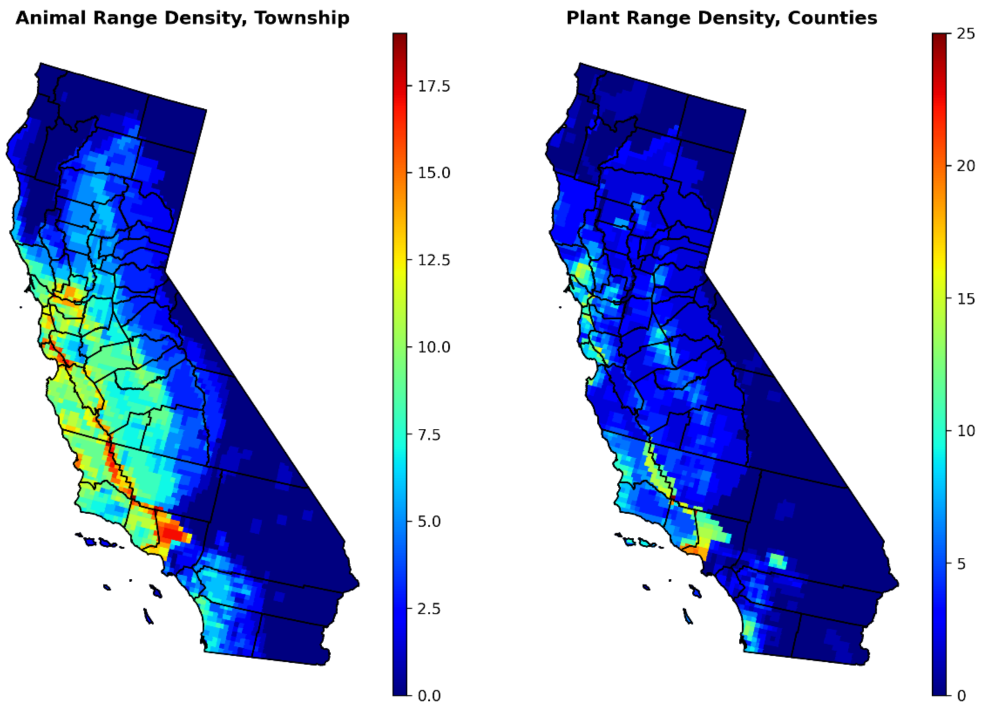

**Fig S2.** Animal and plant range densities at township resolution, for ranges contained solely within California. Colors represent the number of ranges that overlap, at least in part, the underlying township. County outlines are also given in black for reference.

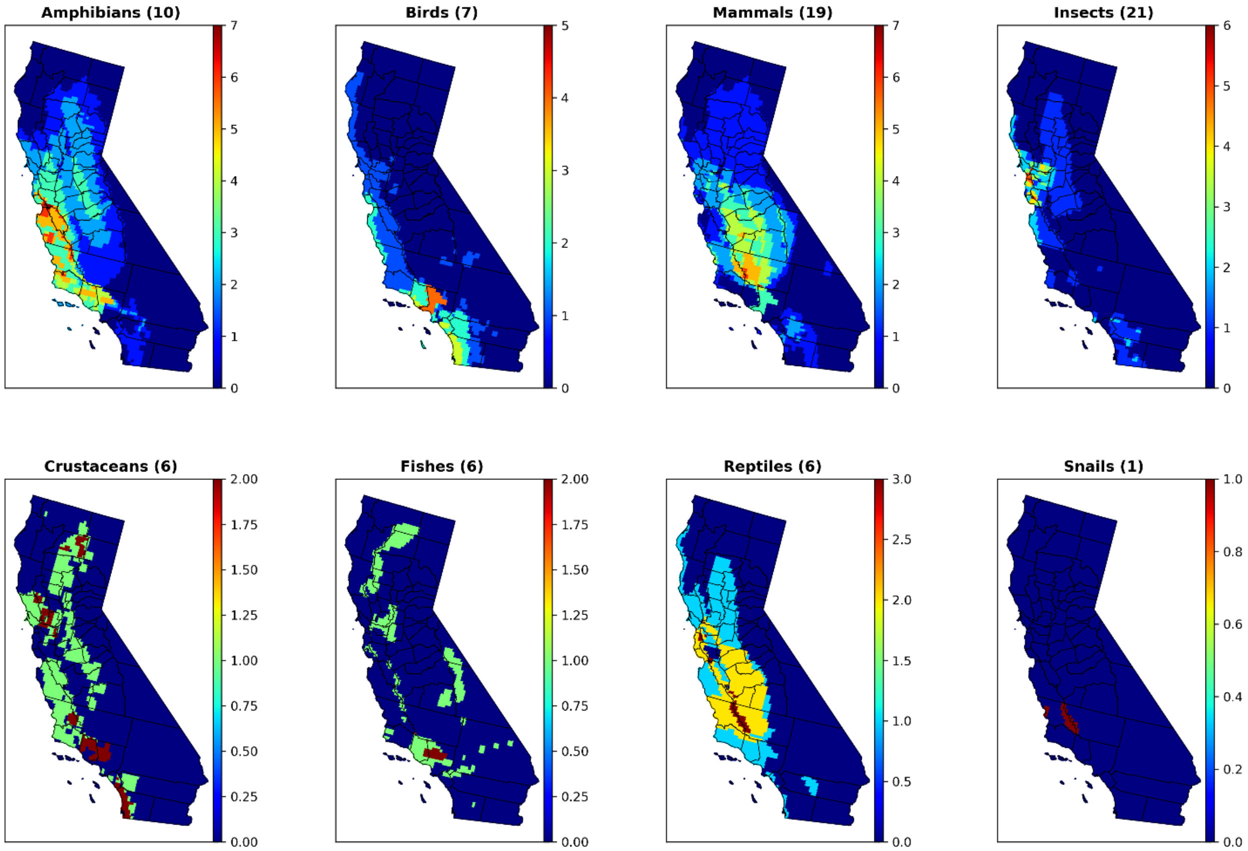

**Fig S3.** Animal range densities by taxon in California. Numbers in parentheses indicate the total number of ranges belonging to each taxon.

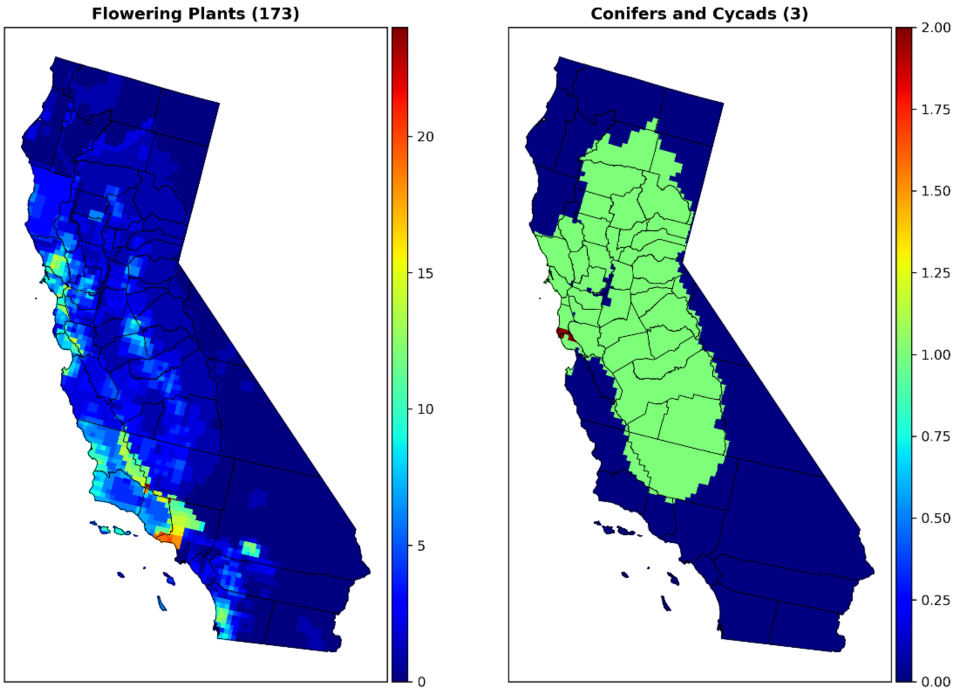

**Fig. S4.** Plant range densities by taxon in California. Numbers in parentheses indicate the total number of ranges belonging to each taxon.

**Ranges restricted to contiguous United States**

We identified 577 animal ranges and 454 endangered plant ranges located within the contiguous United States; as for California, three plant ranges were excluded from further analysis. The raw range geometries, as well as range density at county scale are depicted in Figs. S5 and S6, while Figs. S7 and S8 show range densities disaggregated by taxa at county resolution. State-level range distributions are characterized in Figs. S9-S11.

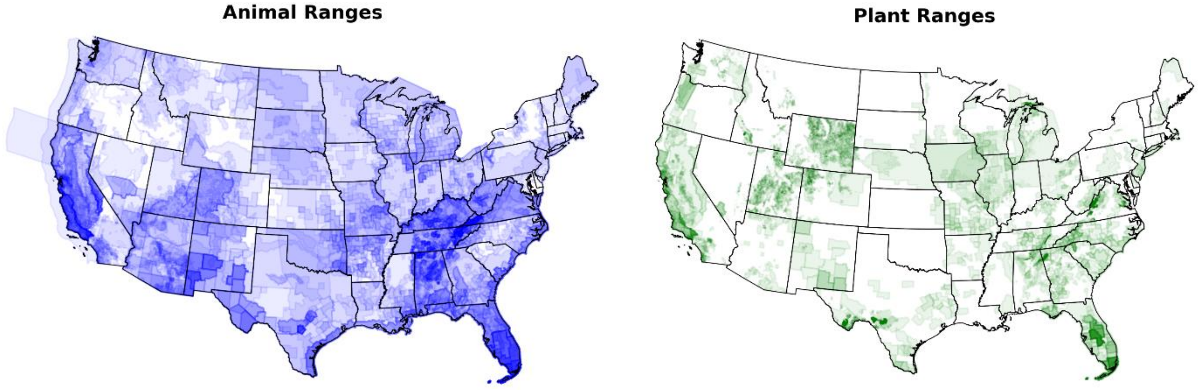

**Fig S5.** 577 animal (left) and 451 plant (right) range geometries restricted to the lower 48 states.

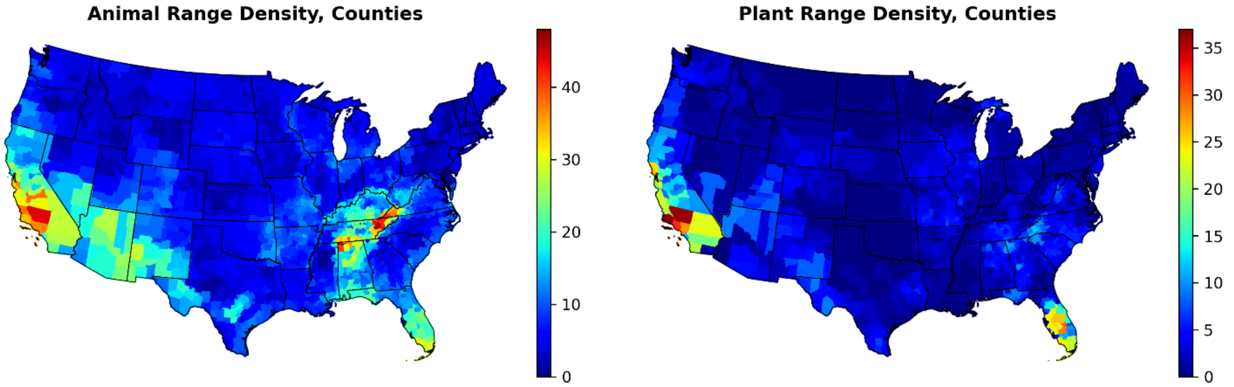

**Fig. S6.** Endangered animal (left) and plant (right) ranges as a density map at county scale. The color represents the number of ranges that overlap, at least in part, a given county.

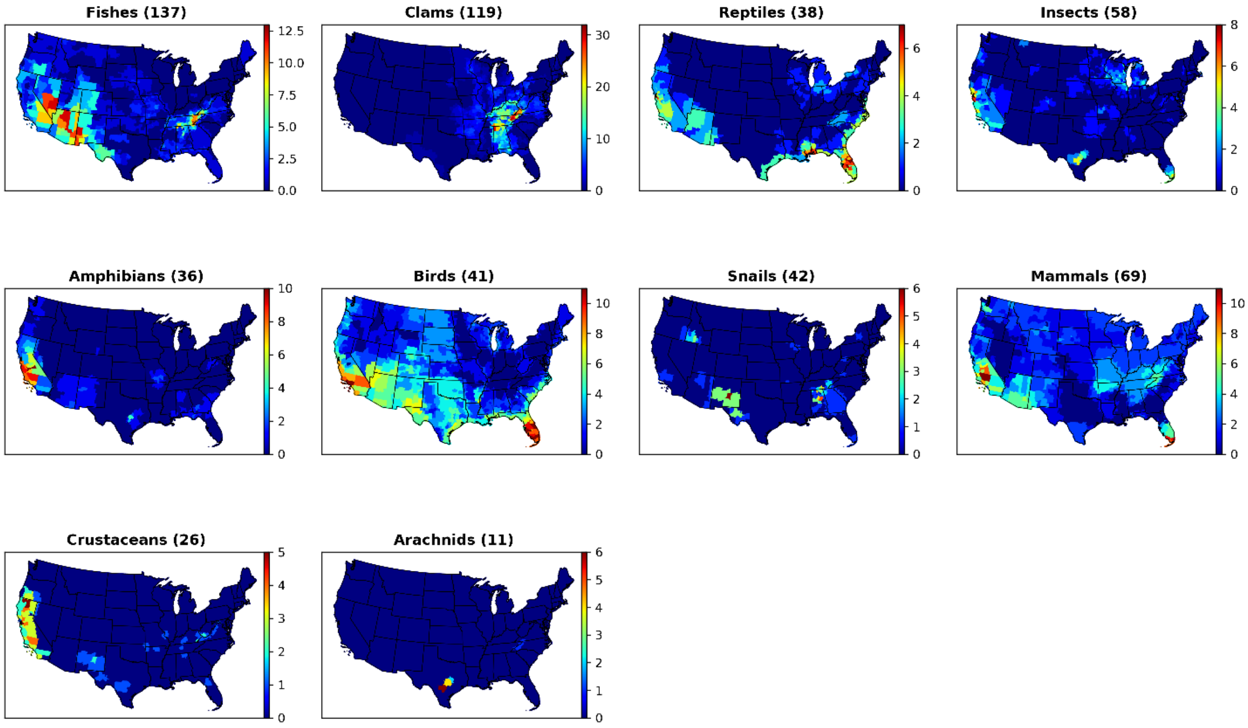

**Fig. S7.** Endangered animal range density, at county scale, by taxa. The total number of ranges for each taxon is given in parentheses in its respective subplot title.

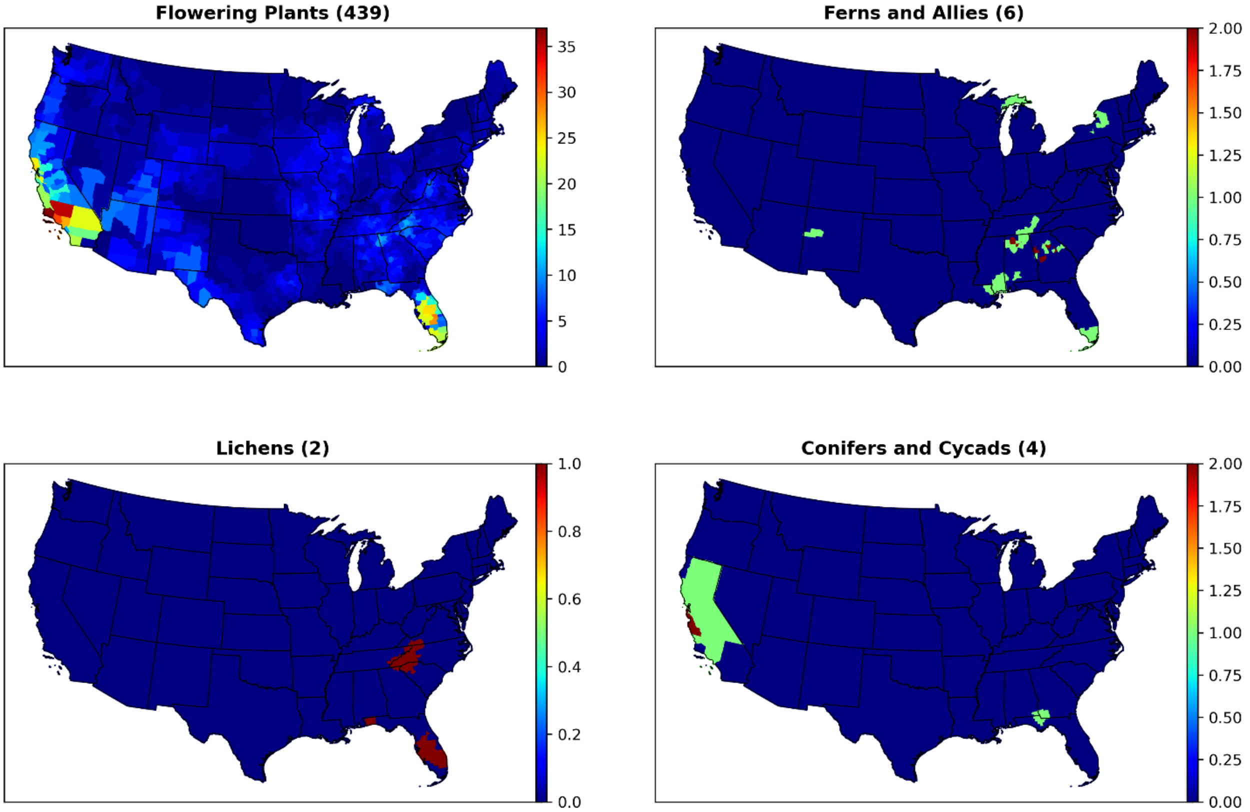

**Fig. S8.** Endangered plant range densities by taxa at county scale. The total number of ranges for each taxon is given in parentheses in the subplot titles.

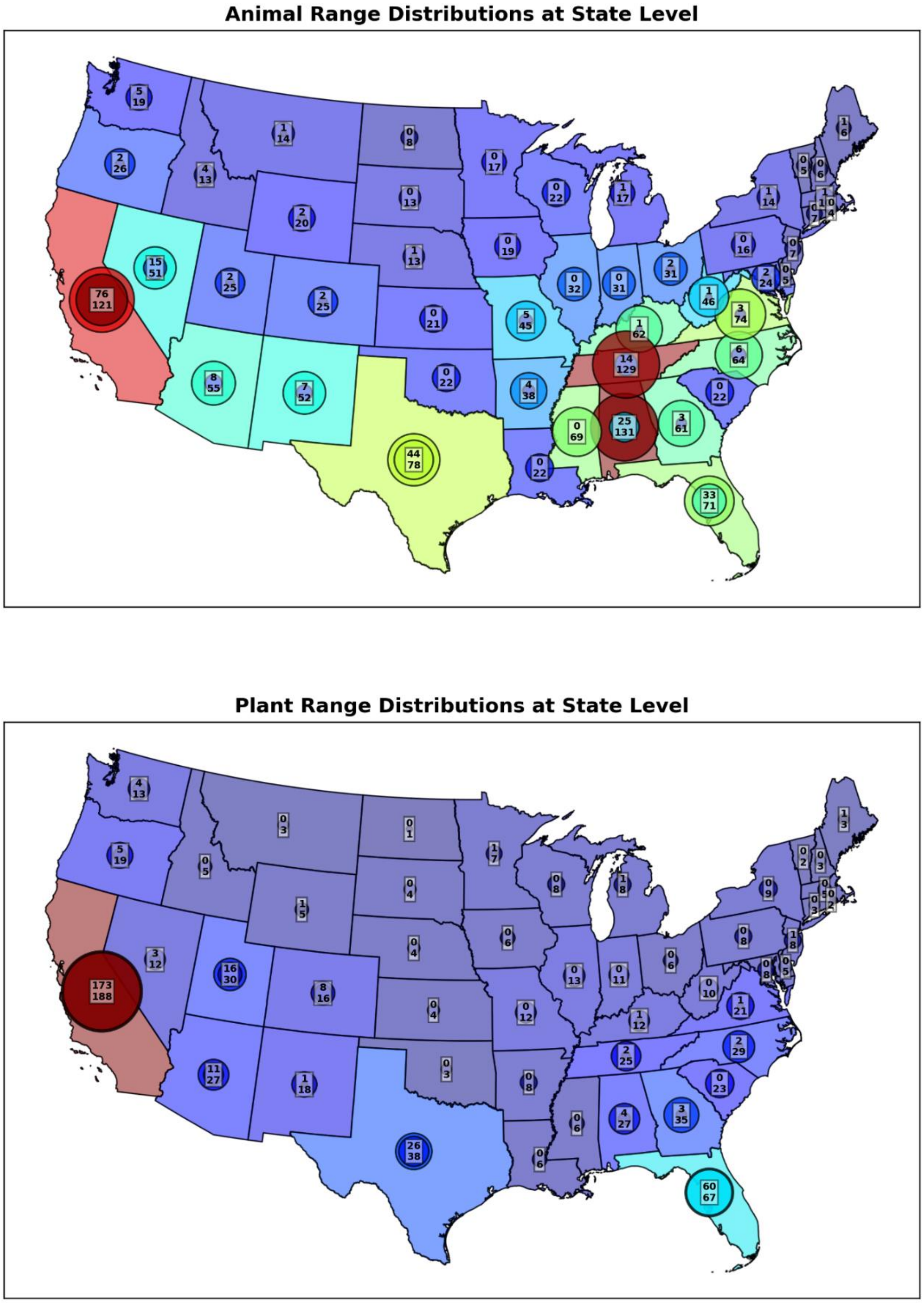

**Fig. S9.** State-level characterization of endangered animal (top panel) and plant (bottom panel) range densities. The upper number within each circle gives the number of range endemic to each state, while the lower number shows how many species ranges partially overlap with a given state. The inner circles then scale with the number of ranges strictly contained within a given state, while the outer circles scale with the number of ranges that at least partially overlap with a state. Animal ranges have a more cosmopolitan distribution and are more likely to cover multiple states. Plants are usually restricted to a single state, with about 58% of ranges contained solely within either California, Texas, or Florida.

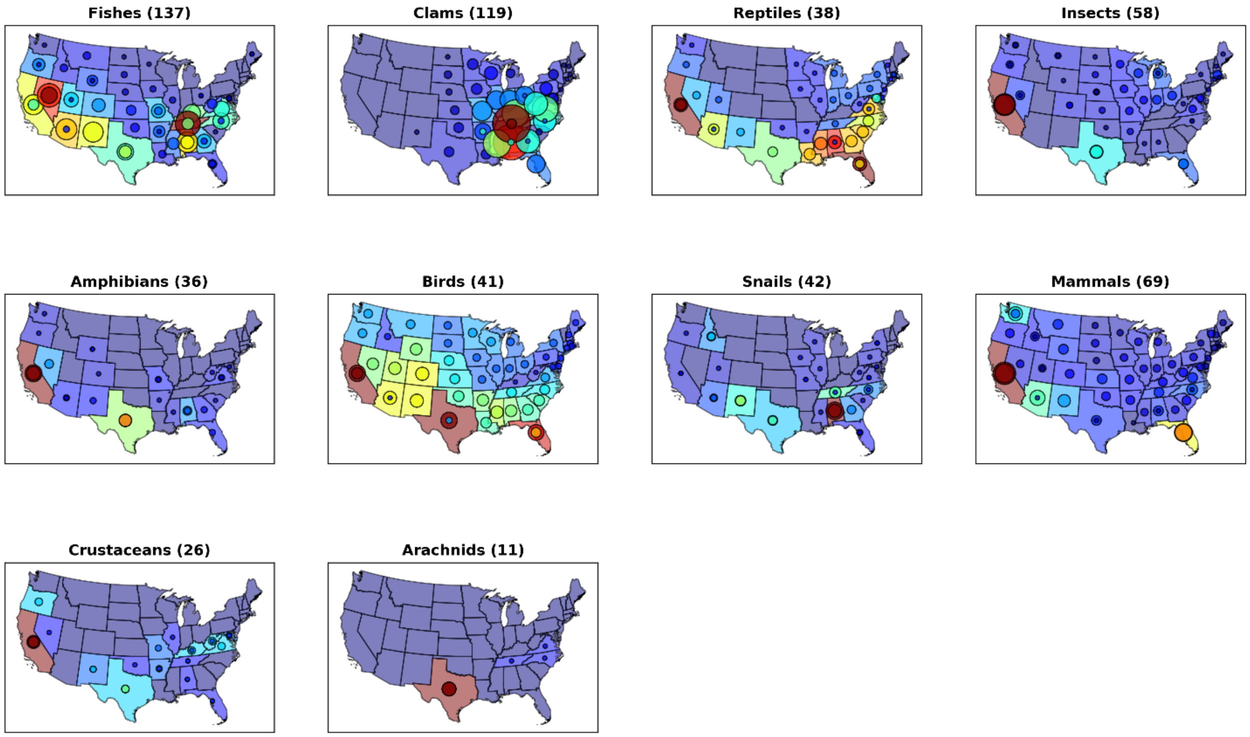

**Fig. 10.** State-level characterization of animal range densities, disaggregated by taxa. As with Fig S9, the inner circles represent the number of ranges contained completely within a state, while the outer circles scale with the number of ranges that overlap with the state at all.

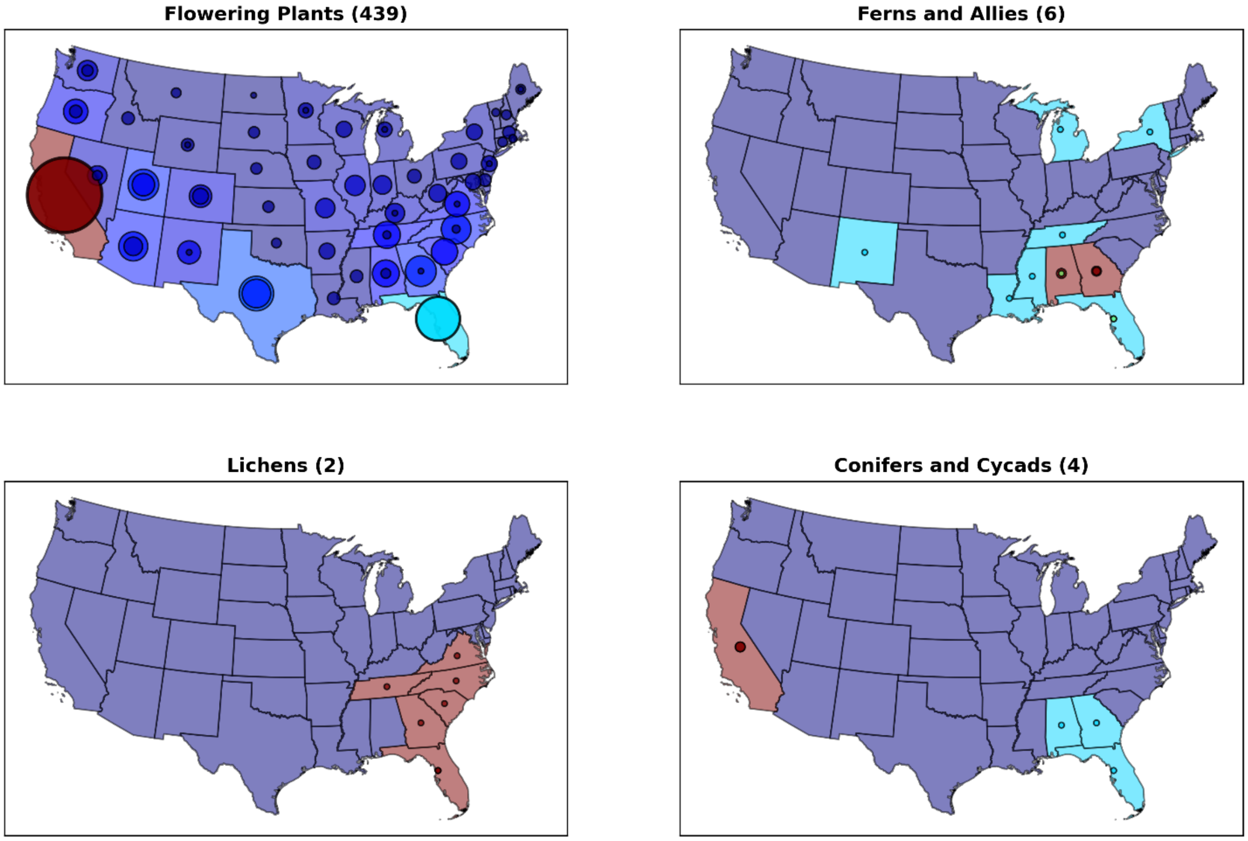

**Fig S11.** State-level characterization of plant range densities, disaggregated by taxa. As with Fig S9, the inner circles represent the number of ranges contained completely within a state, while the outer circles scale with the number of ranges that are at least partially contained within the state.

### 2. Overall geography of pesticide applications

County-scale maps of maximum pesticide mass over 2013-2017 applied in the contiguous US (in terms of kg applied per acre), as well as the absolute number of individual pesticides applied, divided by major class, are given in Fig. S12. Results across all compounds are aggregated in Fig S13. Similar maps of township-scale pesticide applications within California are given in Figs. S14 and S15.

Maps for pesticide application at all available years (1992 through 2017), the maximum over 2013-2017, and at township (within CA only and only for 2013-2017), county, CRD, and state scales can be visualized using a simple iPython/jupyter notebook app available at the link in Eikenberry et al. (2022). The app provides maps of pesticide usage mass on either a continuous scale, or in binary terms (applied or not applied). A screenshot is given in Fig. S16.

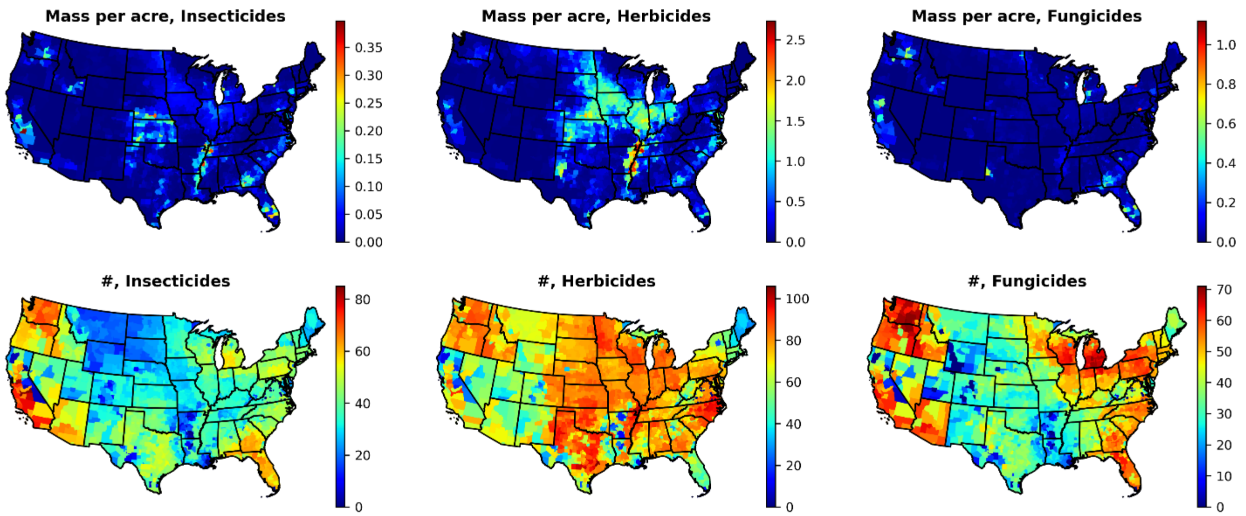

**Fig. S12.** The top three panels give total pesticide mass (in kg) applied per acre across the contiguous US, divided into insecticides, herbicides, and fungicides. Shown is the sum of the county-wise maximums over 2013-2017 for all compounds. The bottom panels give the number of individual compounds applied in any given county, disaggregated by major pesticide class. Again, the county-wise maximum is summed for each county.

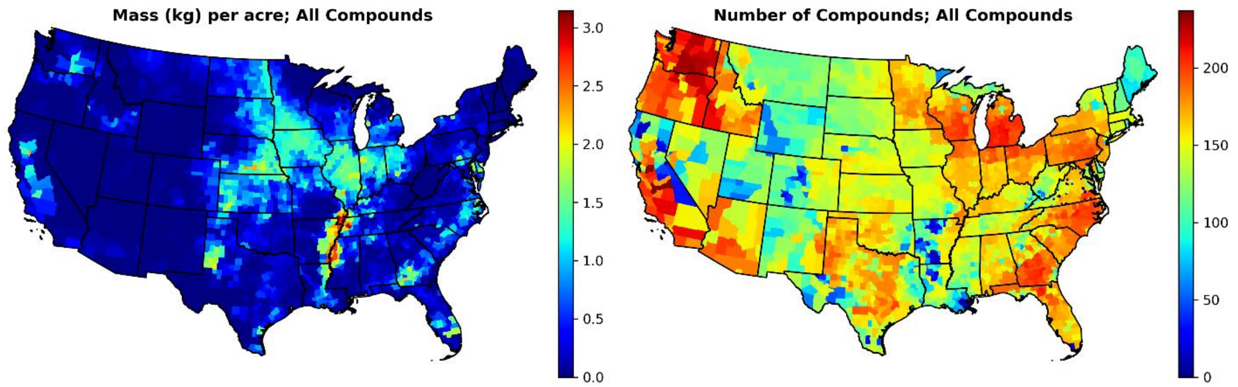

**Fig. S13.** Mass (kg per acre) and total number of compounds applied in each county, summed across all 223 compounds considered in this work. Note that the county-wise maximum across 2013-2017 is used for each pesticide.

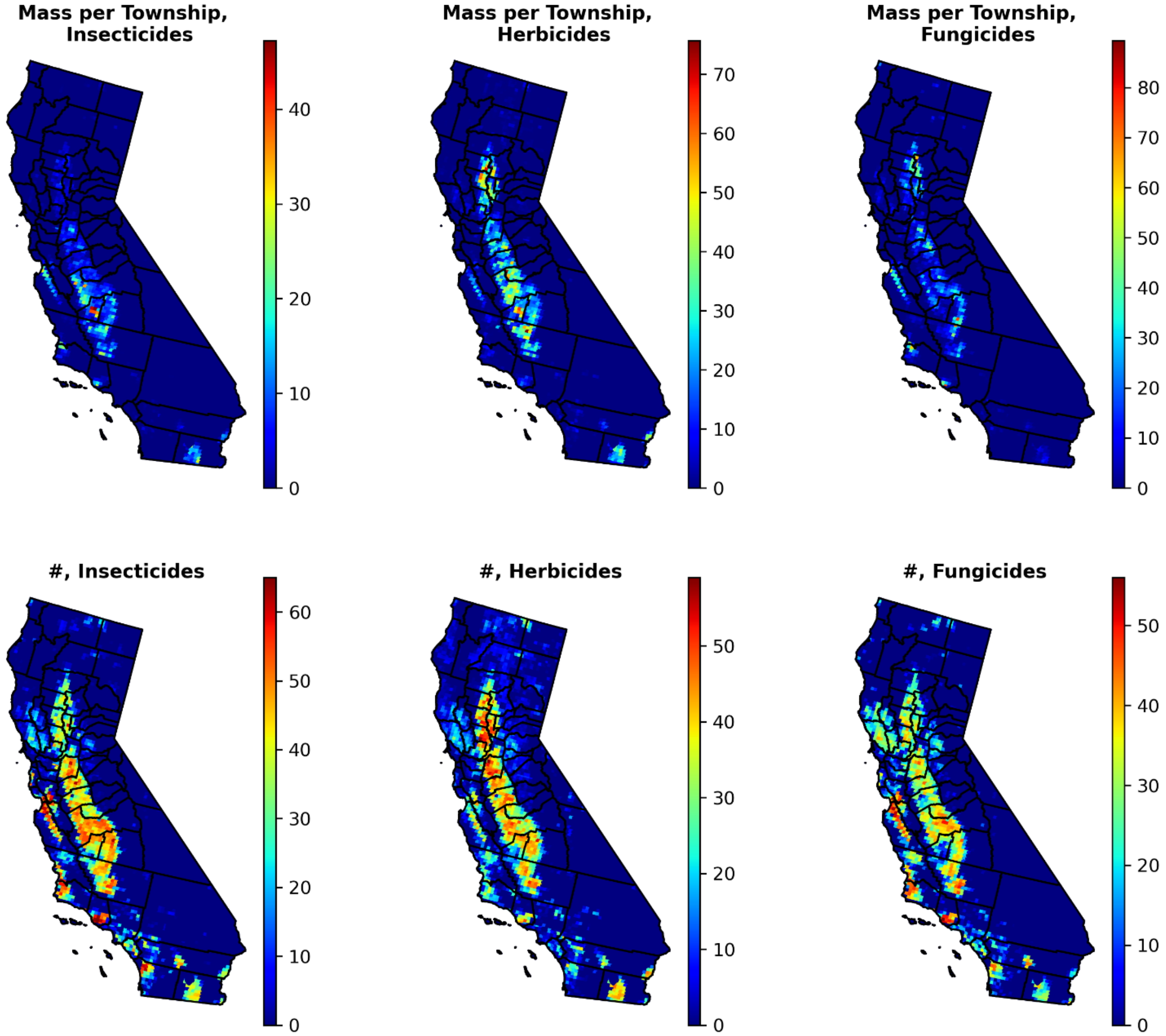

**Fig. S14.** The top three panels give total pesticide mass (in kg) applied per township in California only, divided into insecticides, herbicides, and fungicides. Shown is the sum of the township-wise maximums over 2013-2017 for all compounds. The bottom panels give the number of individual compounds applied in any given township, disaggregated by major pesticide class. The township-wise maximum is summed for each township.

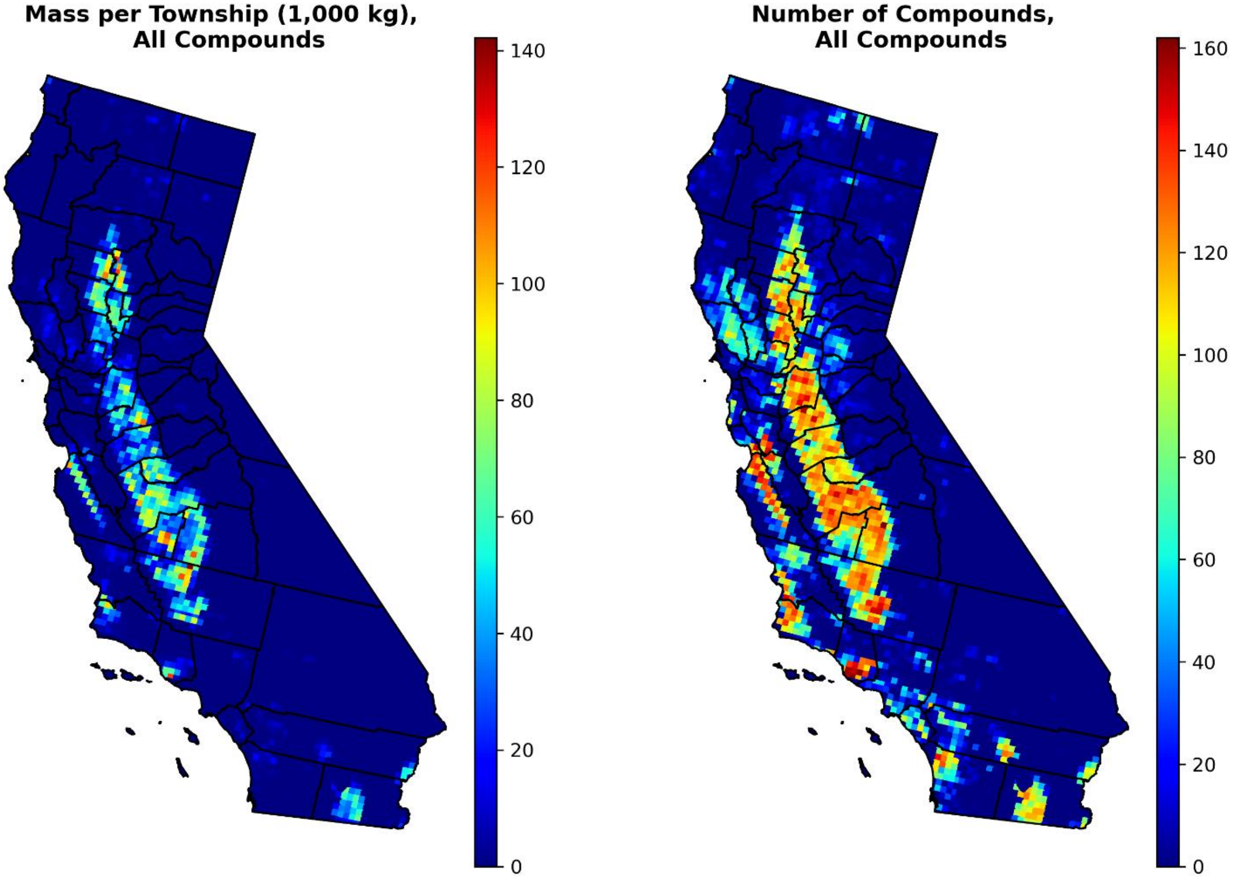

**Fig. S15.** Mass (kg per township, left) and total number of compounds (right) applied in each township in CA only, summed across all 223 compounds considered in this work. The township-wise maximum across 2013-2017 is used for each pesticide.

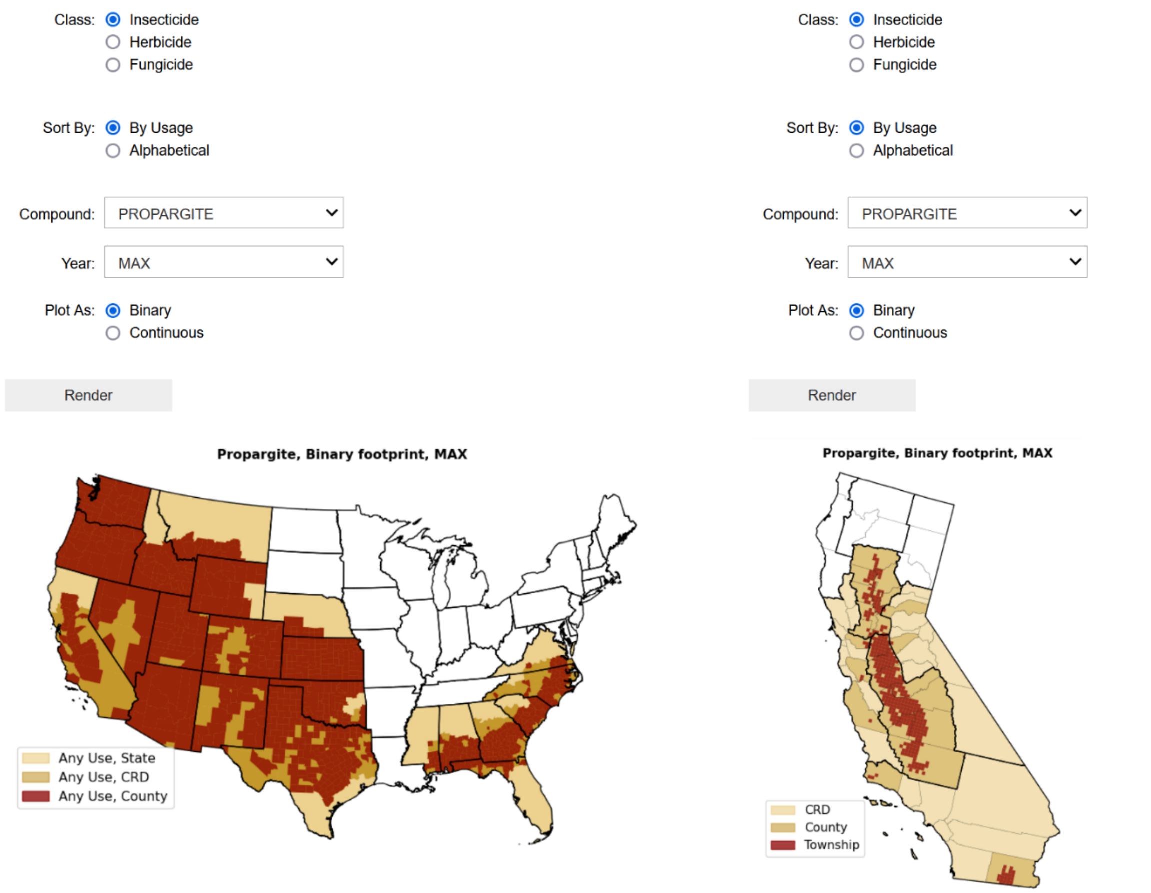

**Fig. S16.** Screenshots of Jupyter notebook apps to visualize usage footprints for various pesticides either across the contiguous US (left panels) or California only (right panels; screenshots have been cropped for clarity). Binary footprint refers to the binary presence of a pesticide in a given geographic entity (state, CRD, county, or township) in a chosen year. Year MAX is the geometry-wise maximum from 2013-2017.

### 3. Range/Compound pair density maps

We take the product of the number of pesticides applied in a base geometry (e.g., county) with the number of ranges (either animal, plant, or both). This product represents the number of range/compound pairs present in each polygon. The resulting maps are a first-order approximation for where high resolution data on pesticide could be most useful: The more range/compound pairs, the more opportunities for data to be of use, while areas with virtually no such pairs are unlikely to be important. It is also potentially instructive to examine disagreement between our primary results and those of the results of this simple method.

The process for constructing county-level maps is illustrated, along with the results, in Fig. S17. Maps with insecticides, herbicides, and fungicides considered separately are also given in Fig. S18. State-level maps are given in Figs. S19 and S20.

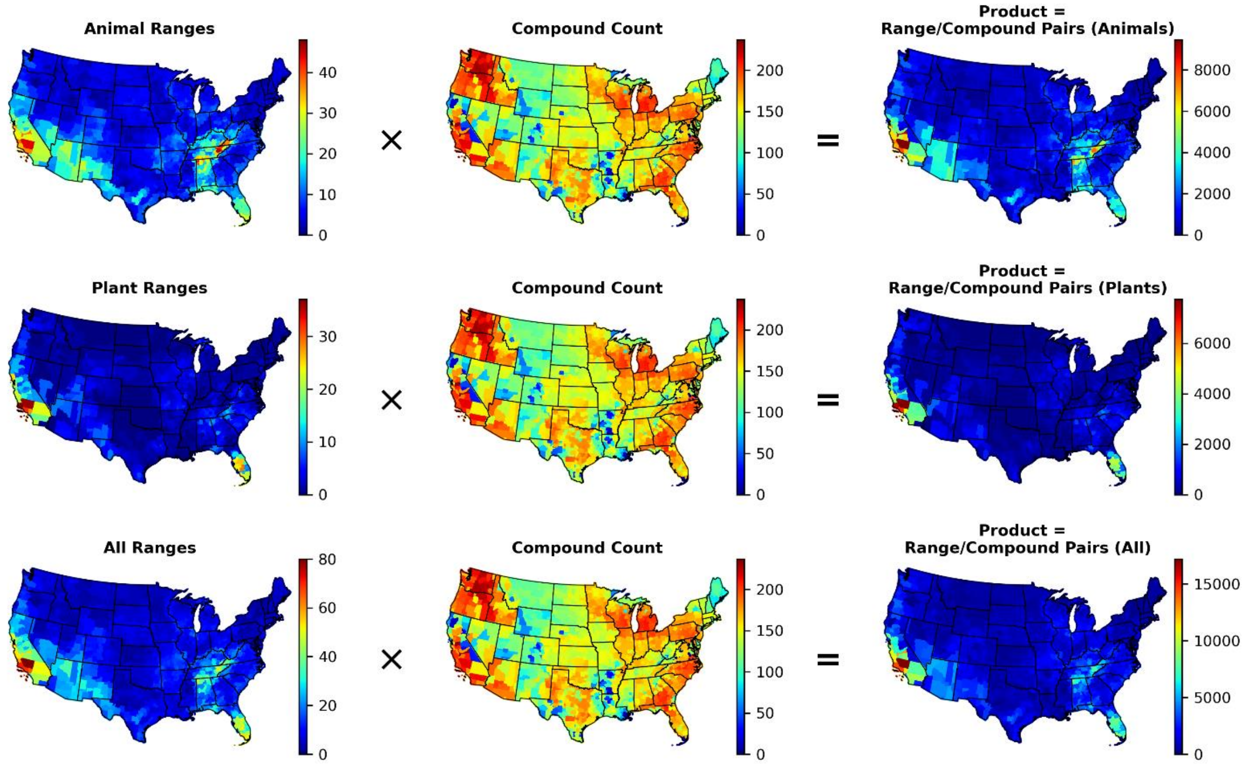

**Fig. S17.** County-wise product of the number of ranges present in each county and the total number of compounds applied in that county (maximum over 2013-2017). From top to bottom, the basic calculation and results (right-most maps) are illustrated for animals, plants, and animals and plants combined.

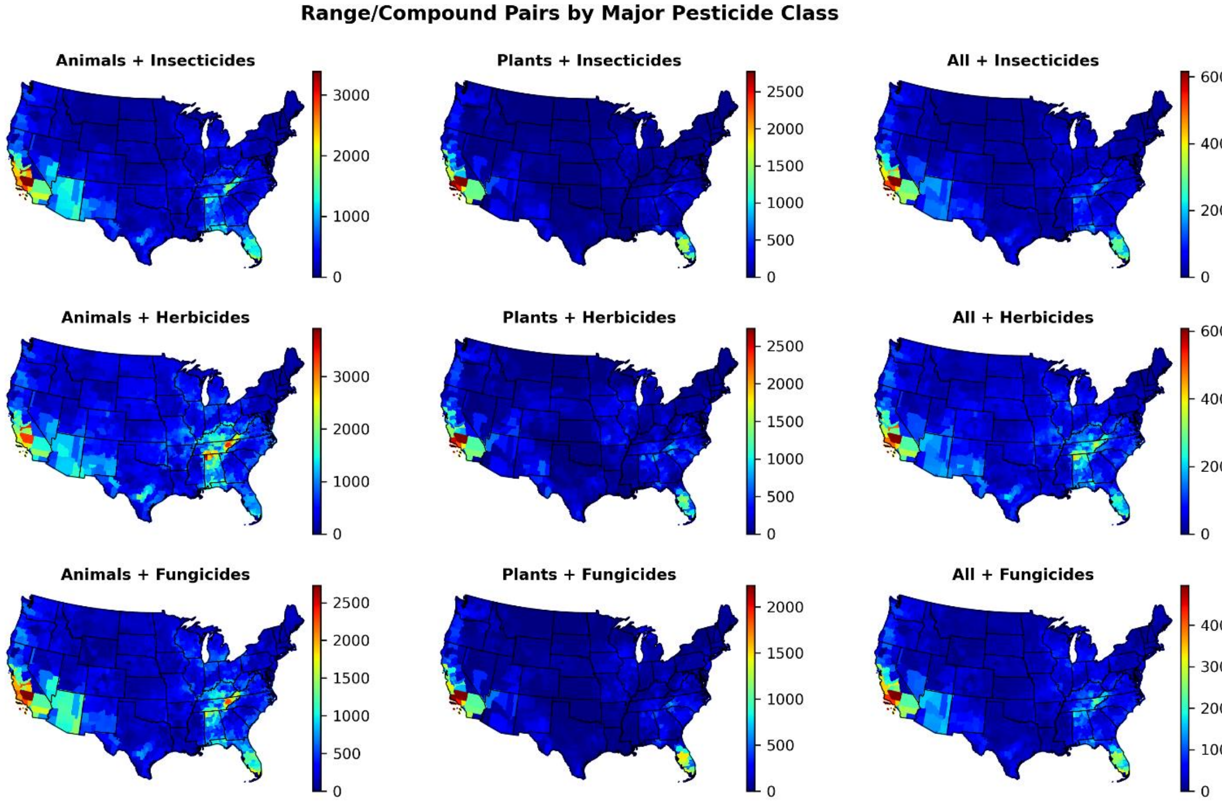

**Fig. S18.** County-wise product of the number of ranges present in each county and the total number of compounds applied in that county (maximum over 2013-2017), for all nine combinations of animals, plants, and animals + plants with insecticides, herbicides, and fungicides.

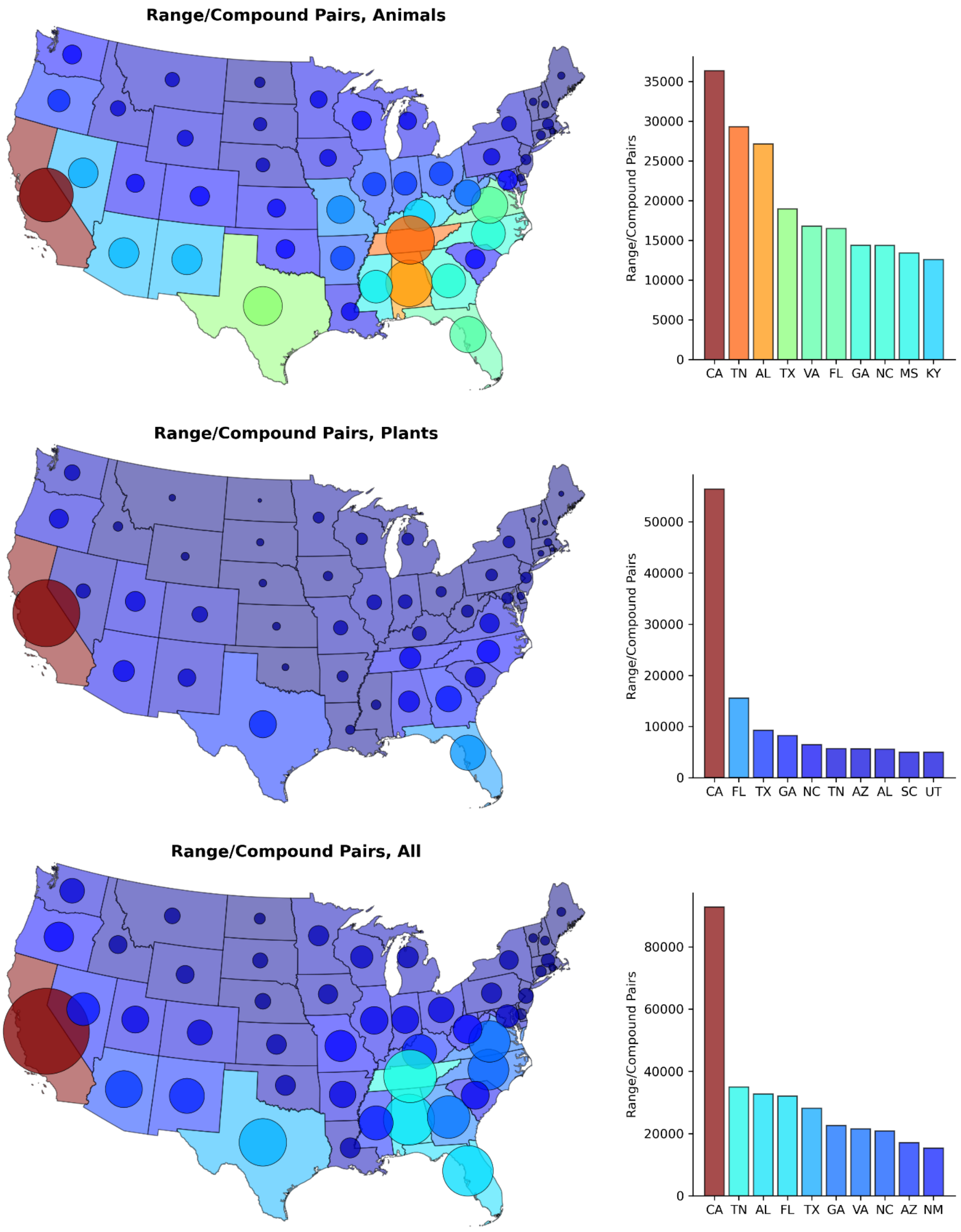

**Fig S19.** The maps on the left illustrate the relative number of range/compound pairs in each state, with the bubble area scaling in proportion to this metric. The bar graphs on the right show the number of range/compound pairs in each state for the top 10 states. From top to bottom, results are given for animals, plants, and animals + plants, in combination with all compounds.

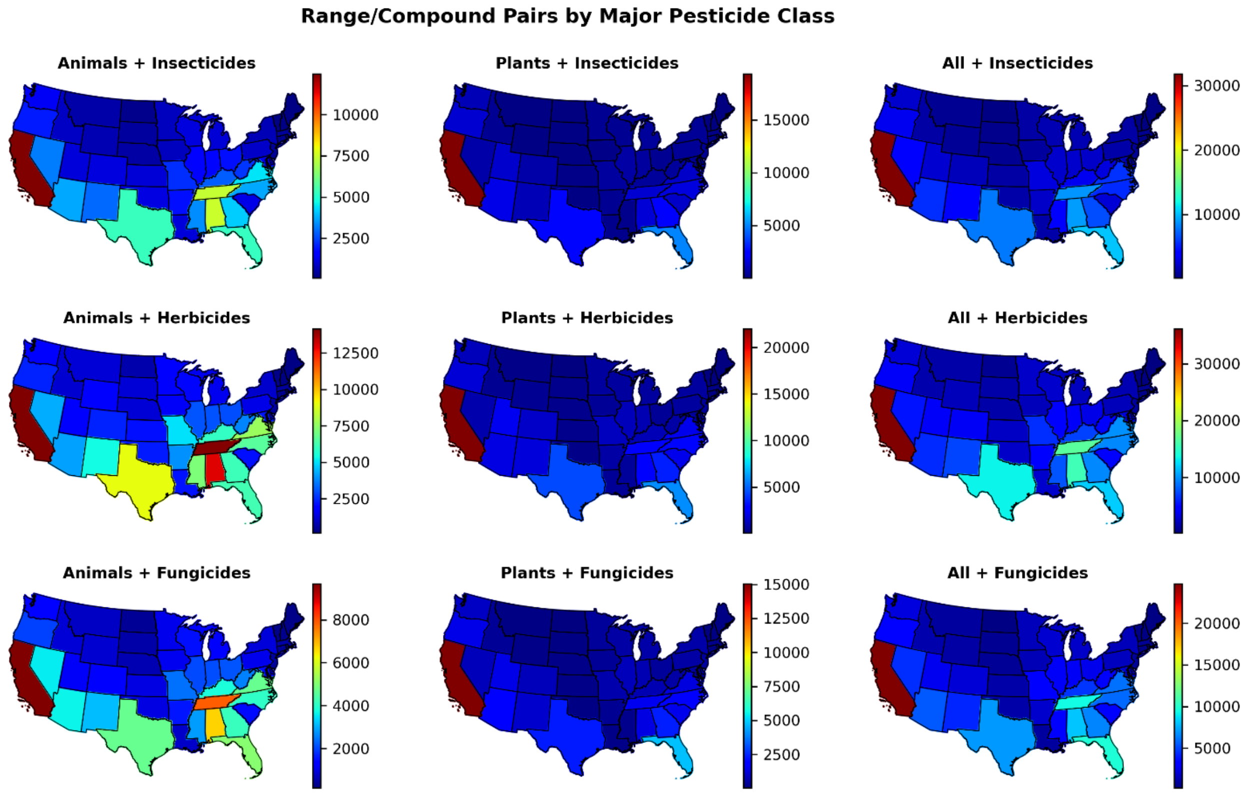

**Fig. S20.** State-wise product of the number of ranges present in each state and the total number of compounds applied in that state (maximum over 2013-2017), for all nine combinations of animals, plants, and animals + plants with insecticides, herbicides, and fungicides.

### 4. Usage resolution and NLAA results in California

The distributions of the number of animal and plant ranges, disaggregated by compound class, that can be designated NLAA (i.e., <1% range/compound overlap) for either CRD, county, or township resolution usage data, are given in Figs. S21 and S22. Figs. S23 and S24 are similar but illustrate the *changes* in how many ranges are considered NLAA under a shift in resolution. We also illustrate the fraction of ranges found NLAA, for the three usage resolutions, as a function of overall land area treated (calculated using the township-level usage footprint), in Fig. S25.

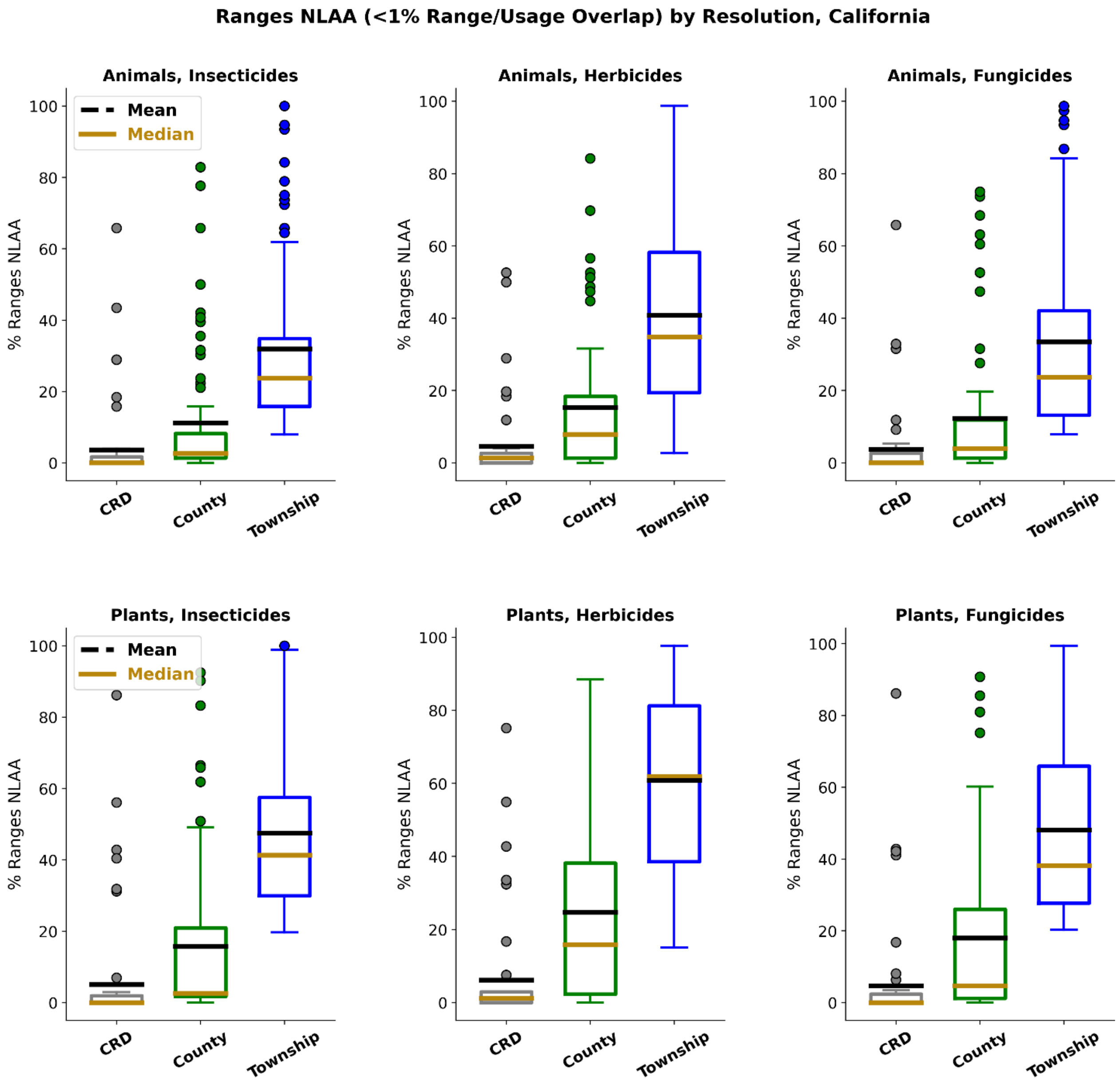

**Fig. 21.** Distribution of ranges designated NLAA in California, under either CRD, county, or township usage resolution across 223 compounds, visualized as boxplots. Distributions are disaggregated by pesticide class (insecticide, herbicide, fungicide, from left to right), with results for animals given in the top panels and results for plants in the bottom panels.

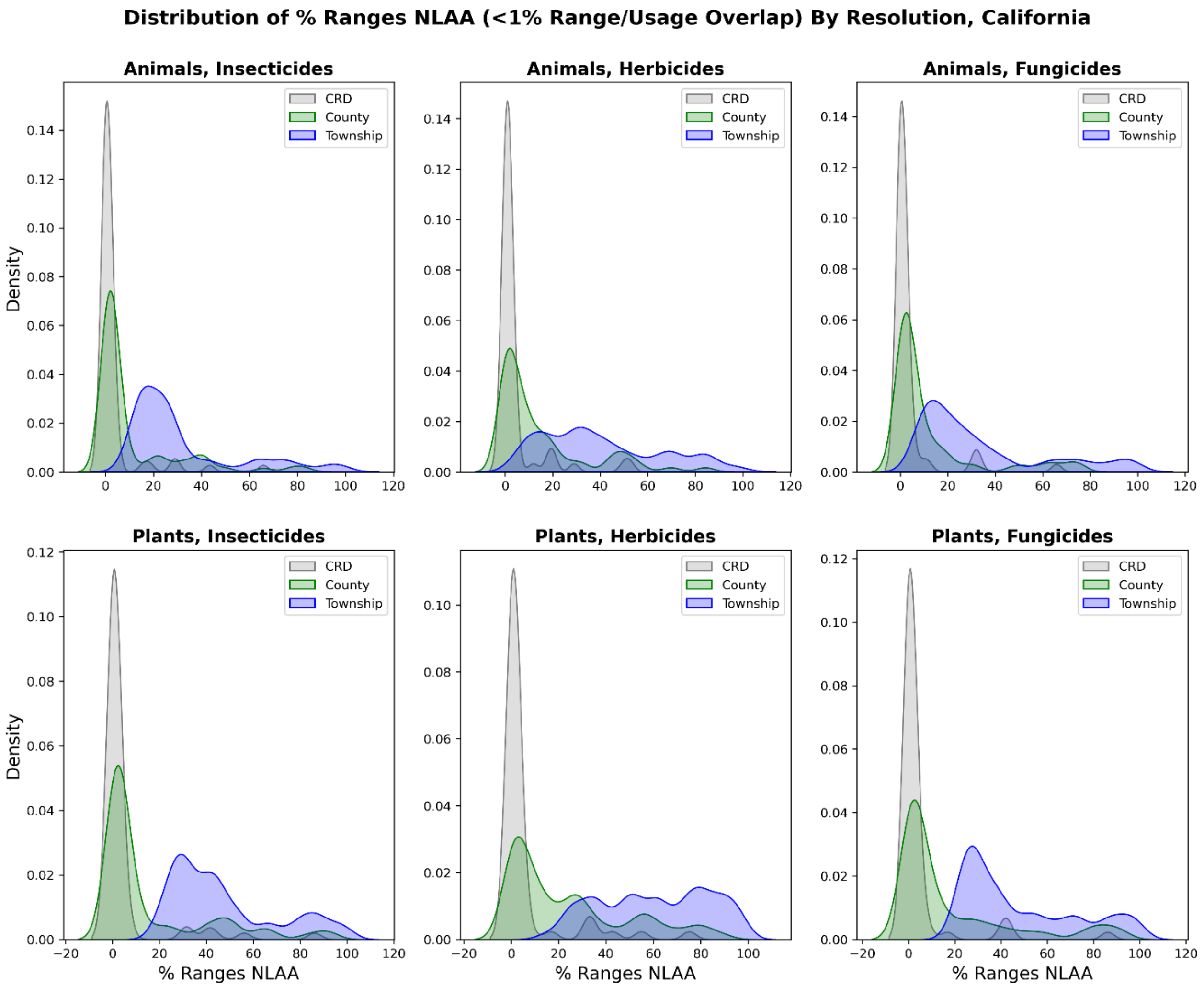

**Fig. S22.** Distribution of ranges designated NLAA in California, under either CRD, county, or township usage resolution across 223 compounds, visualized using kernel density estimates. Distributions are disaggregated by pesticide class (insecticide, herbicide, fungicide, from left to right), with results for animals given in the top panels and results for plants in the bottom panels.

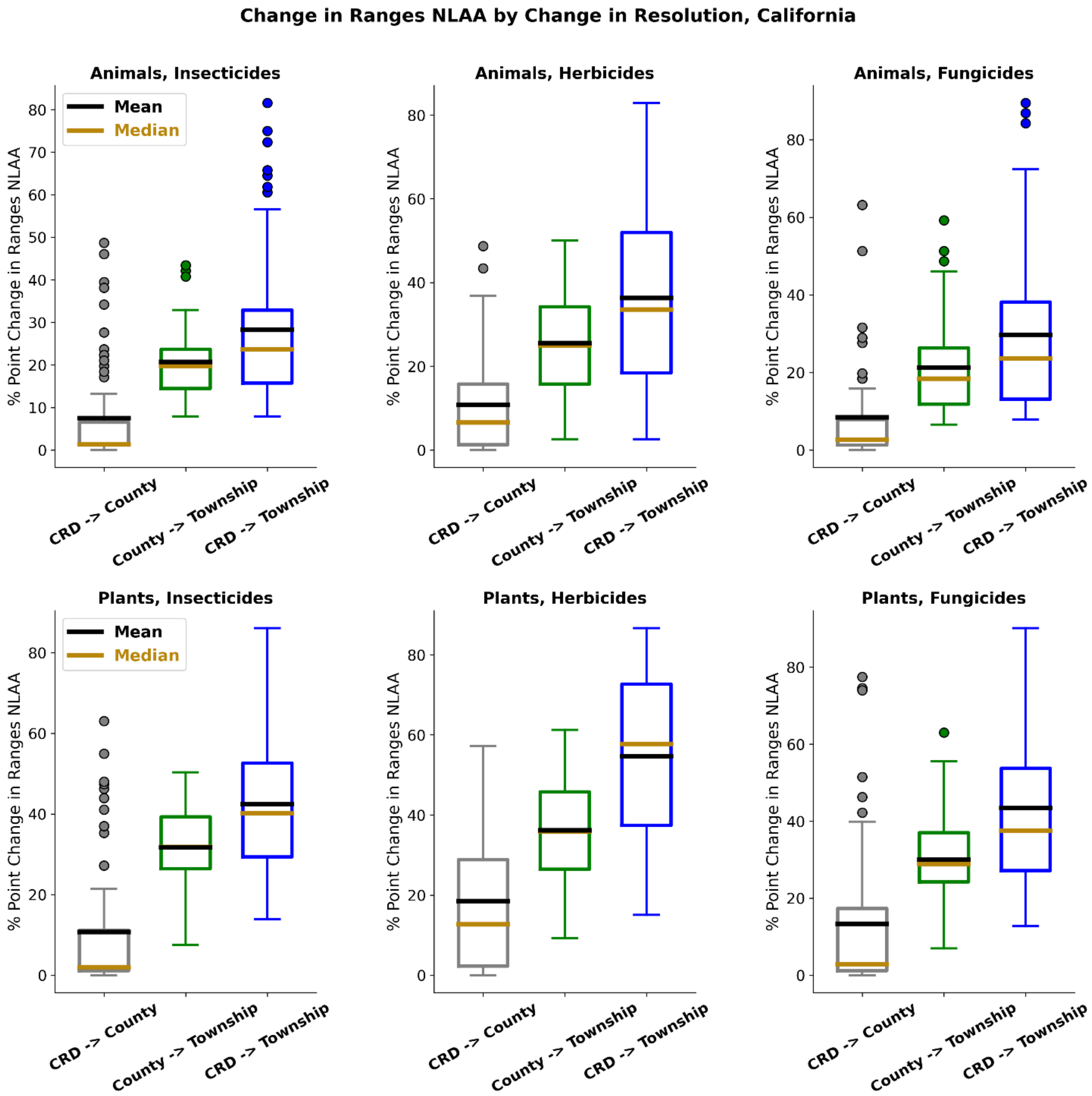

**Fig. S23.** Distributions of the number of species that switch from an LAA to NLAA designation, in California, when switching from either CRD to county, CRD to township, or county to township resolution. Distributions are disaggregated by pesticide class (insecticide, herbicide, fungicide, from left to right), with results for animals given in the top panels and results for plants in the bottom panels.

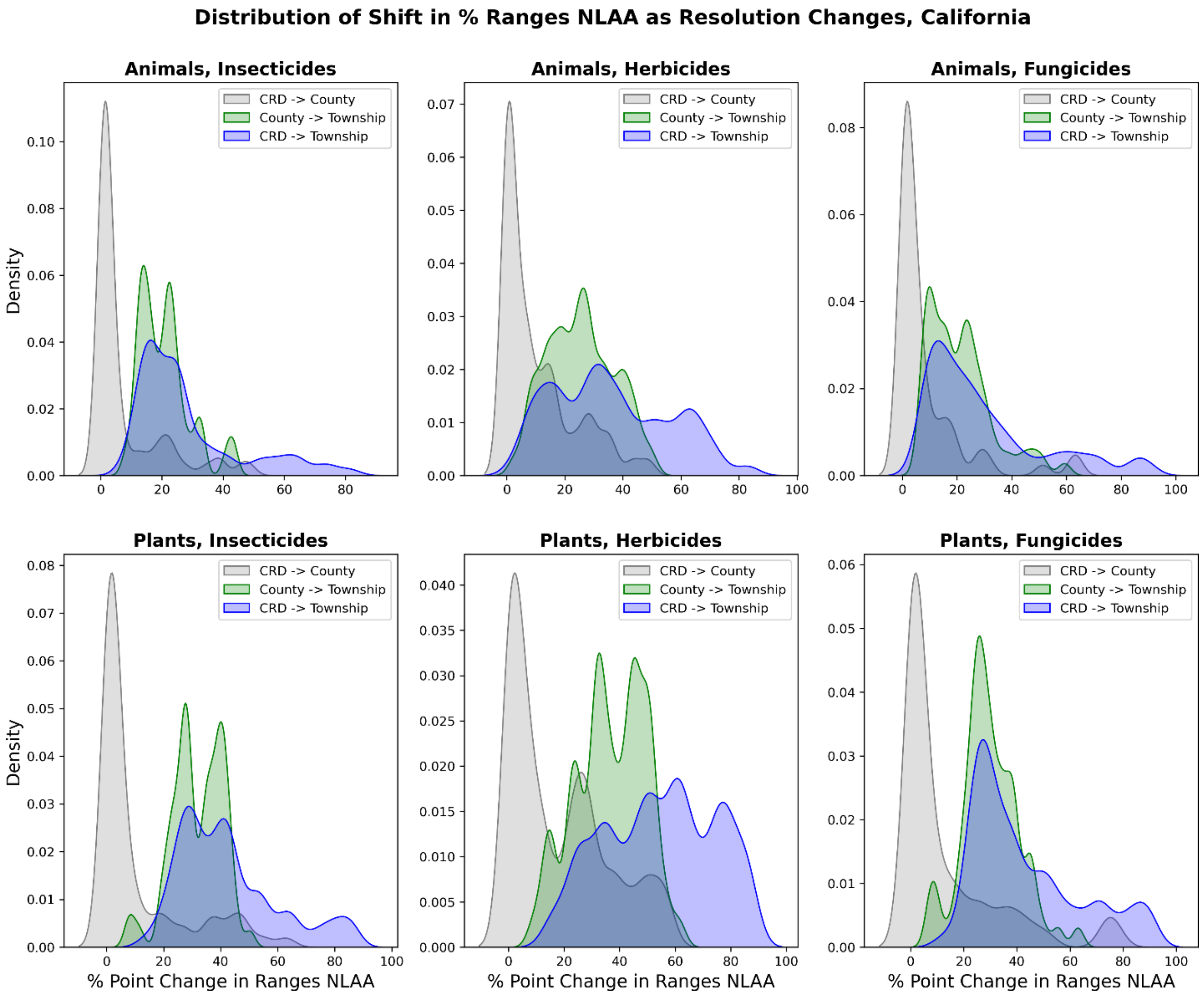

**Fig. S24.** Distributions of the number of species, visualized as kernel densities estimates, that switch from an LAA to NLAA designation, in California, when switching from either CRD to county, CRD to township, or county to township resolution. Distributions are disaggregated by pesticide class (insecticide, herbicide, fungicide, from left to right), with results for animals given in the top panels and results for plants in the bottom panels.

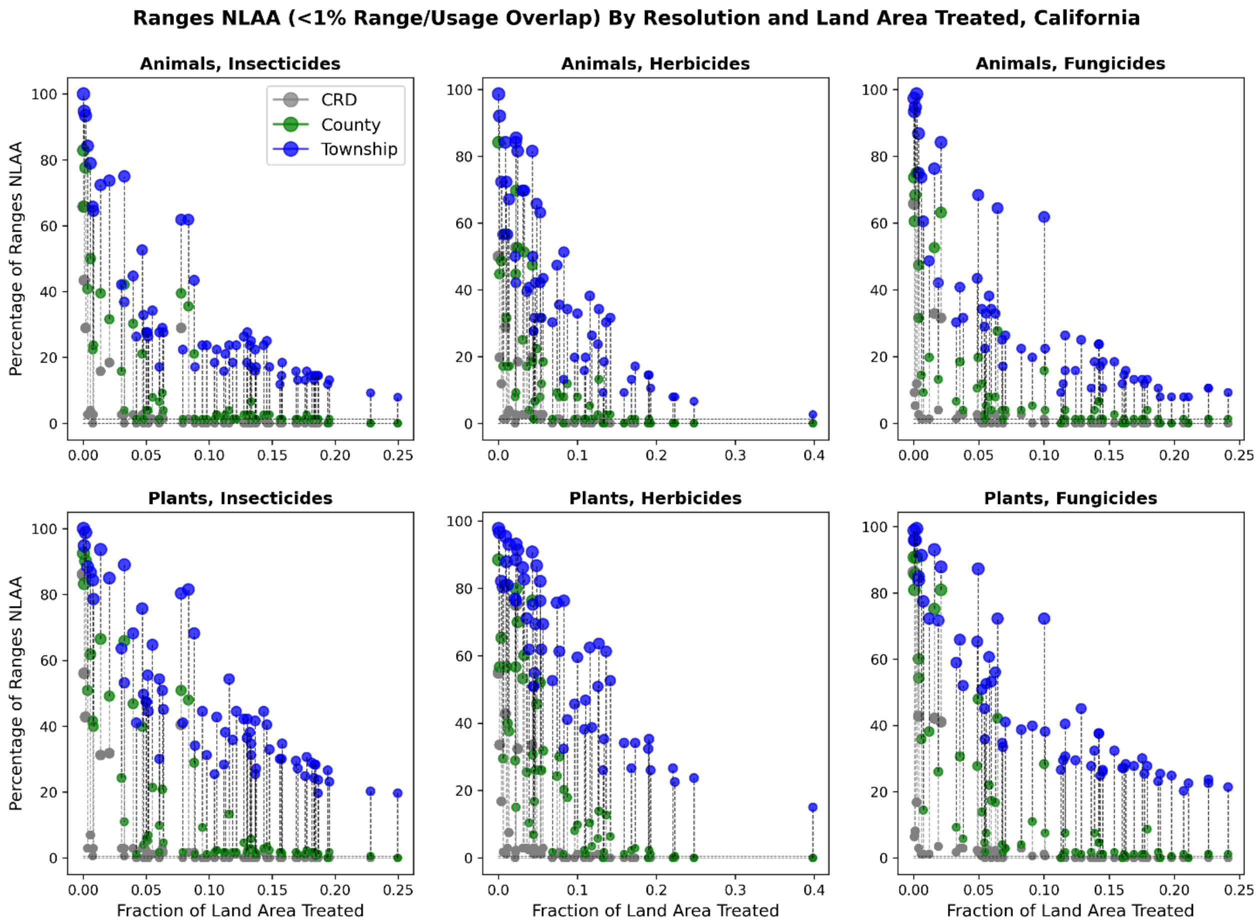

**Fig. S25.** Percentage of ranges designated NLAA for each of 223 compounds and for CRD, county, and township usage resolution, in California, as a function of the land area in California treated. That is, each set of three dots give the percentage of ranges NLAA for a single compound, with the horizontal location determined by how much land area in California was treated (maximum over 2013-2017). Results are disaggregated by pesticide class (insecticide, herbicide, fungicide, from left to right), with results for animals given in the top panels and results for plants in the bottom panels.

| Animal Ranges | | | |
|---|---|---|---|
| | **CRD** | **County** | **Township** |
| **Insecticides** | **189/5168 (3.66%)** | **578/5168 (11.18%)** | **1649/5168 (31.91%)** |
| **Herbicides** | **191/4256 (4.49%)** | **649/4256 (15.25%)** | **1737/4256 (40.81%)** |
| **Fungicides** | **173/4636 (3.73%)** | **566/4636 (12.21%)** | **1550/4636 (33.43%)** |
| **All Compounds** | **553/14060 (3.93%)** | **1793/14060 (12.75%)** | **4936/14060 (35.11%)** |
| **Plant Ranges** | | | |
| | **CRD** | **County** | **Township** |
| **Insecticides** | **591/11764 (5.02%)** | **1852/11764 (15.74%)** | **5584/11764 (47.47%)** |
| **Herbicides** | **596/9688 (6.15%)** | **2389/9688 (24.66%)** | **5893/9688 (60.83%)** |
| **Fungicides** | **492/10553 (4.66%)** | **1899/10553 (17.99%)** | **5071/10553 (48.05%)** |
| **All Compounds** | **1679/32005 (5.25%)** | **6140/32005 (19.18%)** | **16548/32005 (51.70%)** |
| **Plants/Animals Combined** | | | |
| | **CRD** | **County** | **Township** |
| **Insecticides** | **780/16932 (4.61%)** | **2430/16932 (14.35%)** | **7233/16932 (42.72%)** |
| **Herbicides** | **787/13944 (5.64%)** | **3038/13944 (21.79%)** | **7630/13944 (54.72%)** |
| **Fungicides** | **665/15189 (4.38%)** | **2465/15189 (16.23%)** | **6621/15189 (43.59%)** |
| **All Compounds** | **2232/46065 (4.85%)** | **7933/46065 (17.22%)** | **21484/46065 (46.64%)** |

**Table S1.** Total number of range/compound pairs found to be NLAA (<1% range/compound overlap) under different pesticide usage resolutions, in California. Cells give the number of range/compound pairs NLAA out of the total, with the percentage in parentheses. Note that compounds without any presence in CA are excluded from these results. Results are disaggregated by compound class, and are presented for animals, plants, and animals/plants combined.

**5. Representative maps of usage footprint and range classification, by usage scale: California**

We graphically illustrate the (maximum over 2013-2017) usage footprints at township, county, and CRD scale, along with the ranges that can be considered NLAA (i.e., <1% overlap between range and usage) at each scale. Ranges are indicated by their centroid, with size scaled in proportion to range area. Here we provide maps for the top 10 pesticides in each class by mass, nationally, that have any presence in California for animals and plants separately, in Figs S26-S31. A simple interactive iPython/jupyter notebook that provides results for all compounds is available at the link (https://datadryad.org/stash/share/GdNHpEf1xMa9ZRVA4Eo2xiTWlF7cXrzNeQzzhcmLNmU) in Eikenberry et al. (2022). A screenshot of this app is given in Fig. S32.

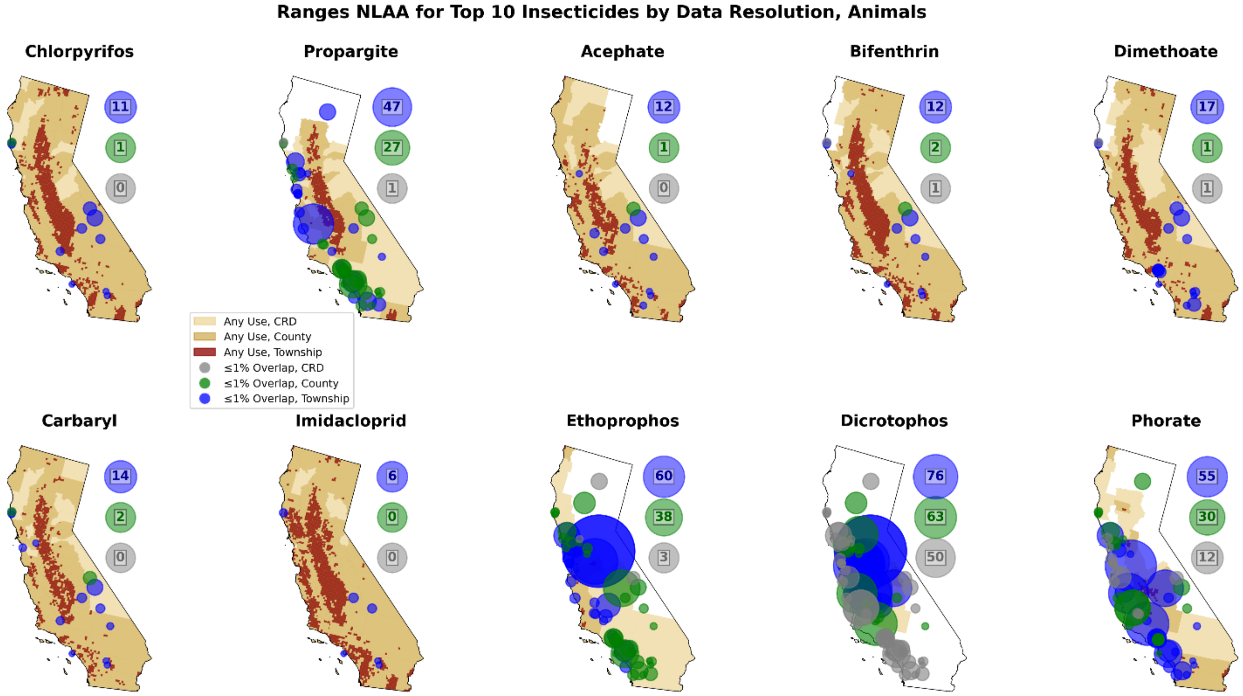

**Fig. S26.** Maps show the binary maximum usage footprint (at CRD, county, and township usage resolutions) for the top 10 insecticides by mass (nationally) that were applied in California, along with the ranges that are designated NLAA (<1% overlap) at each usage resolution. Bubbles on the map are centered at the range centroid, and scale in size based on the total range area. The bubbles to the right of the maps with inscribed numbers indicate the total number of ranges NLAA for each resolution. Results are shown for animal ranges.

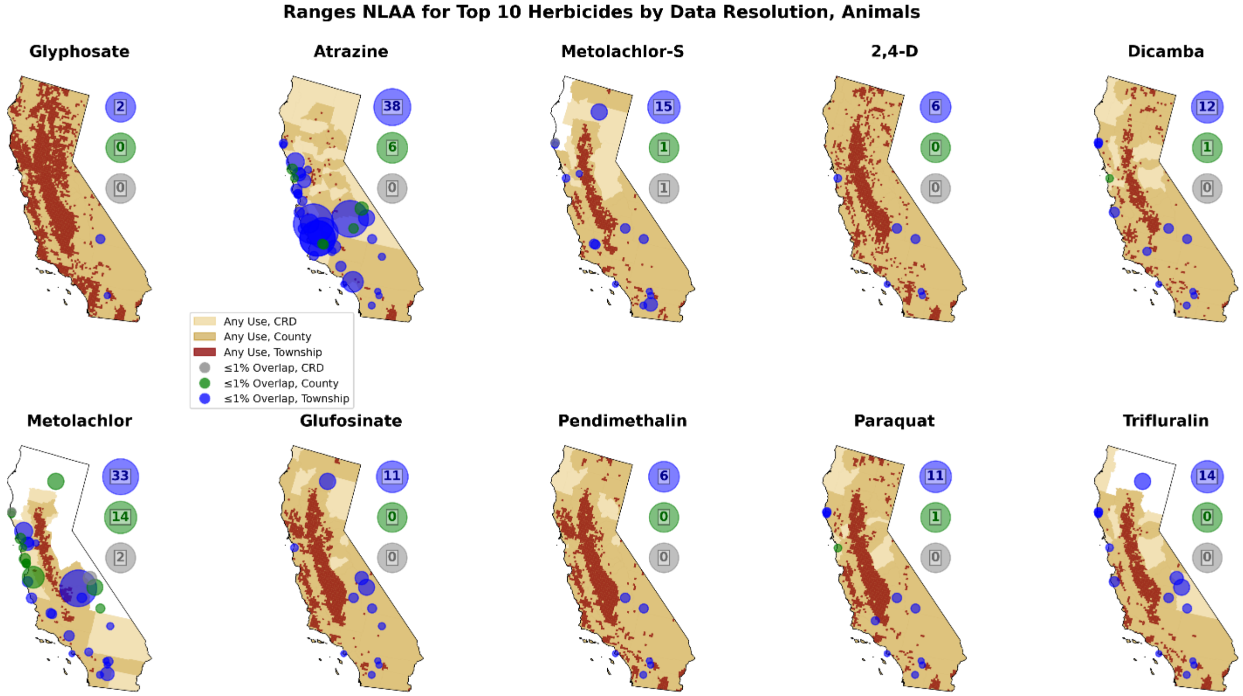

**Fig. S27.** Similar to Fig. S26, maps give the binary maximum usage footprints, locations and areas of ranges designated NLAA at each usage resolution, and total numbers of ranges NLAA at each resolution. Results are shown for the top 10 herbicides nationally with any application in California, and for animals.

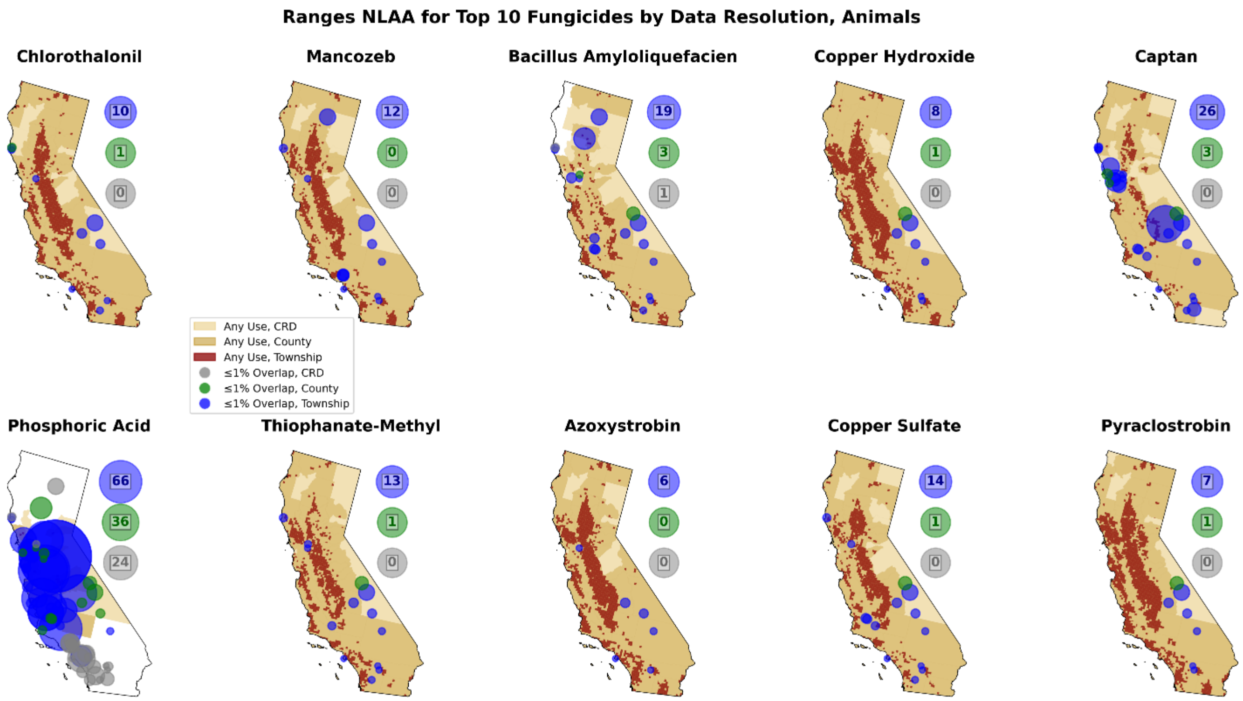

**Fig. S28.** Similar to Fig. S26, maps give the binary maximum usage footprints, locations and areas of ranges designated NLAA at each usage resolution, and total numbers of ranges NLAA at each resolution. Results are shown for the top 10 fungicides nationally with any application in California, and for animals.

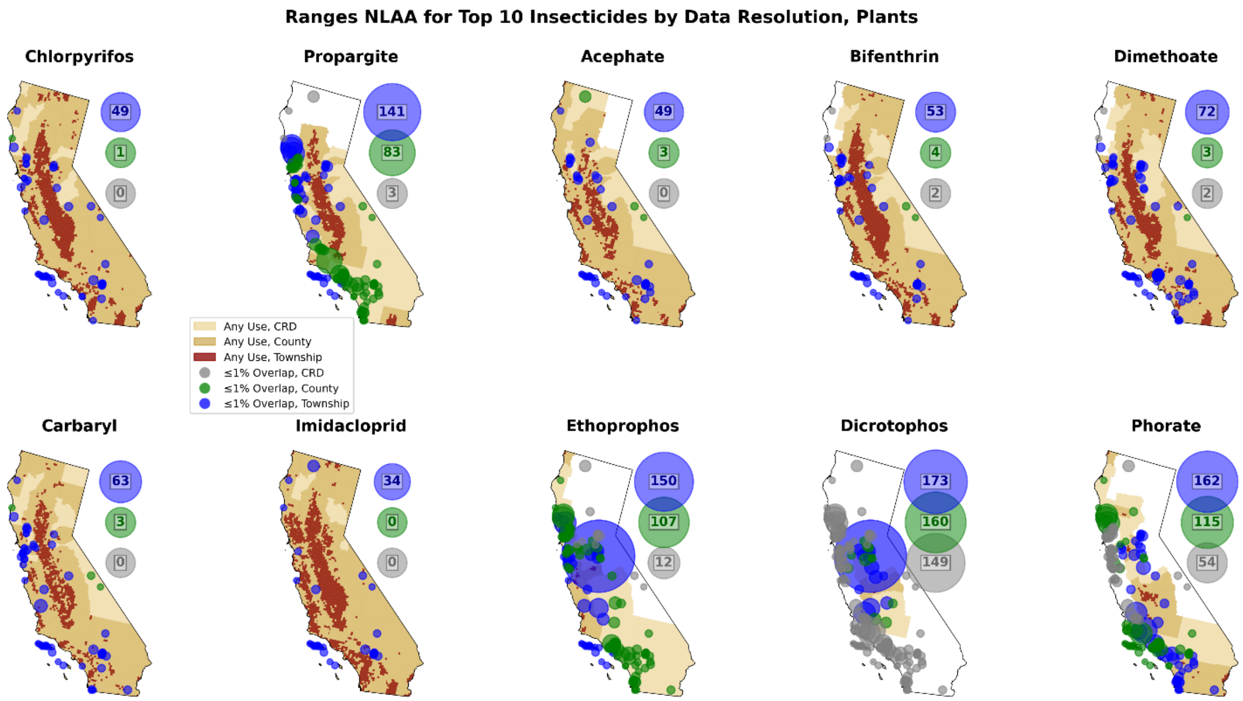

***Fig. S29.*** Similar to Fig. S26, maps give the binary maximum usage footprints, locations and areas of ranges designated NLAA at each usage resolution, and total numbers of ranges NLAA at each resolution. Results are shown for the top 10 insecticides nationally with any application in California, and for plants.

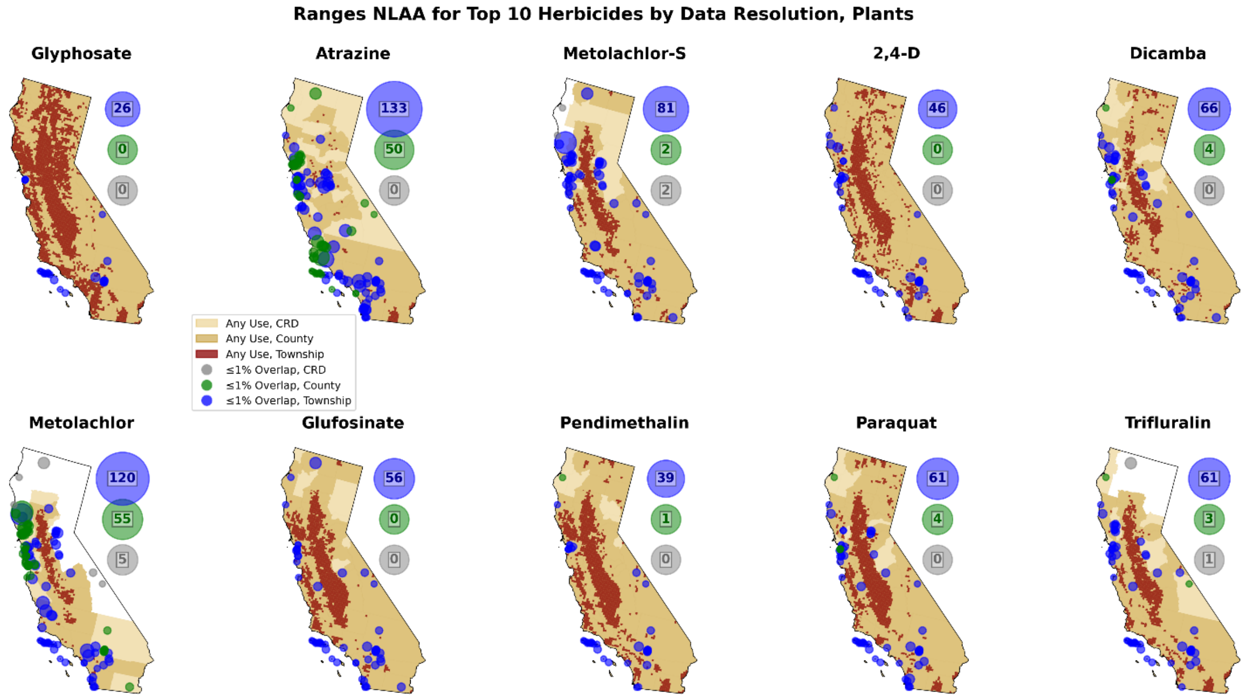

**Fig. S30.** Similar to Fig. S26, maps give the binary maximum usage footprints, locations and areas of ranges designated NLAA at each usage resolution, and total numbers of ranges NLAA at each resolution. Results are shown for the top 10 herbicides nationally with any application in California, and for plants.

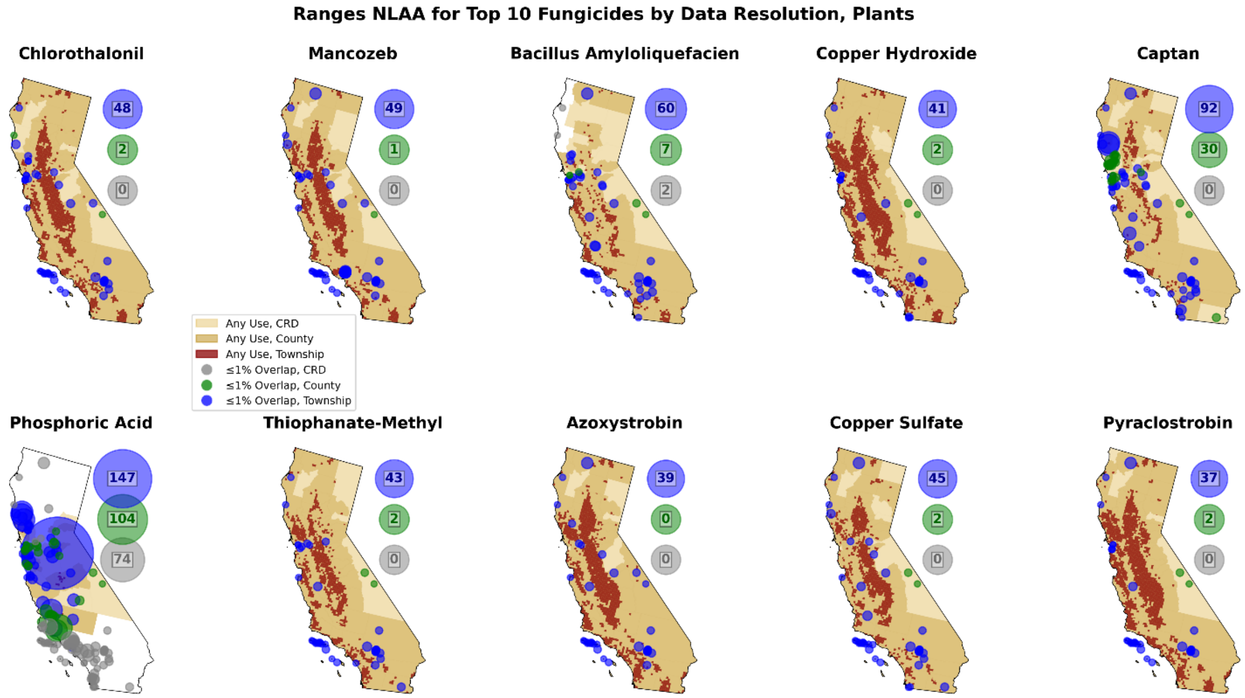

**Fig. 31.** Similar to Fig. S26, maps give the binary maximum usage footprints, locations and areas of ranges designated NLAA at each usage resolution, and total numbers of ranges NLAA at each

resolution. Results are shown for the top 10 fungicides nationally with any application in California, and for plants.

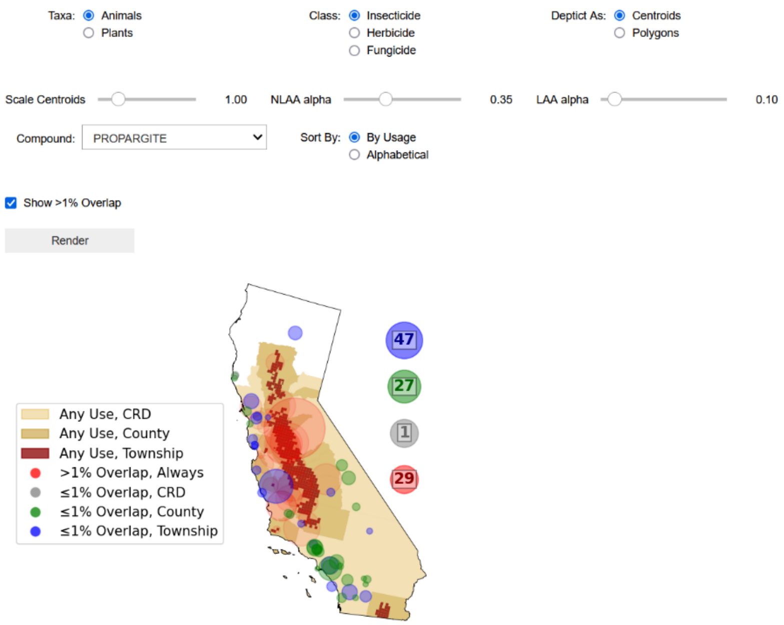

**Fig. S32.** Screenshot of Jupyter Notebook app to visualize overlap between pesticide usage footprint at different resolutions and species' ranges. Screenshot has been cropped for clarity.

#### 6. Classifier and performance in California

For California endemic species and for each compound, we classified animal (plant) ranges as benefiting from township-level data if they were determined to be LAA using county-level usage data, but NLAA under township usage data. We then performed a series of logistic regressions, with the response variable whether a range benefitted from township data, and the regressors either (1) range area, (2) range agricultural land fraction (maximum over 2013-2017), (3) pesticide use intensity within the range area at county scale (kg per km2), or (4) all three. We characterized each model performance with an ROC curve, using the area under the curve (AUC) as our primary metric. We used bootstrap resampling to derive confidence intervals for both the ROC curve and AUC, and to determine if performance varied significantly between the models.

To classify ranges as either likely to benefit or not, we used logistic regression as indicated, with the probability cutoff chosen as the top left point of the ROC curve. Representative results are given for the pesticide propargite and animal ranges in Fig. S33, which shows the comparative performance of the four models, as well as maps of all true and false positives, true and false negatives, and calculated sensitivity and specificity. Results for propargite and plants are similarly given in Fig. S34.

We compared overall model performance across all 223 compounds by determining the AUC for each model/compound, followed by bootstrap resampling to yield bootstrap distributions for the mean AUC. The AUC distributions, for animal ranges, are given in Fig. S35, with Model 4 clearly outperforming the other three. Somewhat similar model performance was observed for plant ranges, as shown in Fig. S36, although Model 1 (range area) and Model 4 (all predictors) had indistinguishable performance for plants. We also performed these analyses using just insecticides, herbicides, or fungicides, with consistent results (not shown).

Overall, the classifiers performed reasonably well in California, with mean ± std AUC, sensitivity, and specificity of 86±7%, 82±9%, and 83±10%, respectively, for animal ranges, and 78±8%, 75±10%, 73±10%, for plant ranges, respectively, across all range/compound combinations.

Results were very similar when disaggregating by compound class, and when we restricted our consideration to only those compounds where at least 20 ranges switched from LAA to NLAA with township usage data.

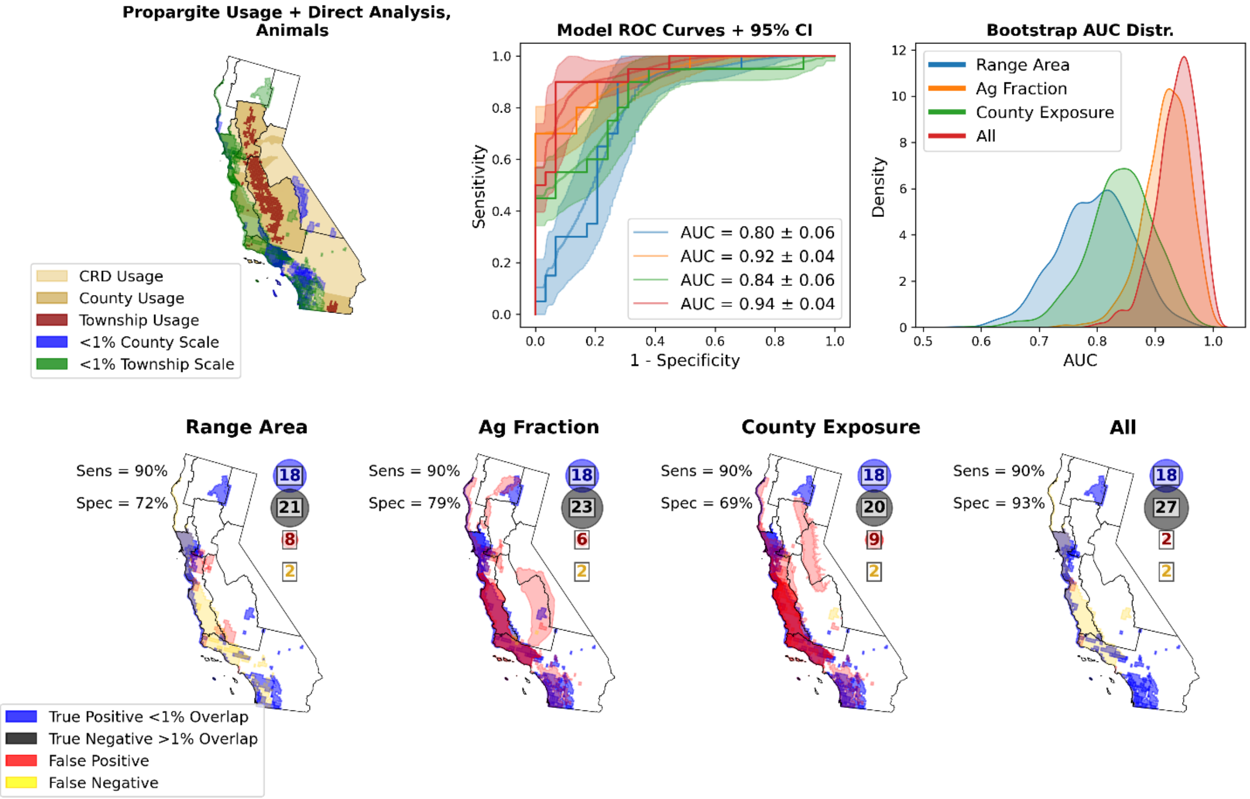

**Fig. S33.** Performance of the four classification models on *animal* ranges in CA for the insecticide propargite. The top left panel shows which ranges are classified as NLAA (<1% usage overlap) under direct analysis for county and township usage resolution. The top middle panel gives the ROC curves for each of the four models with 95% bootstrap Cis. On the top left the bootstrap distribution of the AUC for each model is given. The lower panels show which and how many ranges are correctly predicted to benefit from township usage data (true positives), which are correctly predicted not to benefit (true negatives), which are incorrectly predicted to benefit (false positive) and which are incorrectly predicted to not benefit (false negative). The resulting sensitivities and specificities are also indicated for each model.

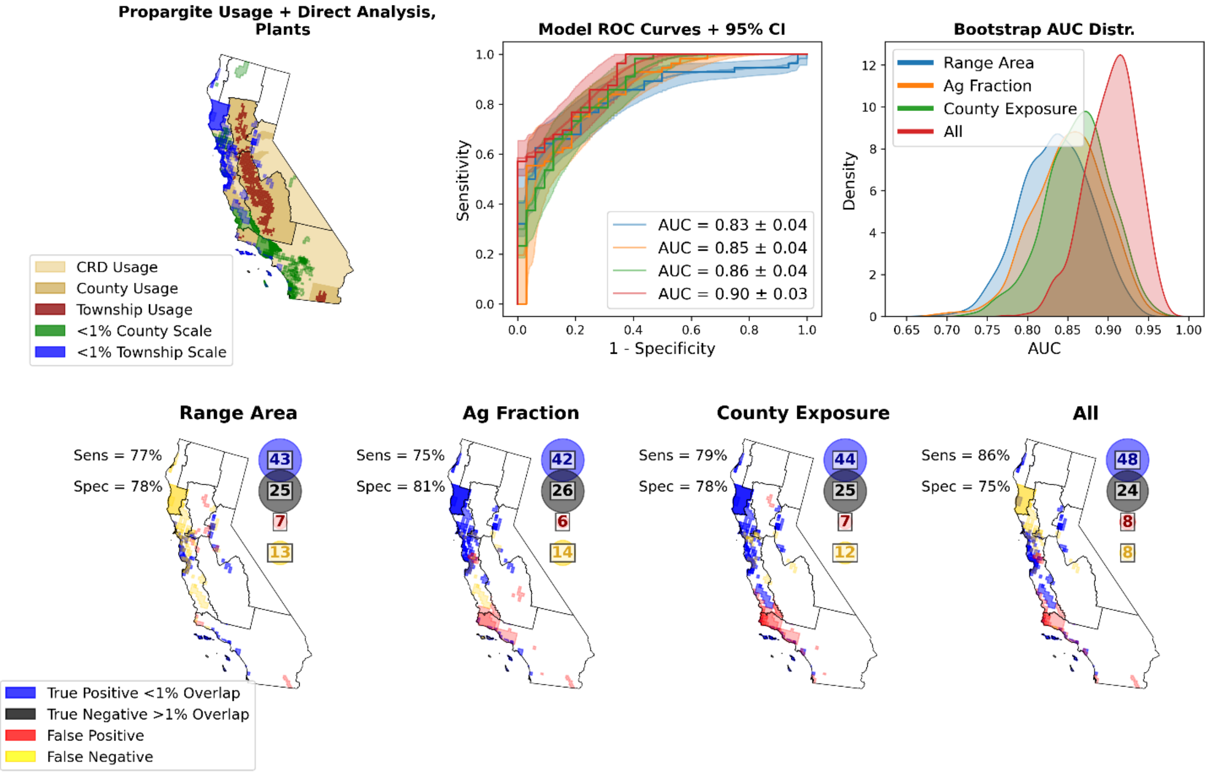

**Fig. S34.** Performance of the four classification models on *plant* ranges in CA for the insecticide propargite. The top left panel shows which ranges are classified as NLAA (<1% usage overlap) under direct analysis for county and township usage resolution. The top middle panel gives the ROC curves for each of the four models with 95% bootstrap Cis. On the top left the bootstrap distribution of the AUC for each model is given. The lower panels show which and how many ranges are correctly predicted to benefit from township usage data (true positives), which are correctly predicted not to benefit (true negatives), which are incorrectly predicted to benefit (false positive) and which are incorrectly predicted to not benefit (false negative). The resulting sensitivities and specificities are also indicated for each model.

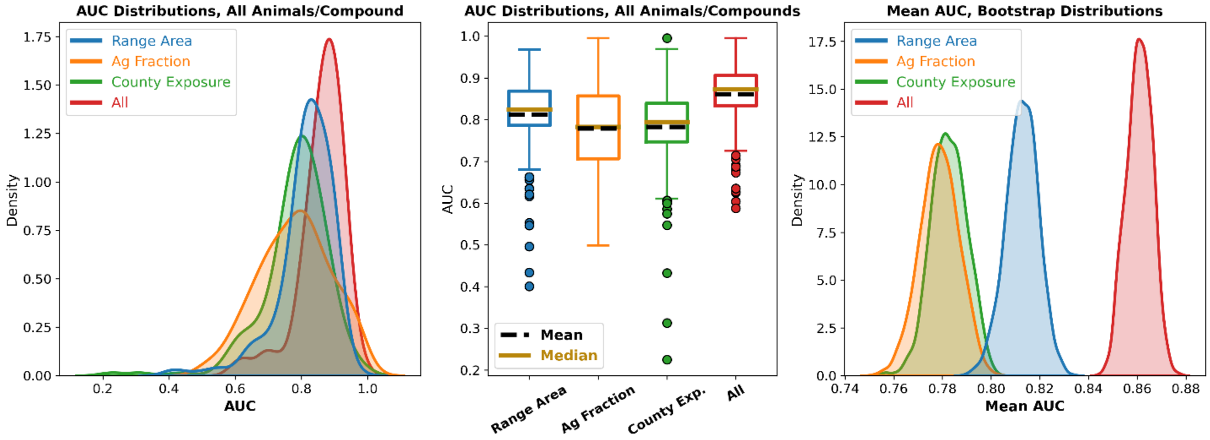

**Fig. S35.** Aggregate performance of the classifiers for animal ranges across all pesticides. The left panel gives the raw distributions of the AUC for each model as a kernel density estimation, and the middle panel gives these distributions as box plots. The right panel gives the bootstrap distributions of the mean AUC, demonstrating that the model with all predictors performs significantly better than the others.

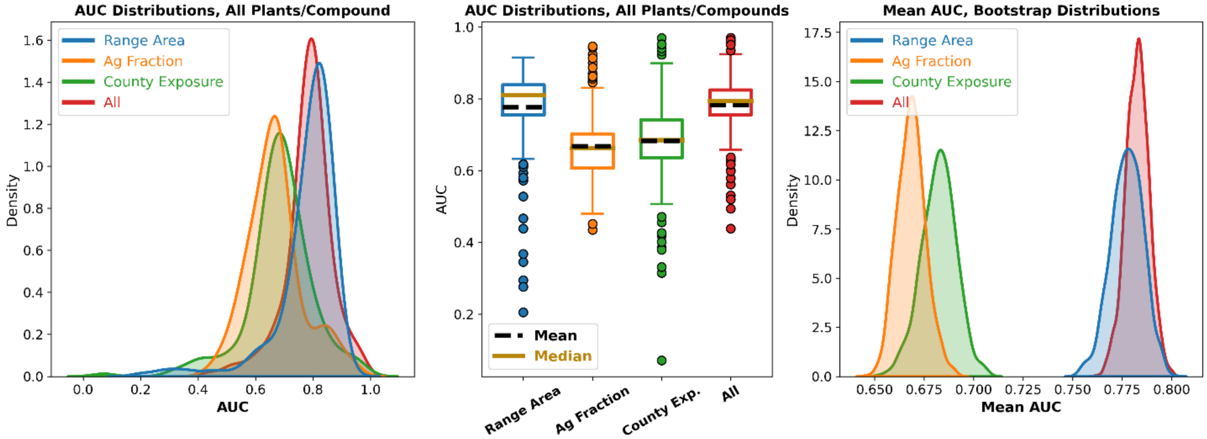

**Fig. S36.** Aggregate performance of the classifiers for plant ranges across all pesticides. The left panel gives the raw distributions of the AUC for each model as a kernel density estimation, and the middle panel gives these distributions as box plots. The right panel gives the bootstrap distributions of the mean AUC, demonstrating that either range area alone or all predictors perform equally well for plants.

| | | | |
|---|---|---|---|
| **Animals** | | | |
| **All compounds (mean ± sd)** | | | |
| **Model Predictors** | **AUC** | **Sensitivity** | **Specificity** |
| ***Range Area*** | **81.23 ± 8.67%** | **79.30 ± 9.01%** | **76.05 ± 9.56%** |
| ***Ag Fraction*** | **77.96 ± 11.93%** | **74.94 ± 12.77%** | **76.17 ± 11.76%** |
| ***County Exposure*** | **78.33 ± 10.09%** | **77.25 ± 11.63%** | **78.03 ± 11.06%** |
| ***All*** | **86.12 ± 7.04%** | **81.61 ± 9.49%** | **82.63 ± 9.83%** |
| **Insecticides** | | | |
| ***All*** | **85.75 ± 6.99%** | **80.56 ± 8.52%** | **82.15 ± 9.91%** |
| **Herbicides** | | | |
| ***All*** | **85.62 ± 8.42%** | **83.15 ± 10.37%** | **81.25 ± 10.31%** |
| **Fungicides** | | | |
| ***All*** | **86.99 ± 5.57%** | **81.34 ± 9.64%** | **84.43 ± 9.15%** |
| **Plants** | | | |
| **All compounds (mean ± sd)** | | | |
| **Model Predictors** | **AUC** | **Sensitivity** | **Specificity** |
| ***Range Area*** | **76.42 ± 12.82%** | **73.48 ± 13.54%** | **72.72 ± 7.85%** |
| ***Ag Fraction*** | **66.79 ± 9.88%** | **65.10 ± 9.94%** | **64.40 ± 12.67%** |
| ***County Exposure*** | **67.95 ± 11.78%** | **63.91 ± 14.66%** | **73.50 ± 11.78%** |
| ***All*** | **77.81 ± 8.43%** | **73.26 ± 10.64%** | **73.41 ± 10.09%** |
| **Insecticides** | | | |
| ***All*** | **77.03 ± 7.60%** | **72.55 ± 10.66%** | **72.33 ± 8.76%** |
| **Herbicides** | | | |
| ***All*** | **80.60 ± 9.63%** | **76.36 ± 11.41%** | **75.99 ± 9.89%** |
| **Fungicides** | | | |
| ***All*** | **76.11 ± 7.56%** | **71.18 ± 9.33%** | **72.24 ± 11.29%** |

**Table S2.** Performance of the different classification models in California. The models predict which ranges are likely to benefit from township usage data, compared to just county scale data.

**7. Predictions for contiguous US**

For the contiguous US (outside of California), we directly calculated the overlap between maximum pesticide usage footprints and endangered species' ranges for CRD and county resolution usage. We then used the regression/classification models trained on California data to predict which range/compound pairs not restricted to California would benefit (switch from LAA to NLAA status) from township resolution data. Note that predictions were not possible for 39 out of 223 compounds, either due to no usage in California, or too few classified as NLAA in California, and these are excluded from the results presented. The distributions of ranges predicted to be classified as NLAA (i.e., <1% overlap) at each usage scale are summarized in Figs. S37 and S38; the shifts in these distributions under changes in usage resolutions are given in Figs. S39 and S40. Figs. S41-S46 give the results for the top 10 compounds in each class for animals and plant, separately.

We visualized those regions that most benefit to township vs. CRD resolution usage data, in terms of number of range/compound pairs predicted to be NLAA for each county, as either choropleth or bubble plots. Figure 3 of the main text depicts these results as choropleth maps, while bubble plots are given in Fig. S47.

We also consider the case where county-scale data is available at baseline and visualize those regions that benefit from township data. These results, as choropleth and bubble plots, are given in Figs. S49 and S50. Exact results disaggregated by broad pesticide class (insecticide, herbicide, fungicide) and animals vs. plants, are given in Table 1 of the main text.

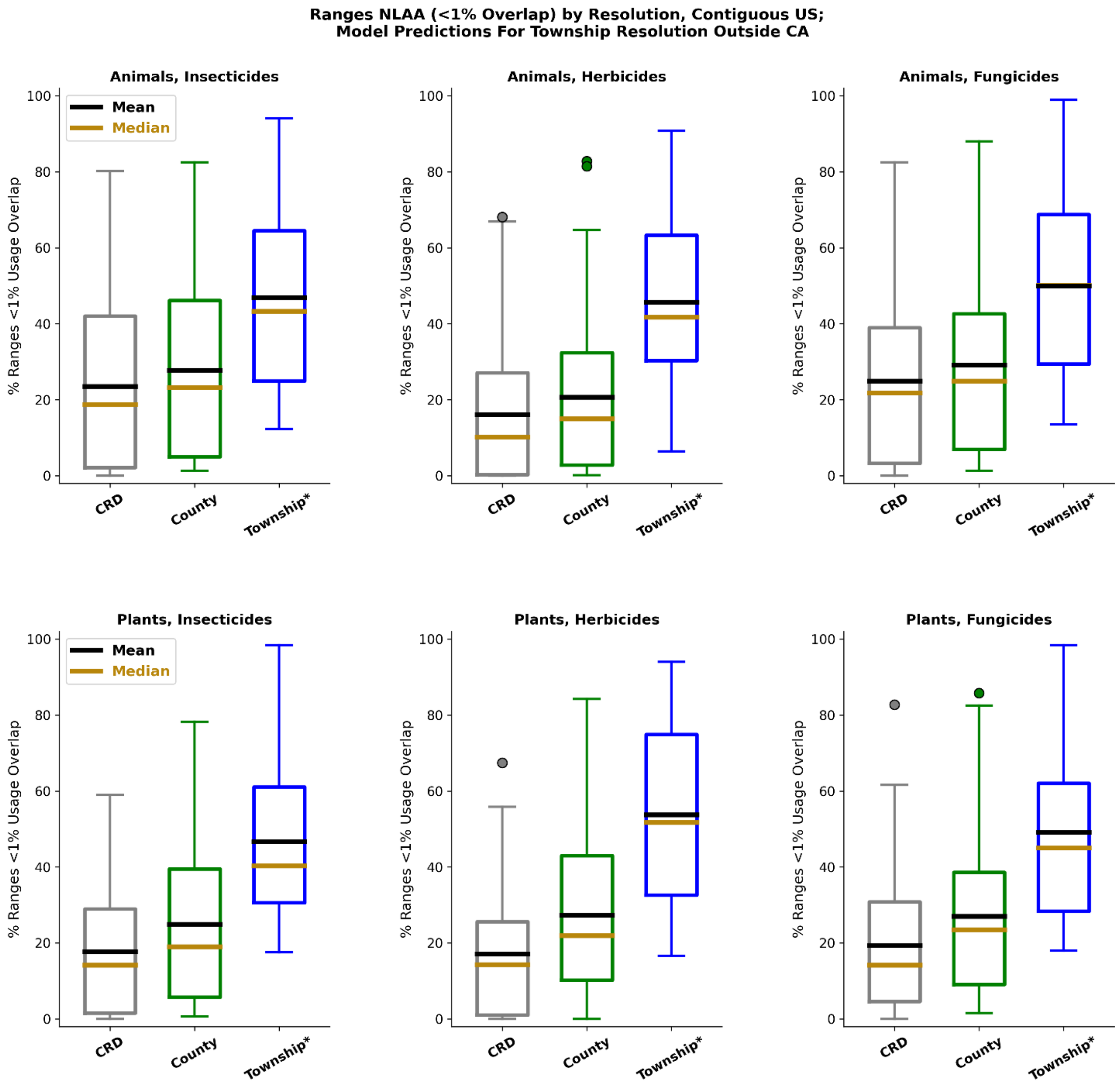

**Fig. S37.** Distribution of ranges designated NLAA across the contiguous US, under either CRD, county, or township usage resolution across 184 compounds, visualized as boxplots. Direct overlap analysis is used for CRD and county results, as well as township results for ranges strictly contained within California, while model predictions are used for township resolution outside of California. Distributions are disaggregated by pesticide class (insecticide, herbicide, fungicide, from left to right), with results for animals given in the top panels and results for plants in the bottom panels.

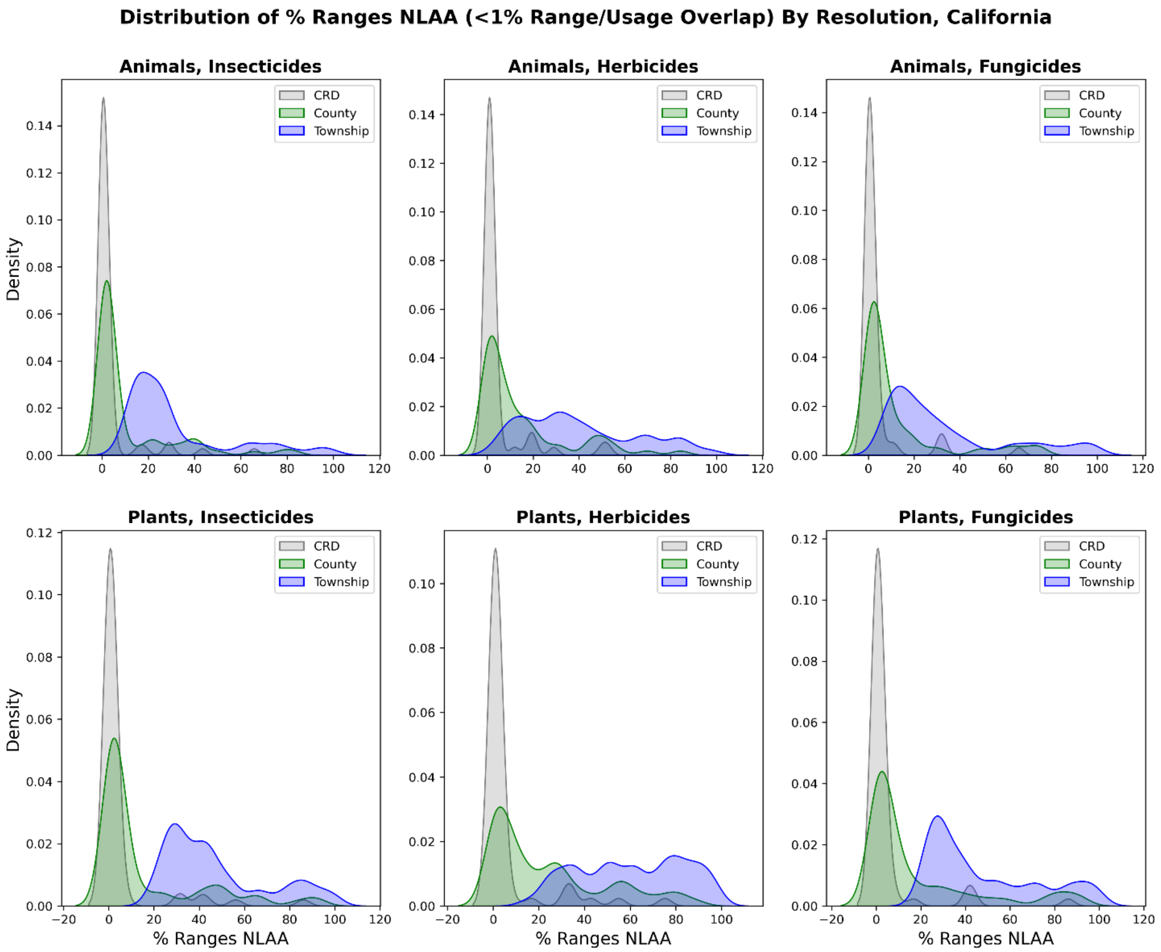

**Fig. S38.** Distribution of ranges designated NLAA across the contiguous US, under either CRD, county, or township usage resolution across 184 compounds, visualized using kernel density estimates. Direct overlap analysis is used for CRD and county results, as well as township results for ranges strictly contained within California, while model predictions are used for township resolution outside of California. Distributions are disaggregated by pesticide class (insecticide, herbicide, fungicide, from left to right), with results for animals given in the top panels and results for plants in the bottom panels.

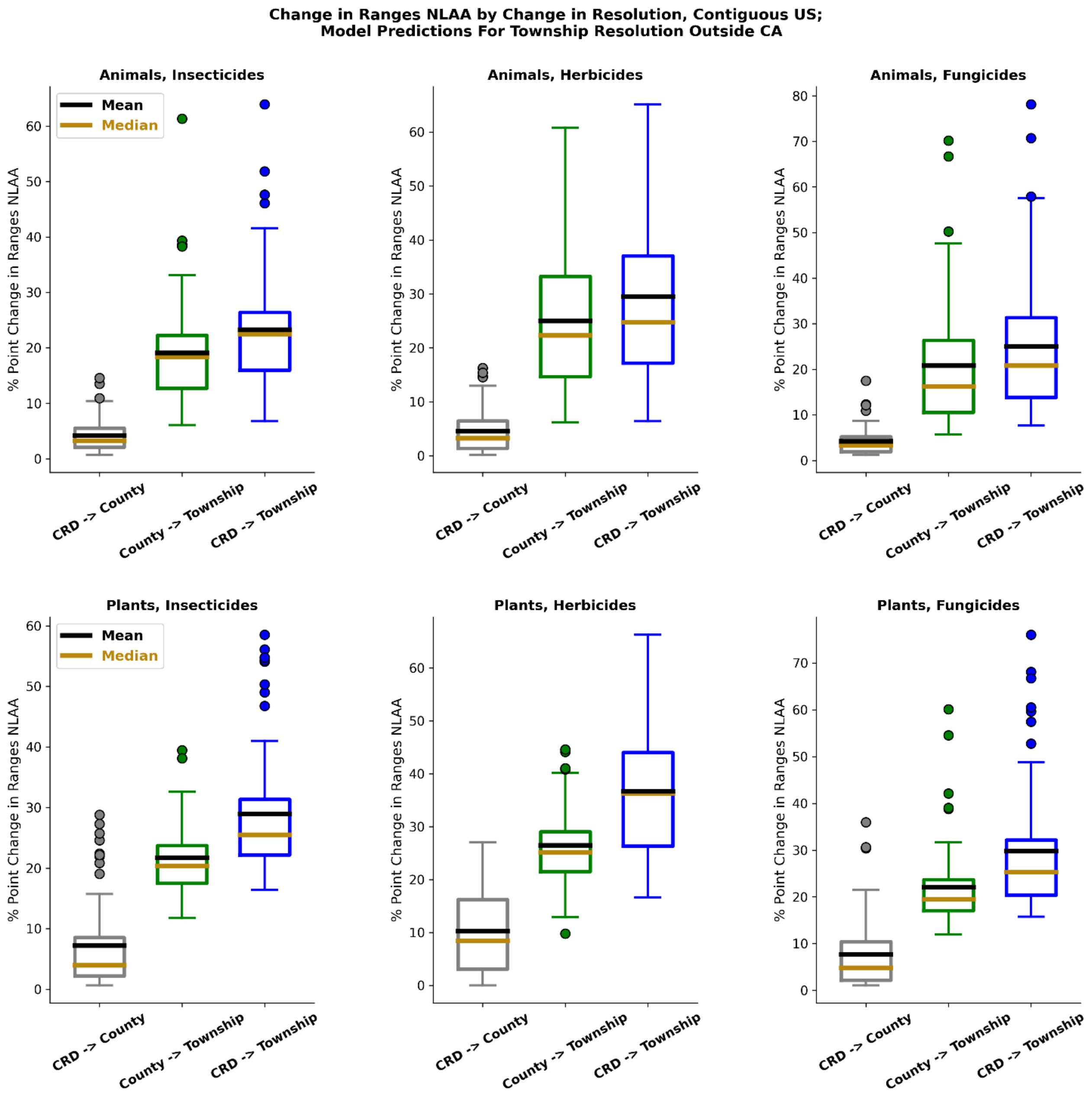

**Fig. S39.** Distributions of the number of species that switch from an LAA to NLAA designation, across the contiguous US, when switching from either CRD to county, CRD to township, or county to township resolution. Direct overlap analysis is used for CRD and county results, as well as township results for ranges strictly contained within California, while model predictions are used for township resolution outside of California. Distributions are disaggregated by pesticide class (insecticide, herbicide, fungicide, from left to right), with results for animals given in the top panels and results for plants in the bottom panels.

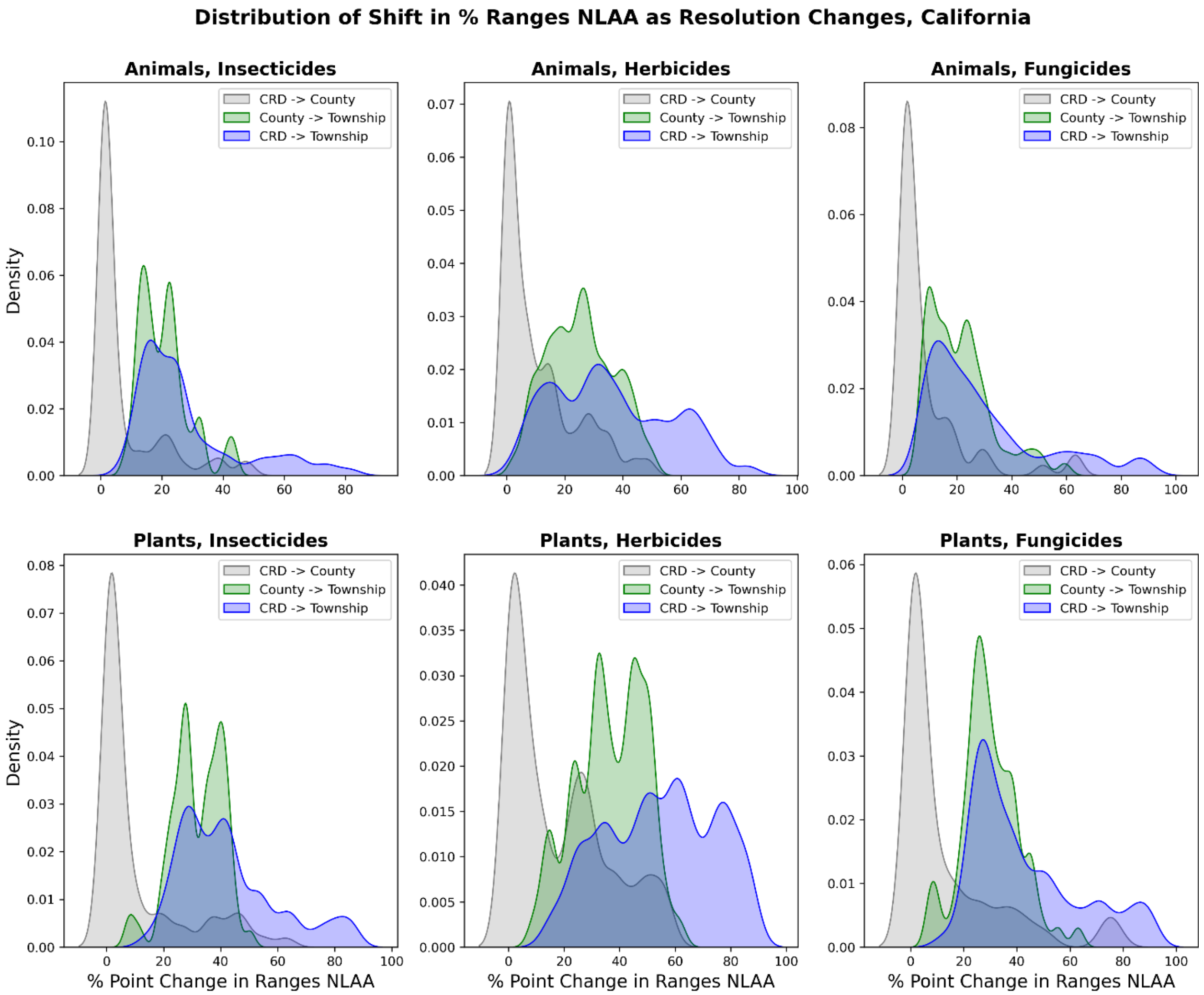

**Fig. S40.** Distributions of the number of species, visualized as kernel densities estimates, that switch from an LAA to NLAA designation, across the contiguous US, when switching from either CRD to county, CRD to township, or county to township resolution. Direct overlap analysis is used for CRD and county results, as well as township results for ranges strictly contained within California, while model predictions are used for township resolution outside of California. Distributions are disaggregated by pesticide class (insecticide, herbicide, fungicide, from left to right), with results for animals given in the top panels and results for plants in the bottom panels.

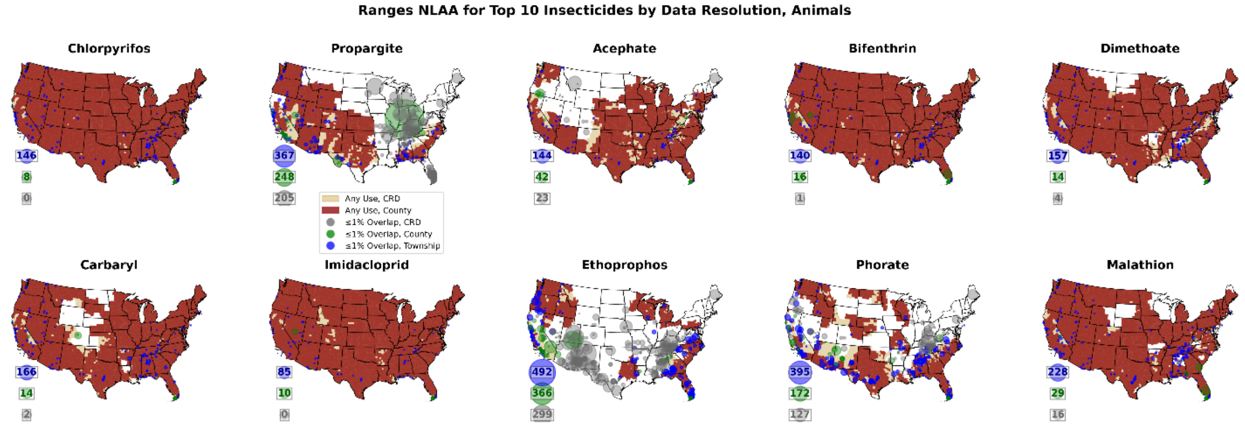

**Fig. S41.** Maps show the binary maximum usage footprint (at CRD, county, and township usage resolutions) for the top 10 insecticides by mass (nationally), along with the ranges that are designated NLAA (<1% overlap) at each usage resolution. Direct overlap analysis is used for CRD and county results, as well as township results for ranges strictly contained within California, while model predictions are used for township resolution outside of California. Bubbles on the map are centered at the range centroid, and scale in size based on the total range area. The bubbles to the right of the maps with inscribed numbers indicate the total number of ranges NLAA for each resolution. Results are shown for animal ranges.

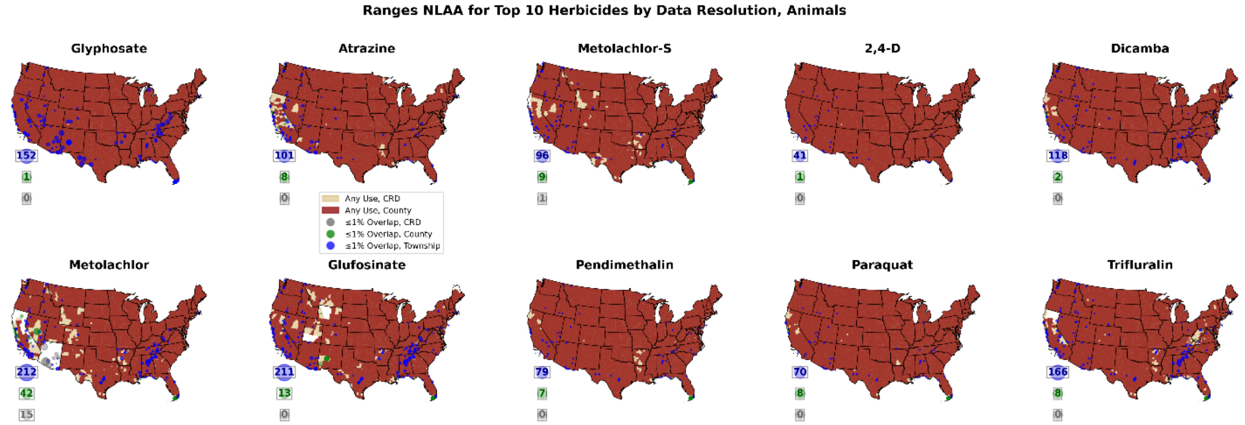

**Fig. S42.** Similar to Fig. S41, maps give the binary maximum usage footprints, locations and areas of ranges designated NLAA at each usage resolution, and total numbers of ranges NLAA at each resolution. Model predictions are used for township resolution outside CA; results are shown for the top 10 herbicides nationally, for animals.

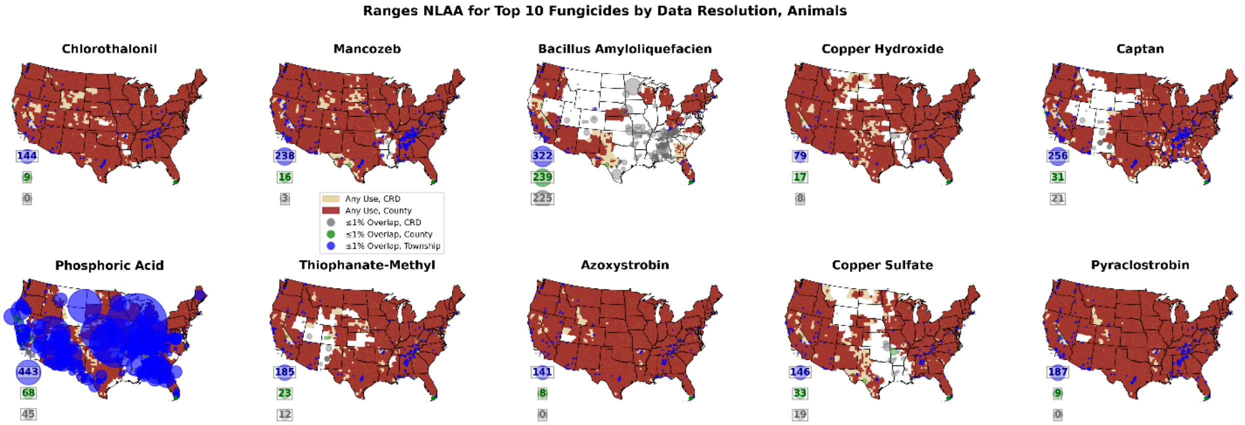

**Fig. S43.** Similar to Fig. S41, maps give the binary maximum usage footprints, locations and areas of ranges designated NLAA at each usage resolution, and total numbers of ranges NLAA at each resolution. Model predictions are used for township resolution outside CA; results are shown for the top 10 fungicides nationally, for animals.

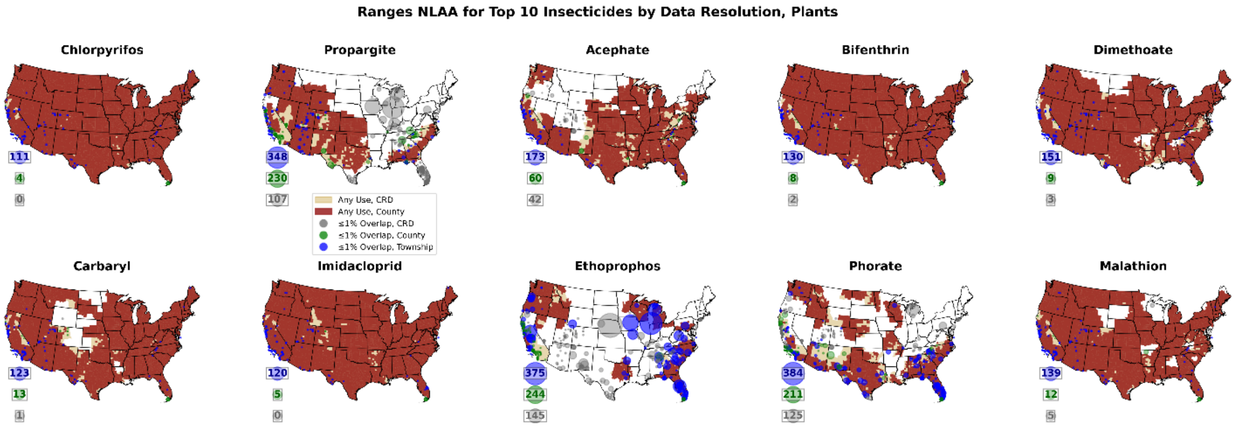

**Fig. S44.** Similar to Fig. S41, maps give the binary maximum usage footprints, locations and areas of ranges designated NLAA at each usage resolution, and total numbers of ranges NLAA at each resolution. Model predictions are used for township resolution outside CA; results are shown for the top 10 insecticides nationally, for plants.

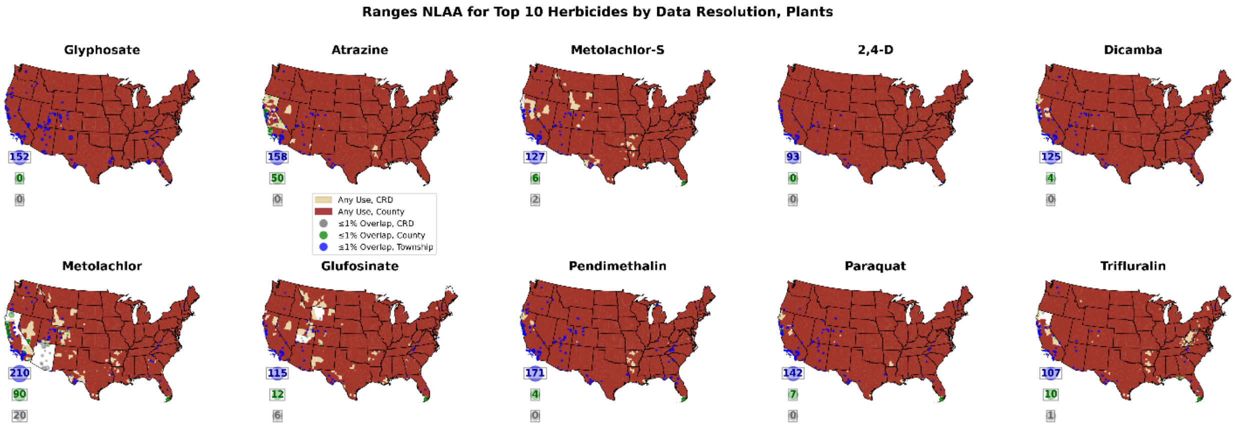

**Fig. S45.** Similar to Fig. S41, maps give the binary maximum usage footprints, locations and areas of ranges designated NLAA at each usage resolution, and total numbers of ranges NLAA at each resolution. Model predictions are used for township resolution outside CA; results are shown for the top 10 herbicides nationally, for plants.

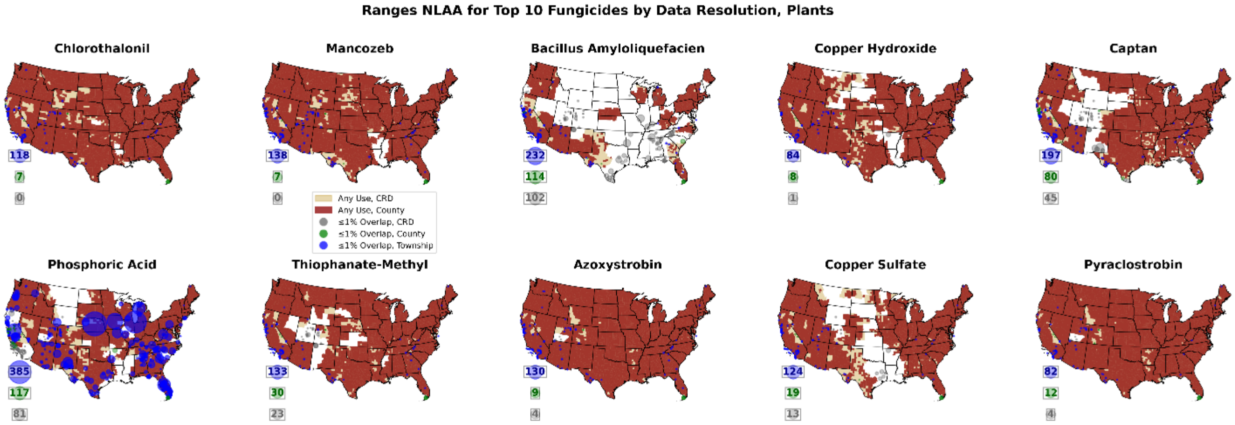

**Fig. S46.** Similar to Fig. S41, maps give the binary maximum usage footprints, locations and areas of ranges designated NLAA at each usage resolution, and total numbers of ranges NLAA at each resolution. Model predictions are used for township resolution outside CA; results are shown for the top 10 fungicides nationally, for plants.

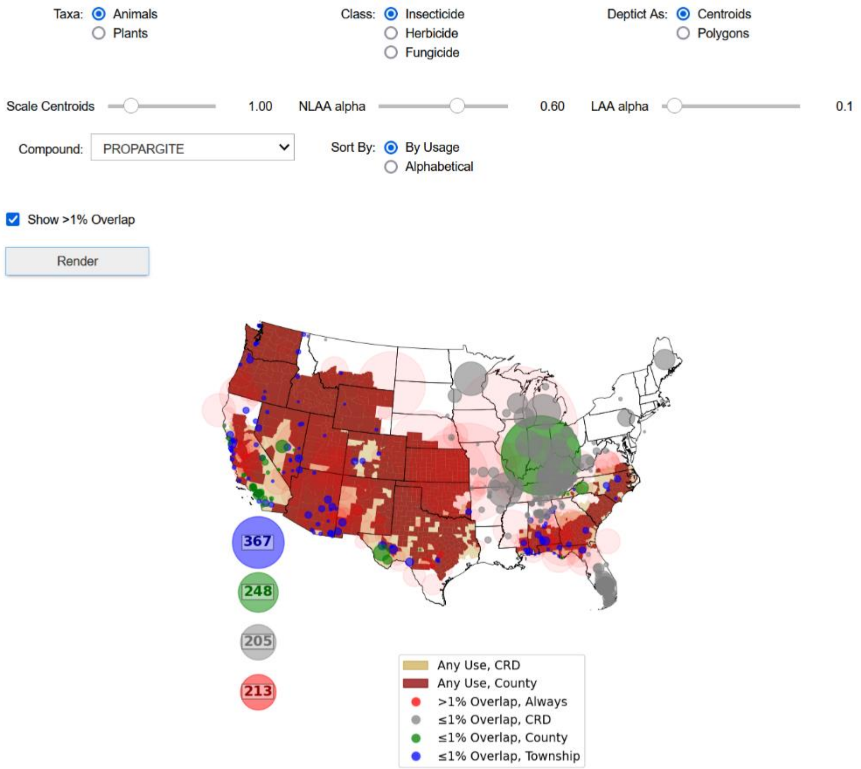

**Fig. S47.** Screenshot of Jupyter Notebook app to visualize overlap between pesticide usage footprint at different resolutions and species' ranges. Model predictions are used for township scale outside of CA. Screenshot has been cropped for clarity.

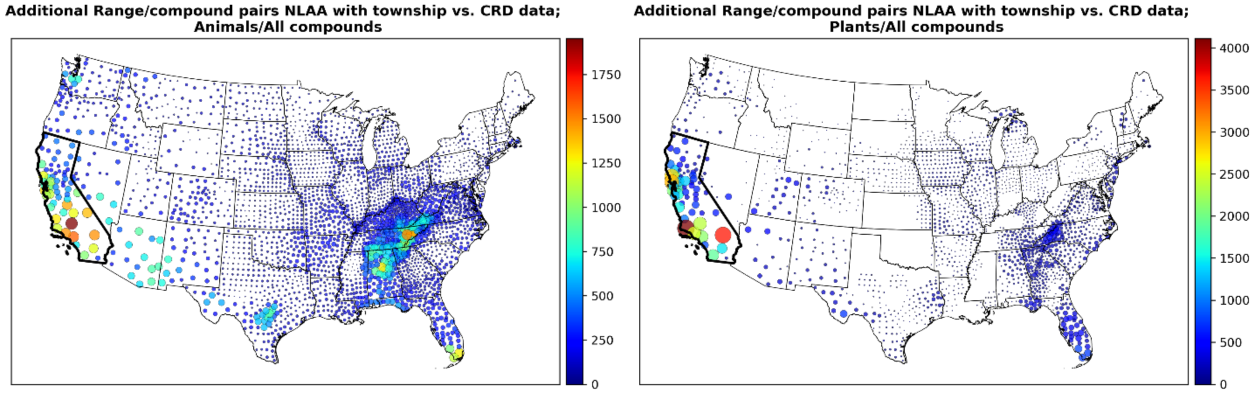

**Fig. S48.** The number of range/pesticide pairs predicted to switch from an LAA to NLAA classification if township-level data is available compared to a baseline where only CRD-level data is available, for each county, visualized as a bubble plot. The left panel gives results for animals, while the right panel shows results for plants. California is highlighted to emphasize that, for species restricted strictly to CA, predictions are determined using direct overlap calculations, while the classification model described in the text is used outside CA. This model is also used for ranges that overlap CA but are not restricted solely to that state.

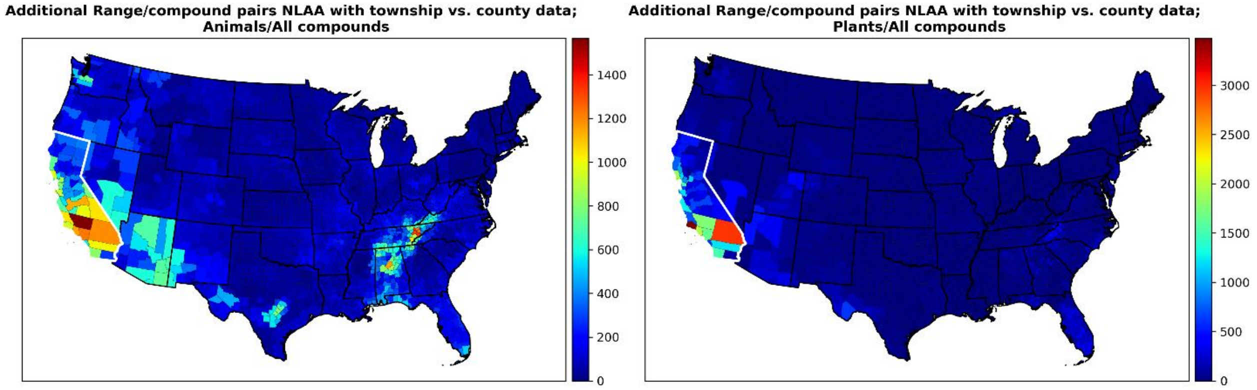

**Fig. S49.** The number of range/pesticide pairs predicted to switch from an LAA to NLAA classification if township-level data is available compared to a baseline where county-level data is available, for each county. The left panel gives results for animals, while the right panel shows results for plants. California is highlighted to emphasize that, for species restricted strictly to CA, predictions are determined using direct overlap calculations, while the classification model described in the text is used outside CA. This model is also used for ranges that overlap CA but are not restricted solely to that state.

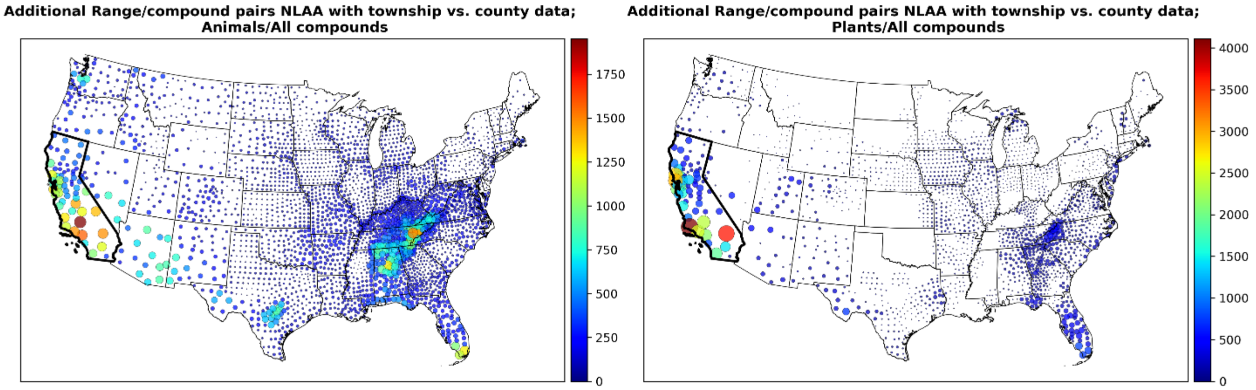

**Fig. S50.** Similar to Fig. S49, these plots give the number of range/pesticide pairs predicted to switch from an LAA to NLAA classification if township-level data is available compared to a baseline where county-level data is available, for each county, but visualized as a bubble plot. The left panel gives results for animals, while the right panel shows results for plants.

### 8. Priority States

Given the uneven geographic concentration of likely benefit to township usage data (compared to a baseline where only CRD usage data is available), we explored which individual states should be prioritized. For each state, we determined:
1. The number of additional ranges predicted switch to NLAA if township-level data was available for that state alone (and CA, given that such data is already available).
2. The number of additional ranges predicted to switch to NLAA if this was the *only* state for which township usage data was *not* available. That is, a "leave-one-out" analysis, meant to identify states that may contain many ranges that overlap with other states.

Both approaches identified largely the same subset of important states, with the most notable disparities seen in the Appalachian states of Mississippi, Georgia, Kentucky, Virginia, West Virginia, and North Carolina, where the leave-one-out approach indicated relatively greater importance.

Results are summarized in Figs. S51 and S52, which gives both the maps and bar graphs of the top states identified in each analysis. States in common in the top 10 are indicated by a solid bar, while those that disagree are indicated by a hatched bar. Figs. S53 and S54 give the priority states when comparing the benefit of township resolution usage data to county resolution data. Overall, the priority states are very similar as those identified under a CRD usage baseline.

We also determined how many range/compound pairings are predicted to switch from LAA to NLAA as a function of how many states (in addition to CA) have township resolution usage data. These states were added in order of priority determined either by the single state or leave-one-out method described above, and the results are summarized in Fig. 4 of the main text.

Finally, as described in the main text we identified 11 priority states other than California under a second leave-one-out analysis: FL, AL, TX, TN, NV, AZ, UT, GA, NC, OR, and WA. Fig. S55 plots these 11 states and indicates their relative priority.

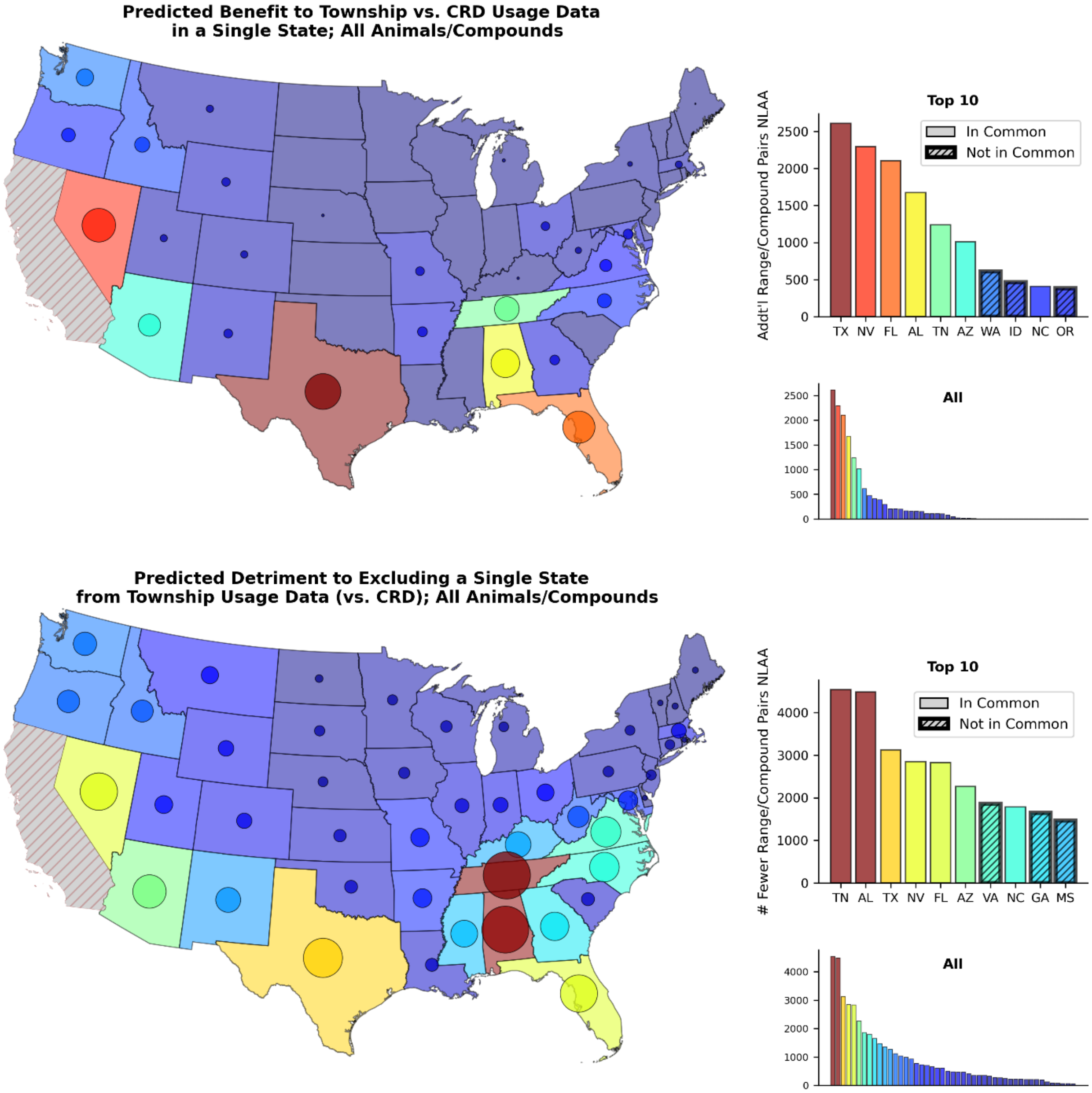

**Fig. S51.** The top panel gives the number of range/compound pairings expected to benefit from township-level usage data (relative to CRD data) when each state is the only state with such data. Conversely, the bottom panel gives the number of pairings that would fail to be classified as NLAA if township-level usage data was missing for a single state. The top 10 states are also indicated in the bar graphs, with states common to the two top 10 lists indicated with solid bars, and states not in common shown with hatched bars. The other set of bar graphs give results for all 48 states, but these are not labelled and meant only to show the shape of the distribution. Figure gives results for animal ranges and all pesticides.

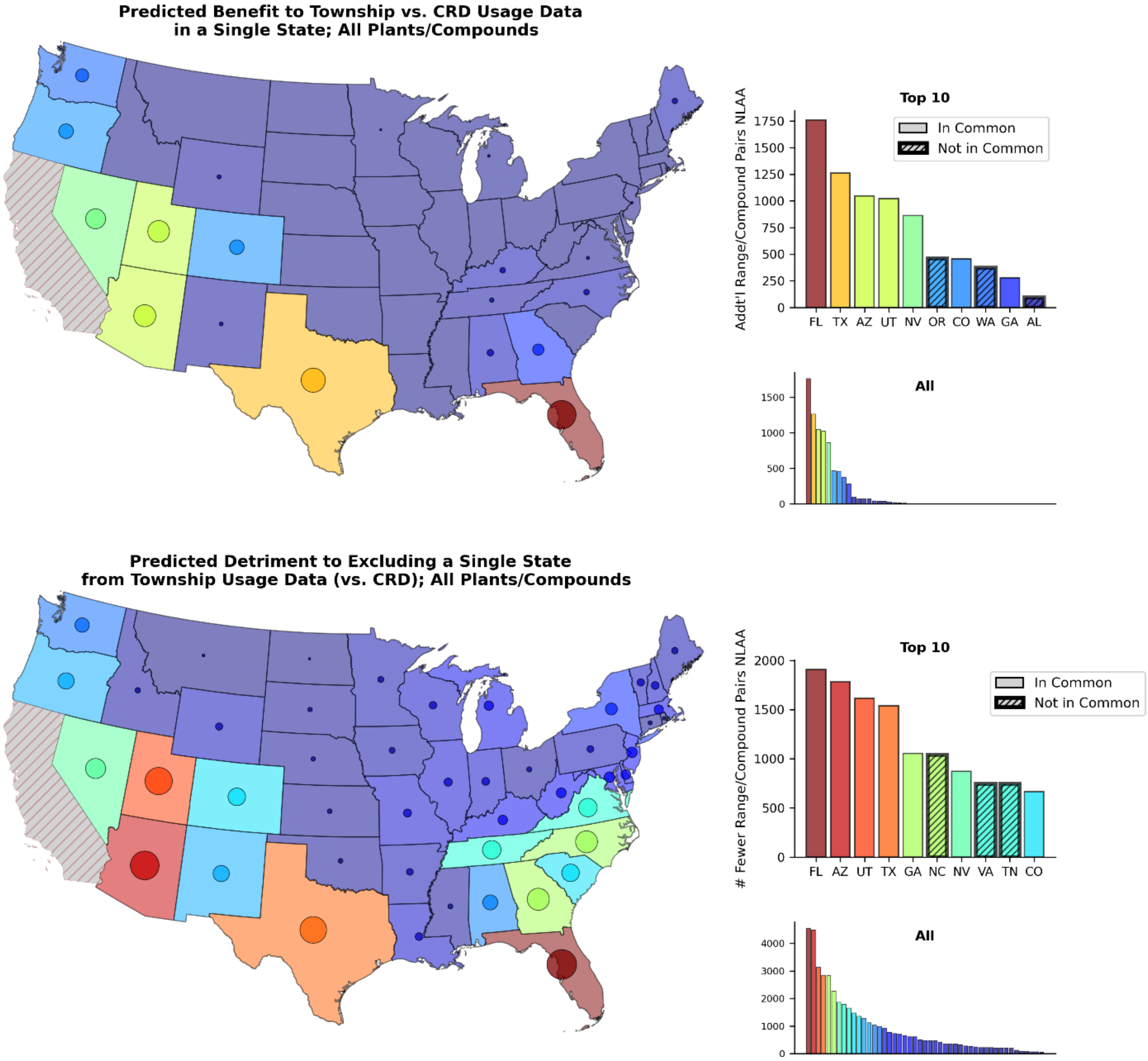

**Fig. S52.** Similar to Fig. S51, but for plant ranges and all pesticides. The top panel gives the number of range/compound pairings expected to benefit from township-level usage data (relative to CRD data) when each state is the only state with such data. Conversely, the bottom panel gives the number of pairings that would fail to be classified as NLAA if township-level usage data was missing for a single state. The top 10 states are also indicated in the bar graphs, with states common to the two top 10 lists indicated with solid bars, and states not in common shown with hatched bars. The other set of bar graphs give results for all 48 states, but these are not labelled and meant only to show the shape of the distribution.

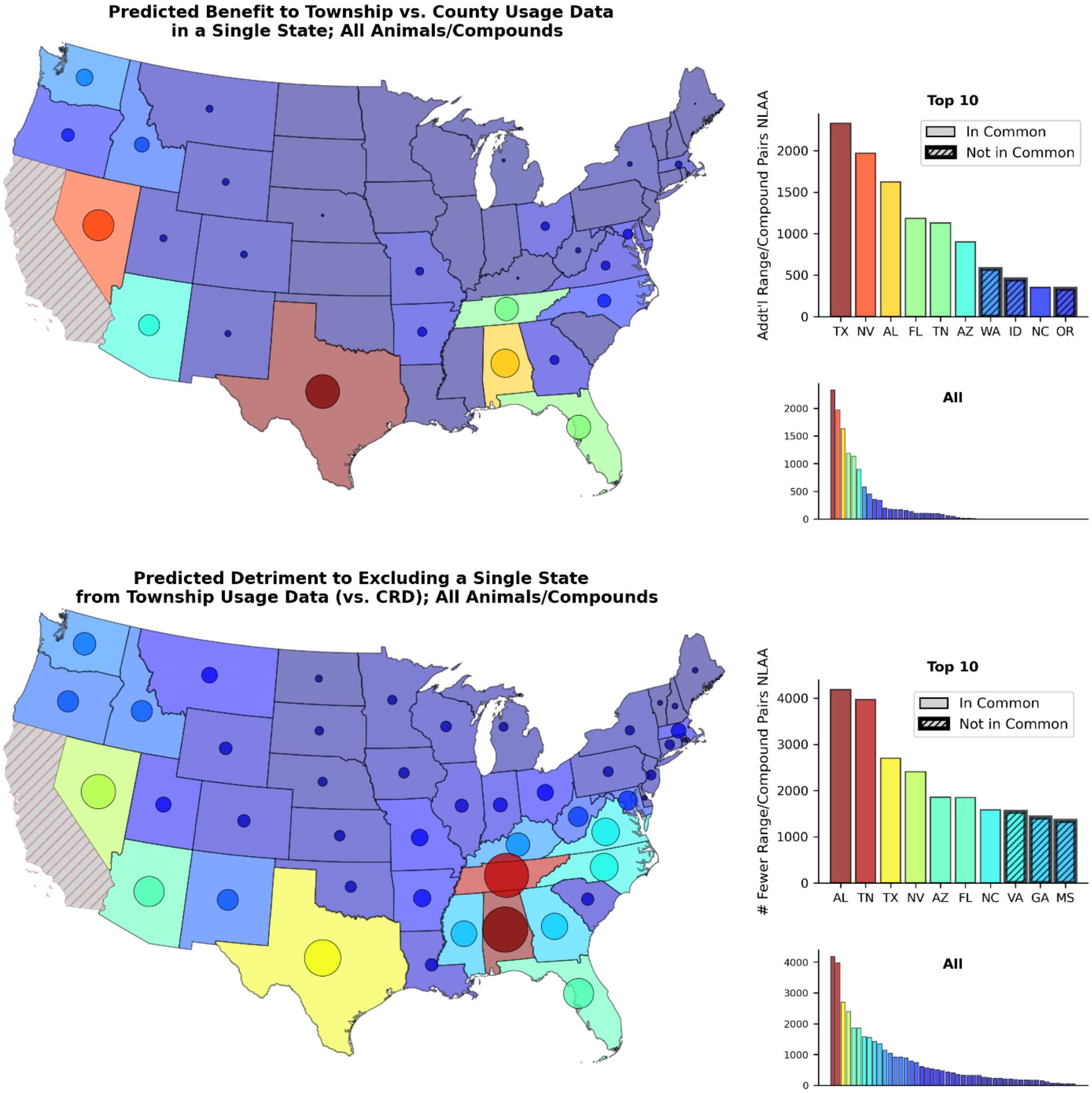

**Fig. S53.** Similar to Fig. S51 (and presenting result for animal ranges), but the benefit to township data is relative to a baseline where county usage data is available.

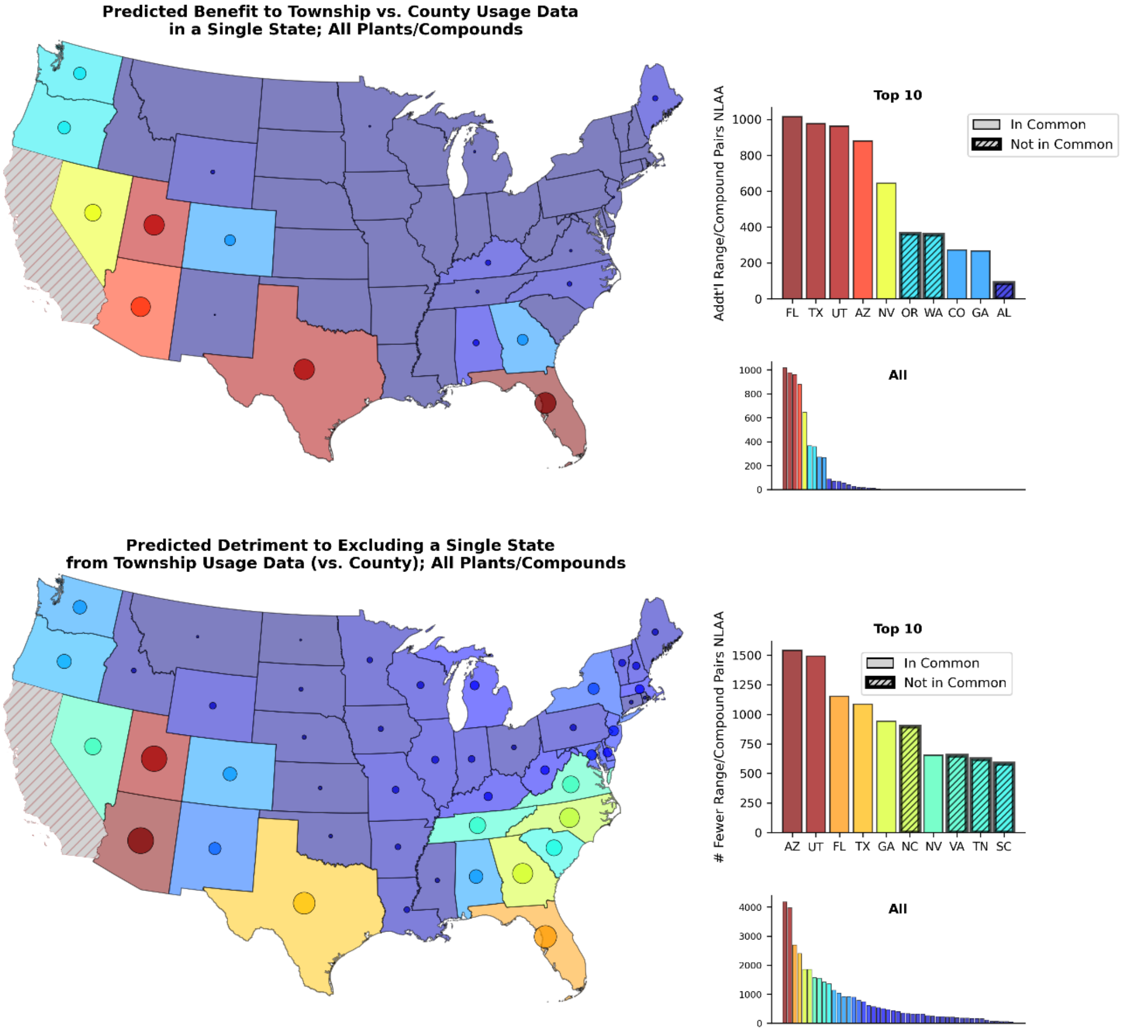

**Fig. S54.** Similar to Fig. S52(and presenting result for plant ranges) , but the benefit to township data is relative to a baseline where county usage data is available.

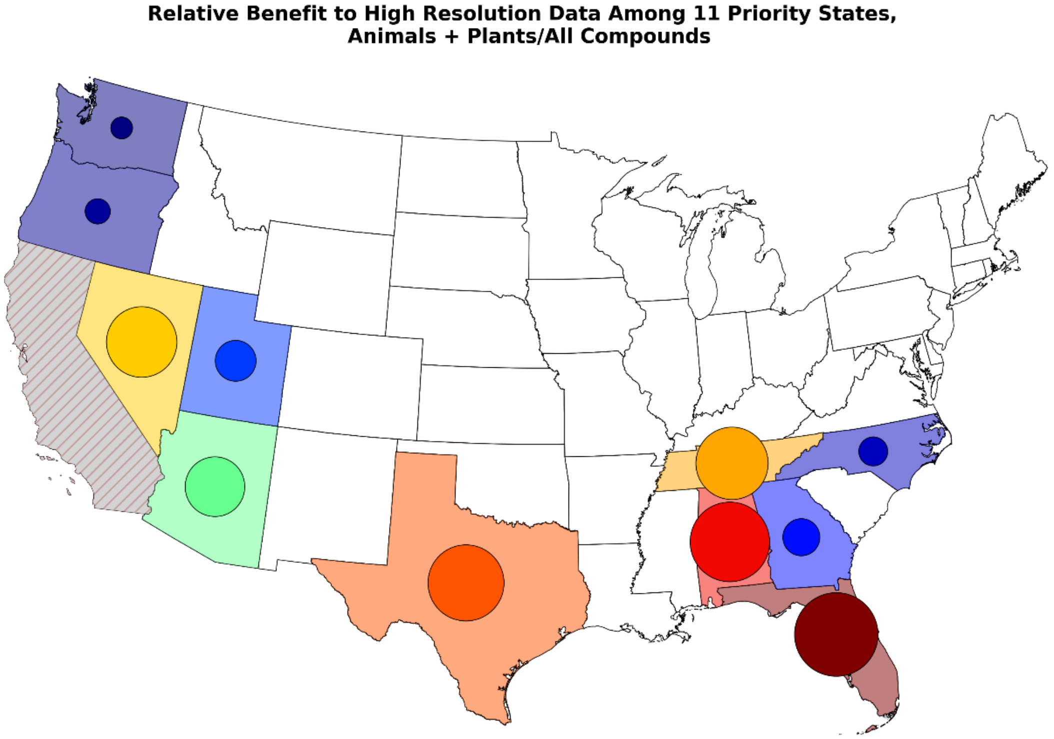

**Fig. S55.** The top 11 states (outside CA) identified as having the highest priority for high resolution data collection. Relative priority is indicated by the bubbles, with bubble size proportional to the number of pairings that would fail to be classified as NLAA if township-level usage data was missing for a single state.

#### Code to reproduce figures

Processed data and Python code (as Jupyter notebooks) to reproduce all figures is available on the Dryad repository (Eikenberry et al., 2022).

### 9. Overview of the EPA Biological Evaluation Methods for Spatial Analysis

The US Environmental Protection Agency (EPA) is responsible for assessing potential risk to ESA listed species. Under the current revised method, the EPA generates a Biological Evaluation for each species that designates a potential level of risk (EPA, 2021). Then the Services (US Fish and Wildlife Service and the National Marine Fisheries Service) review the designations that suggest potential risk and either concur or engage in a formal consultation to determine potential for jeopardy to the species (Fig. S56). Here we provide a summary of the detailed methods provided in the Biological Evaluation for carbaryl (EPA, 2021).

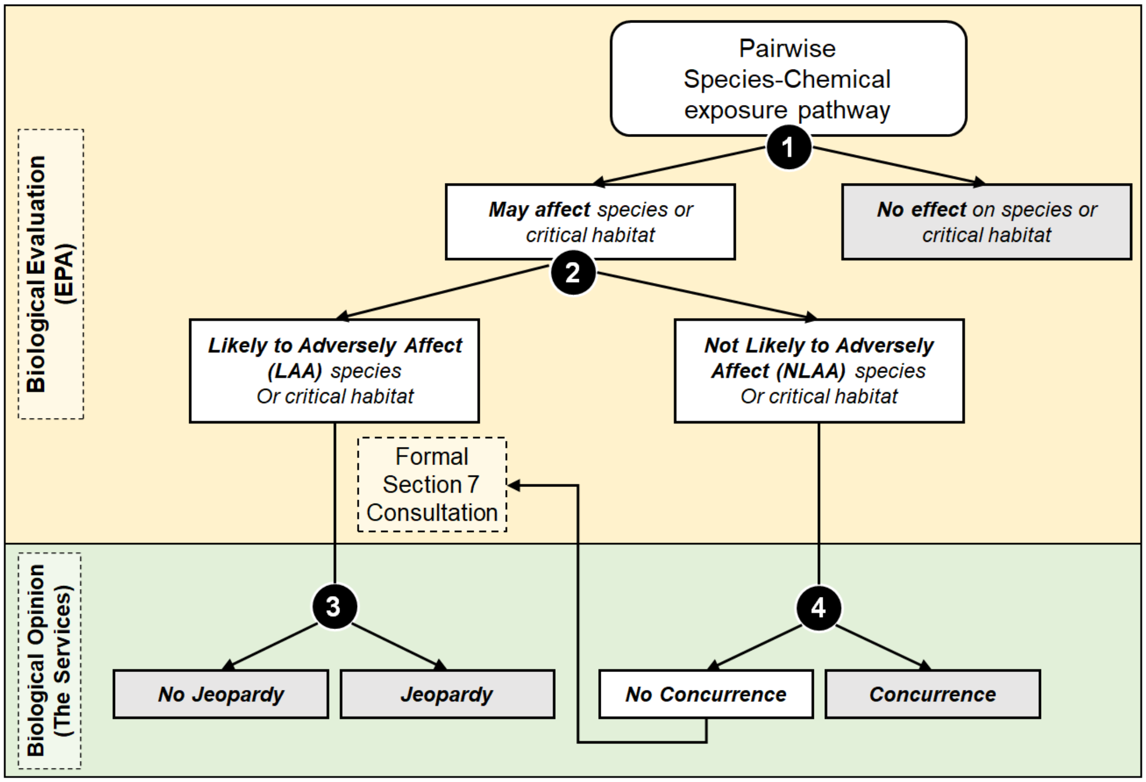

**Fig. S56.** The three-step process in the unified interagency framework for national level risk assessments for pesticides. The first two steps make up the EPA's Biological Evaluation and the third step is The Services Biological Opinion.

There are three designations a species can receive in the EPA method for Biological Evaluation. If a species is designated as 'No Effect' then no review from the Fish and Wildlife Service is necessary. This designation is given if there is no overlap between *potential* pesticide use areas—the use footprint—and species range or critical habitat area.

A designation of 'Not Likely to Adversely Affect' (NLAA) is given if there is less than 1% overlap between the action area and the species range or critical habitat. Based on conservative assumptions (i.e., assume maximum application and upper bound of overlap), if the overlap between the species range or critical habitat and the usage footprint suggests that less than one individual from the species will be exposed, assuming equal distribution of individuals throughout their range, then a designation of NLAA is also given. Otherwise, EPA uses a weight of evidence approach that considers three different usage levels, three different usage distributions, and five different scenarios for overlap to determine the potential risk of pesticide exposure to each species and critical habitat (described below). If the EPA determines the risk of exposure is low, then a designation of NLAA can be given.

A designation of 'Likely to be Adversely Affected' (LAA) is given if there is >1% of overlap between a species range and areas of actual pesticide use. In this instance, FWS must perform a formal Section 7 consultation.

Below, we detail the steps of the spatial analysis that determines this overlap as described for the biological assessment of Carbaryl (Fig. S57). We designed our VOI approach to test the critical aspects of this spatial analysis.

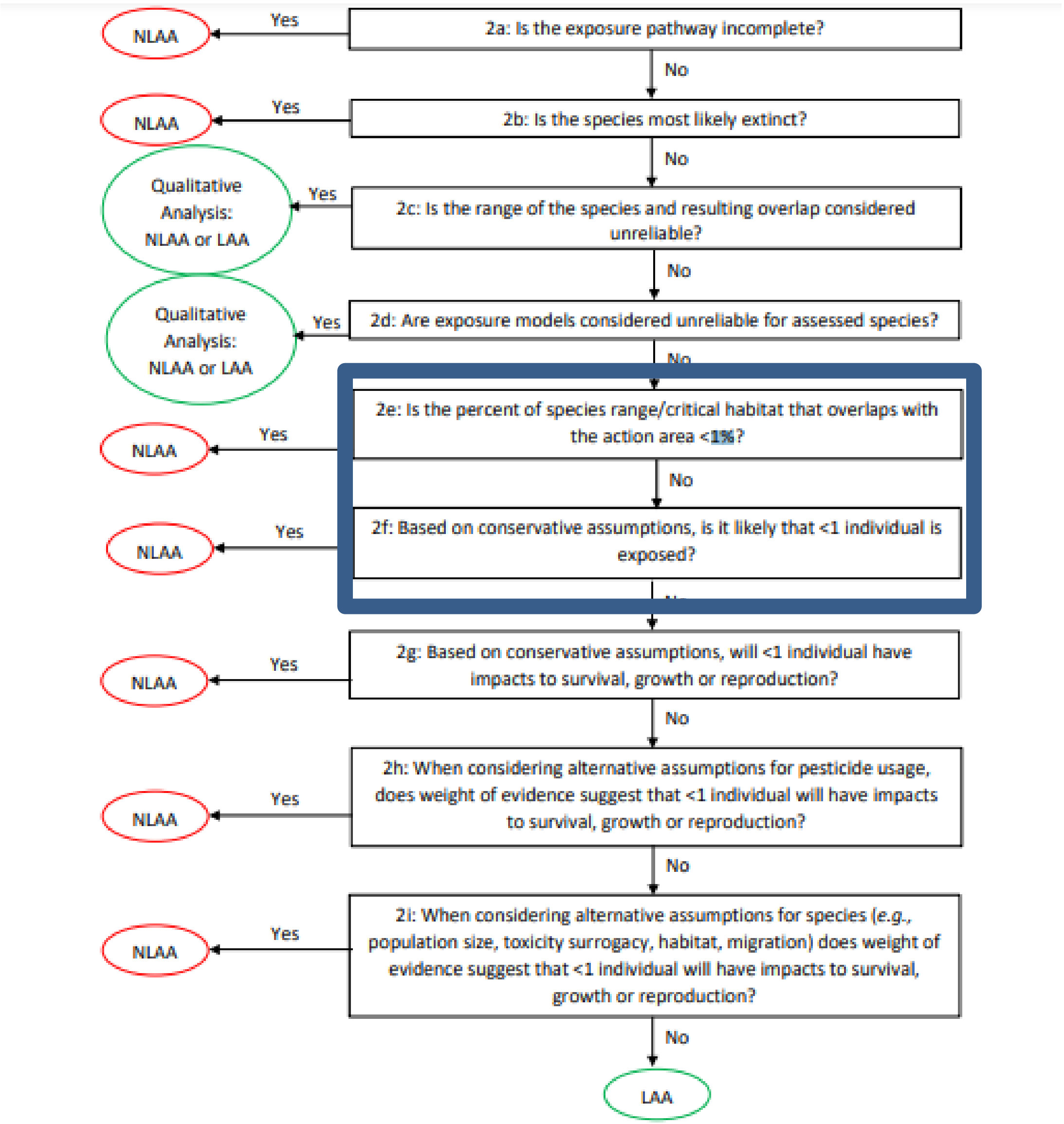

**Fig. S57.** Adapted from Figure 4 in the EPA's Revised Method for National Level Listed Species Biological Evaluations of Conventional Pesticides (U.S. EPA, 2021). Summary of how the spatial analysis can lead to an NLAA determination with boxes around the steps where we evaluated the benefits of increased spatial resolution for pesticide usage data. These steps are only taken once a 'No effect' determination is excluded.

*Steps of EPA revised methods spatial analysis:*

1. Create the action area
   a. Create the agricultural use data layer
   b. Create non-agricultural use data layer
   c. Compile action area with drift zones
2. Collect species range and critical habitat data
   a. Master species list
   b. Species ranges and critical habitat areas
3. Create usage footprint
   a. Calculate percent crop treated
   b. Compile usage data with use footprints
   c. Calculate usage drift

4. Calculate pesticide-species range/critical habitat overlap
5. Results of analysis

**Step 1. Create the action area**

In our analysis, we do not calculate an action area, but instead assume that the pesticides can be used anywhere, making all of California or the whole contiguous U.S. the action areas for our respective analyses. As a result, the only possible designations for species in our analyses were LAA or NLAA. This is more conservative than the EPA approach that limits potential overlap to the action area, as described below.

**A. Create the agricultural use data layers (UDLs; summarized from Appendix 1-6 in EPA, 2021)**

EPA uses the Cropland Data Layer (CDL), produced by the U.S. Department of Agriculture, to create maps of specific agricultural crops throughout the contiguous U.S., as this is the best data available. The CDL has over 100 cultivated classes that EPA groups into 13 general classes for this analysis (see Table 1 in Appendix 1-6 of (EPA, 2021)) to reduce errors of omission and commission between similar crop categories. The 13 classes were informed by the U.S. Geological Survey (Baker and Capel, 2011) and the Generic Endangered Species Task Force (Amos et al, 2010).

Once these classes are developed, the data is temporally aggregated. EPA aggregates the most recent five years of available CDL data to capture temporal heterogeneity in crop cover in the U.S. The resulting crop layers, which captures both the crop classes and temporal variation are referred to as Use Data Layers (UDLs)—the areas where a pesticide, herbicide, or fungicide may be used.

The agricultural classes are further refined by comparing the UDLs to the county level National Agricultural Statistics Service (NASS) 2012 Census of Agriculture (CoA) data (See attachment 1-3 in *Final National Level Listed Species Biological Evaluation for Carbaryl* for filtering process to create NASS data layer). If UDL acreage is less than NASS acreage, the raster is expanded in 1pixel iterations in the cultivated are until the NASS acreage value is reached to avoid buffering into any non-agricultural landcover types. This method aims to reduce landcover mapping errors by adjusting the extent of each category to the CoA values.

The CDL is only available for the contiguous U.S. (ConUS), so other data sources are used for states and US territories outside of ConUS (referred to as NL48). Methods for the NL48 are available in Appendix 1-6.

Once the UDLs are created, every chemical assessment begins with cross-walking of registered uses for the given chemical with the landcover categories. Some chemicals specify geographic restrictions for a given use as well (e.g., it can only be used in each state). These geographic restrictions are extracted from the appropriate UDL before it is aggregated with other chemical uses to generate the chemical's action area.

**B. Create the non-agricultural use data layers**

Chemicals also have non-agricultural label uses, which include a wide range of landcover and land use categories. As for agricultural land uses, each of these uses is cross walked with the available landcover data. Depending on data availability, the 2011 National Land Cover Dataset (NLCD) is used first to represent non-agricultural uses, but when it is not available, the NOAA C-CAP dataset, GAP Protected Areas Database, LandFire and NAVTEQ are used. A complete crosswalk for the non-agricultural uses is provided in Table 2 in Appendix 1-6 of EPA (2021).

#### C. Compile the action area with drift zones

The action area is the aggregation of all the agricultural and non-agricultural UDLs into a single layer. This composite layer is then buffered out based on the greatest drift potential for the chemical. Then the composite drift area is refined for individual species based on the species' range and the use overlap assuming the maximum buffer for the chemical.
.

#### Step 2. Collect species range and critical habitat data (summarized from Appendix 1-7 in EPA, 2021)

#### A. Master Species list

Species subject to Section 7 under the ESA during the registration time are obtained from the US Fish and Wildlife Threatened and Endangered Species System (TESS). For species under the jurisdiction of the National Marine Fisheries Service (NMFS), TESS data is supplemented with information from the NMFS website. If data between these sources' conflict, then EPA uses the information from the NMFS website. We used the same species list for our starting analysis, though we ultimately only included species in the contiguous U.S. and excluded three plants with erroneous range maps.

#### B. Species location

The FWS ECOS Portal (http://ecos.fws.gov) houses the best available data on species' ranges and critical habitats. Data for species under NMFS jurisdiction that is not available on the ECOS website is taken from the NMFS website. NMFS scientists were contacted directly if range data was not available on either website.

The co-occurrence analyses are completed using the ESRI ArcGIS Union Toolbox. This tool combines species' range data into non-overlapping 'zones'. This allows for the overlap analysis to be run only one, because all species associated with each zone are captured (Figure S58).

We also pulled species' range maps from the FWS ECOS Portal for our analyses. However, our analysis differs in that we only considered ranges, not critical habitats. Additionally, we performed a separate analysis for each range, and considered distinct ranges for the same species as separate entities in our analysis.

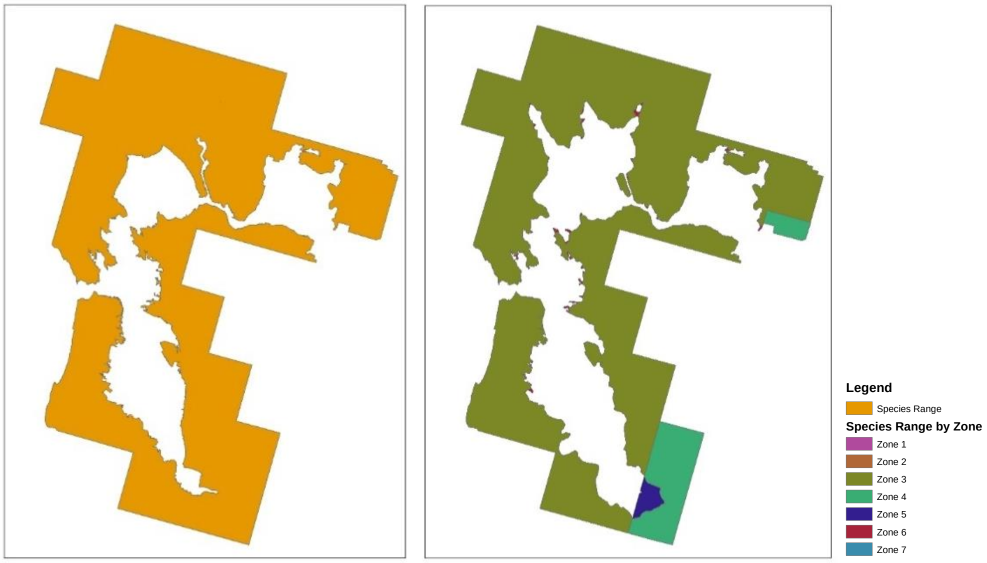

**Fig. S58.** Figure 3 from Appendix 1-7 (EPA, 2021). Example of species range in zones used as the input for the co-occurrence analysis.

**Step 3. Create usage footprints (summarized from Appendix 1-7 for EPA, 2021)**

The goal of the third step is to determine the area within each state that is treated with a pesticide (pesticide usage) and the extent of its overlap with species range or critical habitat (Fig. S59 gives overview). This is accomplished in three steps: 1) calculate the percent of crop treated, 2) combine usage data with the potential use sites, and 3) calculate percent overlap for each species and critical habitat.

Our methods differ from this in a few ways. First, this method begins with an action area, smaller than the area of the state. We assume the whole state is the action area. Second, the EPA method is only partially spatially explicit, with three different assumptions for pesticide usage within the action area considered at the state scale, while our analysis is wholly spatially explicit. The CRD scale in our analysis however, is closest to the analysis done by EPA.

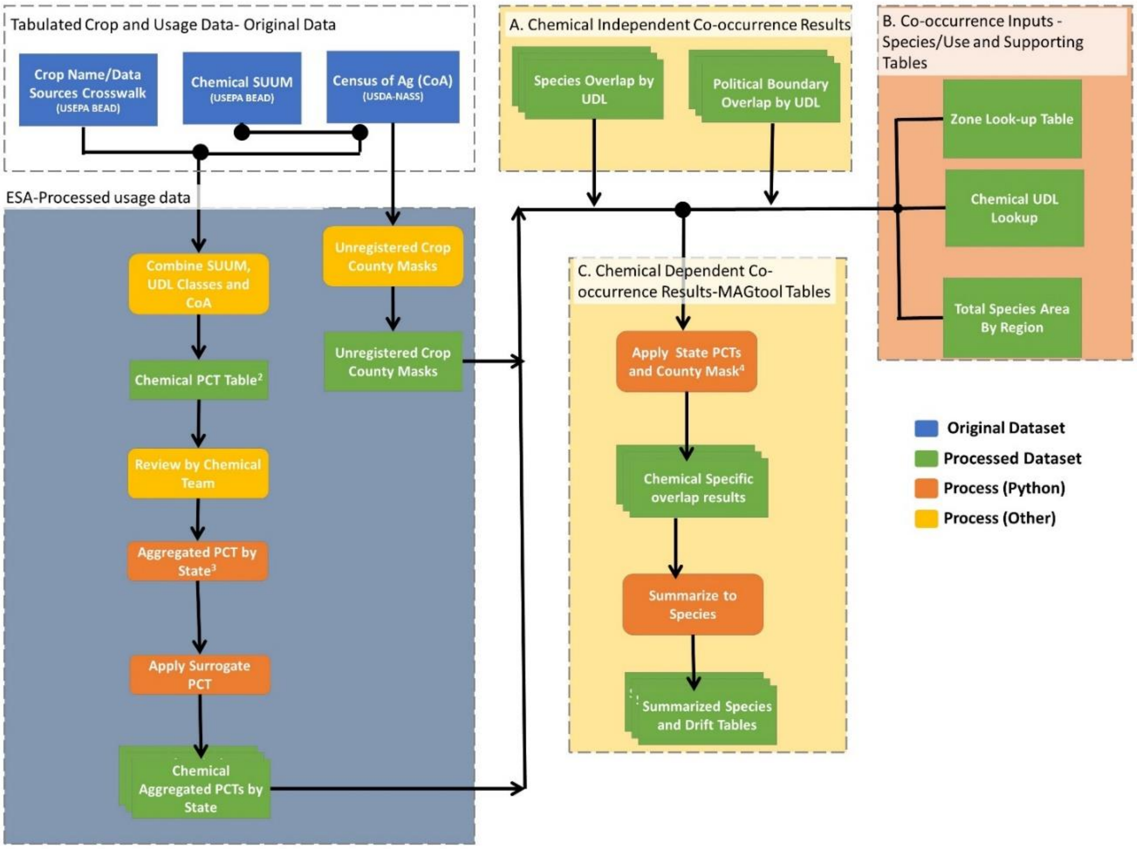

**Fig. S59.** Figure 1 in Appendix 1-7 in EPA, 2021. This Flow chart shows the how usage is applied to the UDL and the compared with species range to create the species' and UDLs co-occurrence results.

**A. Calculate the acres treated**

Acres treated is calculated by first identifying the percent of crops treated (PCT) and multiplying the UDL area by the PCT to identifying the total acres treated. Percent of crop treated (PCT) is the percent of the acres grown for a crop that are treated. These are presented for each UDL using minimum, maximum, and average PCT over the five-year observation period to address temporal variability and spatial uncertainty in pesticide usage (described below).

*Average PCT*: The total number of pounds applied is divided by ½ of the maximum application rate on the label for the given UDL rate plus the minimum rate. The use of ½ of the maximum application was selected because average reported applications are approximately half of the maximum rate. If a use specific 'typical' rate is identified, this can be used instead of the average rate. Both the typical and average rates are more conservative than the maximum PCT.

*Minimum PCT*: Assume pesticide is universally applied at the minimum application rate on the label for the given UDL. Assuming the minimum application rate per acre corresponds to the maximum treatment area possible for a given amount of chemical.

*Maximum PCT*: Assume pesticide is universally applied at the maximum application rate on the label for the given UDL. Assuming a maximum application rate per acre corresponds to the minimum treatment area possible for a given amount of a chemical.

As we use spatially explicit pesticide usage data, we only use one estimate for the acres treated, and we assume equal distribution of the pesticide within the highest resolution spatial unit for a given analysis. For example, if 10 kg is used in a given township over a year, we assume that these 10 kgs are used with an equal distribution across the township.

*Calculating acres treated for agricultural usage for ConUS*:

Percent of crop treated are reported at the state level because there are uncertainties in extrapolating from national level usage data to regional and state level ranges of protected species. National level data does not distinguish if there are areas of a species' range where usage is greater or less than the average national usage.

The pesticide usage data (agriculture and non-agriculture) are obtained from both public and private (proprietary) sources: Kynetec USA, Inc. AgroTrak study, USDA NASS, CADPR PUR data and non-agricultural market research data (NMRD). Only PUR is all usage and not survey data. The presented usage data are averaged over the number of years of available survey data during the most recent five years of available data, based on sampling frequency (five years for Kynetec and CADPR, and 1-2 years for NASS and NMRD), regardless of whether usage is observed in each surveyed year. The presented data may thus underestimate the maximum yearly usage. Kynetec is the primary data source as it collected annually and tends to provide the most robust usage data among the available data sources. NASS data are used for crops which are not surveyed by Kynetec data. For crops with less than 80% California production, Kynetec is the primary source of usage data. The presented data may not be a reliable indicator of the variability in usage between individual years.

PCT data are available for specific crops and states. The method discussed below is applied separately to the average, minimum and maximum annual PCT data to quantify the overlap of species range and exposure areas, while accounting for variability in usage over time.

Crops reported in the Science Information and Analysis Branch (SIAB) Use and Usage Matrix (SUUM; APPENDIX 1-4 of EPA, 2021) are crossed with the categories used for the UDLs. For categories where a landcover UDL and SUUM both represent a single crop (e.g., soybean), the PCT data available for a given state are applied directly to the UDL in that state to calculate the acres treated (acres treated = acres grown x PCT). If the PCT is not available for a specific state/crop combination, a surrogate PCT is applied using the process described below.

When a UDL or SUUM covers multiple crops (e.g., vegetables), an aggregate PCT is calculated. The aggregate PCT for a state is calculated by summing all the acres treated for every crop in the UDL and diving this by the total acres grown for all crops (Eq. 1).

The PCT and the total acres of each crop are found in the SUUM for each state/crop combination with reported usage. Acreage in the SUUM can come from market research data, USDA's National Agricultural Statistics Service (NASS), California's Pesticide Use Reporting (PUR), and other sources. If the state/crop combination does not have reported usage in the SUUM, data from the 2012 Census of Agriculture is used (USDA-NASS, 2012).

Therefore, there are three ways the acres treated can be calculated:

- The Census of Agriculture crop/state combination is found in the SUUM, and the acres grown and crop specific PCT from the SUUM are used.
- The Census of Agriculture crop is registered but there is no state usage information in the SUUM. Here, the CoA data on acres grown for the state are used and a surrogate for the crop specific PCT is assigned (method described below).
- The Census of Agriculture crop is not a registered use. In this scenario, the acres grown come from Census of Agriculture and a PCT of 0 is used

**Eq.1**

$$PCT_{tot-j} = \frac{Acres\ treated}{Acres\ grown} = \frac{\sum_{i=1}^{n}(PCT_i * G_i)}{\sum_{i=1}^{n} G_i}$$

Where: i = crop (within land cover class j) that is surveyed in state, j = land cover class (e.g., vegetables and ground fruit), n = # of crops (within land cover class j) with acres grown in state, $PCT_j$ = % crop treated of crop i (from extended SUUM), $PCT_{tot-j}$= aggregated PCT (for land cover class j in state), $G_i$= acres of crop i grown (in state) (from extended SUUM) (Eq. 1 can be found in Appendix 1-7, EPA, 2021).

Once the total PCT is calculated for UDLs with multiple crops, the total acres treated is calculated by multiplying the aggregated PCT by the area of the state's UDL. The total area of the UDL for the state only includes those counties with at least one registered use as reported in the Census of Agriculture.

This approach is conservative, as it does not account for multiple applications to the same fields in a single year and assumes all acres treated are independent. Therefore, if the available usage data includes sites where multiple applications per year occurred, then the extent of the treated acres will be an overestimate.

*Calculating acres treated for non-agricultural usage for conUS:*

*Developed land:* Create a minimum, maximum and average number of acres treated and use these to inform likelihood of exposure. Each of these values are divided by the total number of acres in the potential use site UDL for developed land to derive the average, minimum and maximum national level PCT for the developed landcover. PCTs are applied to the area of the developed UDL found in each state to estimate the treated acres of the developed landcover for the state.

*Open space developed:* The available usage information for open space developed is regional and includes all use sites under the heading Ornamental Lawn & Turf; applied by Lawn Care Operators, Applied by Landscape Contractors, In Institutional Turf Facilities, Golf Course and Ornamental Sod Farms (Turf). Maximum, average and minimum treated acres for open space develop are estimated for each region. The number of estimated treated acres is divided by the number of potential use site acres in the region to generate each regional level**.** The regional PCT are applied to the open space developed UDL for each state in the region.

*Right of way (ROW):* ROW usage includes pesticides applied to roadways, electrical areas, railroads etc. Maximum, average and minimum treated acres for nurseries are estimated using the same methods described for developed land. Insecticide usage is low on electrical and railroad ROW and is not surveyed. In this approach, the conservative assumption of the acres treated is assumed to help offset the uncertainty associated with a lack of survey data for non-roadway rights of way. The total number of treated acres is distributed uniformly throughout the entire US to all potential use sites located in the US. Therefore, PCT is calculated by dividing each of the estimated treated acres by the total acres found in the ConUS right of way landcover for minimum, maximum and average estimated treated acres**.** These national PCT values are applied to right of way landcover for each state.

*Forest trees:* Usage data on forest tree application come from USDA's forestry service. These data include acres treated of all lands in the 50 states that fall under the responsibility of the Forestry Service. The usage information collected from USDA's forestry service is used as a surrogate for applications to forest trees not made by the Forest Service. The regional PCT are applied to the forest trees landcover for each state in the forest service region; values less than 1 are rounded to 1. For un-surveyed regions the PCT surrogacy method describe below is applied.

*Rangeland*: Usage data on rangeland come from USDA APHIS. The usage information collected from USDA's APHIS is used as a surrogate for applications to rangeland not made by USDA. The state PCT are applied to the rangeland landcover by state; values less than 1 are rounded to 1. For un-surveyed states, the surrogacy method describe below is applied.

*Nurseries*: The available usage information for nurseries is regional and includes the Ornamentals (Unspecified): Covers Trees and Plants, Woody Shrubs and Vines grown in Nurseries. Maximum, average and minimum treated acres for nurseries are estimated. The number of estimated treated acres is divided by the number of potential use site acres from the nurseries landcover in the region (multiple states) to generate each regional level PCT.

*Applying surrogate usage data*: Some uses are not surveyed for usage and others are only surveyed in some states. The surrogacy approach uses the best available data to estimate the likely extent of treated area when usage data are not available. Surrogate data are ideally assigned using crops within the same landcover (UDL) and then using data from the same crop but different locations. If the first two options are not available, then a conservative approach is employed where the greatest extent of usage on any crop-state combination is used. After applying the surrogacy method all UDL/state combinations will have an associated aggregated PCT. In cases where a species has potential risk concerns, a Weight of Evidence analysis is conducted prior to making the determination and the impact of the surrogacy assumptions are considered.

**B. Combining usage data (i.e., acres treated) with potential use sites.**

As part of the Weight of Evidence approach, three measurements are calculated to estimate how the acres treated can be attributed to potential use sites within the species range: upper bound, uniform distribution and lower bound. In all three approaches the estimated treated area within the species range or critical habitat is used to calculate the direct overlap (i.e., the total treated area within the species range divided by the total area of the species range or critical habitat). As previously stated, our analysis assumes the entire contiguous U.S. is a potential use site and uses spatially explicit usage data for the co-occurrence analysis (described below).

The upper bound approach assumes all the treated acres within a state occur in the species range (or critical habitat). If the number of treated acres in a state is greater than the number of acres in the UDL overlapping the species range, it is assumed that the entire species range within that state that overlap with the UDL are treated (if a species range occurs partially outside the action area, the assumption is this area is not treated). If the number of treated acres is less than the number of acres in the species' range within the state, then the number of acres overlapping is assumed to be the number of treated acres in the state.

The uniform distribution approach assumes the treated acres are uniformly distributed throughout the state. Therefore, the aggregate PCT is applied directly to the acres of the UDL occurring within the species range to calculate the estimated treated acres.

The lower bound approach assumes that the treated acres are distributed outside of the species range as much as possible. If the number of treated acres is less than the number of acres outside of the species' range in the state, it is assumed that there is no direct overlap between species range and treated acres. If the number of treated acres is greater than the number of acres in the UDL outside of the species range, the number of treated acres in the species range is estimated as the remaining acres after all acres within the UDL and outside the species range are accounted for as treated.

When a species range spans multiple states, the uniform, upper, and lower bound approaches are applied to each state separately. Then the treated acres across all states the species inhabits are summed to calculate the number of treated acres overlapping with the whole species range (or critical habitat).

**C. Calculation of Composite Drift Layer Overlapping with Species Range or Critical Habitat**
When usage data are considered, the EPA accounts for a decrease in the extent of areas receiving spray drift because the treated area is smaller. Since the actual location of the treated acres within a state is unknown, specific areas are not buffered. Instead, a factor is applied to the composite drift area, previously described, based on a state aggregate PCT for all the uses. Different factors are applied to account for the distribution of theses acres under the three different distribution scenarios. For the upper bound scenario, no additional factor is applied to the aggregate PCT. For the uniform and minimum scenarios, the ratio of the number of treated acres calculated for the uniform or lower bound scenario to the upper bound scenario is applied to the PCT. This composite factor is used to scale the number of acres impacted by off-site drift and lowers the total predicted overlap with a species range (or critical habitat) due to drift.

Wind direction is also considered in these calculations. A wind direction scaling factor is applied, where the factor is scaled to 25% for each application allowed, to represent movement of a chemical off-site in only one direction (i.e., ¼ of a circle from where one application is made). To capture repeated sprays at one site, the factor is scaled by the number of possible applications. So, if 2 applications are allowed, a factor of 0.5 is used instead of 0.25.

The total number of acres in a species range or critical habitat potentially exposed due to spray drift is then divided by the total acres in the species range to determine the overlap area due to drift.

**Step 4. Determination of Overlap pesticide and Species Range or Critical Habitat.**

The co-occurrence analysis the extent to which a species range or critical habitat and UDL overlap. The overlap is equal to the number of treated acres within the species range plus the number of acres receiving spray drift, divided by the total number of acres in the species range.

Required data for conducting the co-occurrence analysis include a list of species, species location files, pesticide Use Data Layers (UDLs), usage data, and any additional supporting species life history information used to supplement the analysis. Information on the tools that EPA uses to conduct these analyses can be found in the appendices and attachments of EPA, 2021.

The five overlap scenarios used to estimate overlap are summarized below and discussed in more detail in the following section:

- Overlap Scenario 1: Spatial Co-occurrence of Species Location and Potential Use Sites
- Overlap Scenario 2: PCT Overlap (overlap with usage)
- Overlap Scenario 3:PCT and Redundancy (addresses duplicates in UDLs)
- Overlap Scenario 4:PCT, Redundancy, Off-site (considers life history and offsite only exposure)
- Overlap Scenario 5: PCT, Redundancy, Off-site, Beach (considers beach specialists)

Using the three different assumptions for distribution of treated acres within the species range (*i.e.,* upper bound, uniform distribution, and lower bound) and the three different assumptions about usage rates (*i.e.,* maximum, average, or minimum annual PCT), nine estimates of the overlap of the species range and the exposure area emerge for each overlap scenario.

**Overlap Scenario 1: Spatial Co-occurrence of Species Location and Potential Use Sites**
The first overlap scenario only considers where pesticides can legally be used (i.e., potential use site) and where species may occur based on their range, excluding consideration of actual pesticide usage or species life history. If there is no overlap between the potential use site with their drift buffer and the species range, then a 'no effect' designation can be given; however, this is very rare. As we assumed pesticides can be used anywhere within the country, we did not consider scenario 1.

**Overlap Scenario 2: Calculate overlap with PCT**
This method uses the upper, lower, and uniform distributions of pesticide usage, following the methods described above. After the treated acres for each state within the species range (or critical habitat) are calculated, the total treated acres for the species in each state are summed and divided by the total acres in the species' range to calculate the overlap. This is done for all UDLs and each of the aggregate PCTs; minimum, maximum, and average (Fig. S60). This scenario is closest the analysis we performed, if the UDL is assumed to be the whole state. However, this analysis is not spatially explicit, where as our pesticide usage maps are spatially explicit.

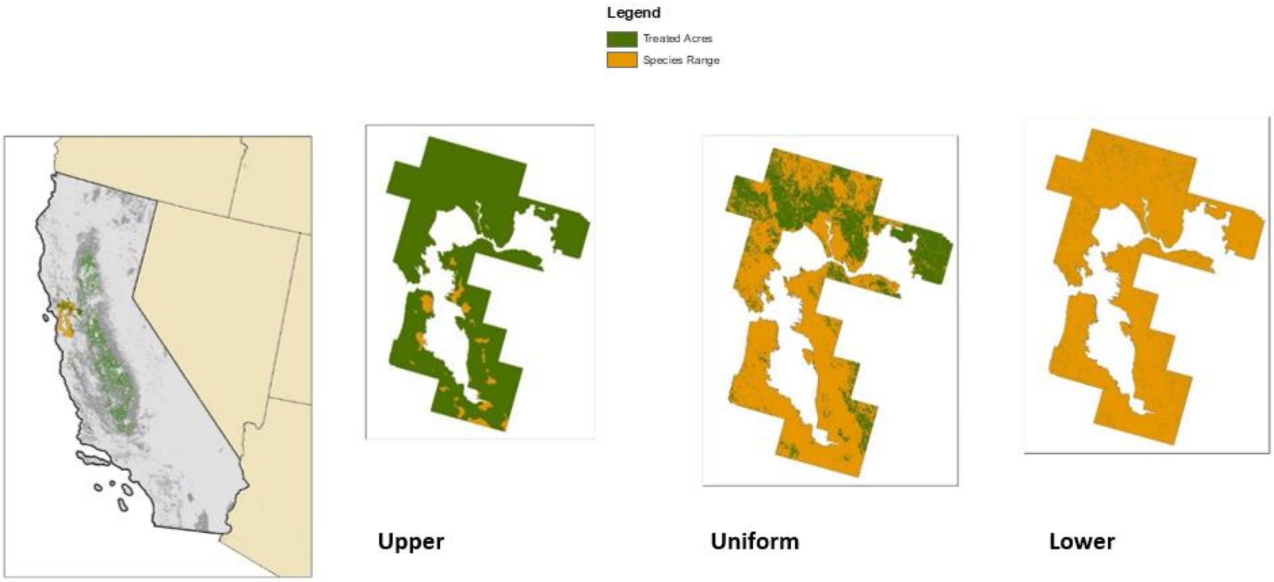

**Fig. S60.** This is Figure 4 of the Appendix 1-7 of EPA (2021). It provides an example of the three possible distributions of treated acres (upper, uniform and lower) for an aggregate PCT in a UDL.

**Scenario 3: Scaling for redundancy in the UDLs**
Many UDLs for a given chemical overlap, which causes the sum of the UDLs to be greater than the action area, and often greater than 100% of the species range. To account for this redundancy three factors are applied to results for individual UDLs: the composite factor, the agricultural factor, and the non-agricultural factor.

The composite factor accounts for redundancy between agricultural and non-agricultural uses. It is the sum of all agricultural UDLs and non-agricultural UDLs divided by the action area. If all uses are independent the sum of two composites would equal the action area, and the factor would be 1. Each individual UDL is divided by this composite factor.

The agricultural factor accounts for redundancy between agricultural UDLs. It is the sum of all the agricultural UDLs divided by the agricultural composite. Similarly, the non-agricultural factor accounts for redundancy between non-agricultural UDLs and is the sum of the non-agricultural UDLs divided by the non-agricultural composite. Then the calculations described in Scenario three are completed using the updated UDLs.

**Scenario 4 and 5: Considering life history**

Species life history information is incorporated in Scenario 4 and 5. Scenario four considers if the species will be found on potential use sites or exclusively off the use sites. These determinations are informed by documentation from the Services (e.g., Recovery Plan, 5-year Reviews). Scenario 5, a proof of concept, considers if a species is found exclusively in specific habitats, but is not currently used in the contiguous U.S.

Our analysis does not consider life history data. The inclusion of life history data of this nature would ultimately increase the number of species receiving an NLAA determination through the spatial analysis.

**Step 5. Results Co-occurrence Analysis**

The results of the co-occurrence analysis provide the percent of the species range (or critical habitat) that overlaps with each UDL. Figs. S61 and S62 provide examples of analyses. The first shows a species where inclusion of usage data significantly reduced the overlap (i.e., high impact of pesticide usage on outcomes), while the second shows a species where usage data had little impact on overlap (i.e. low income of pesticide usage on outcomes).

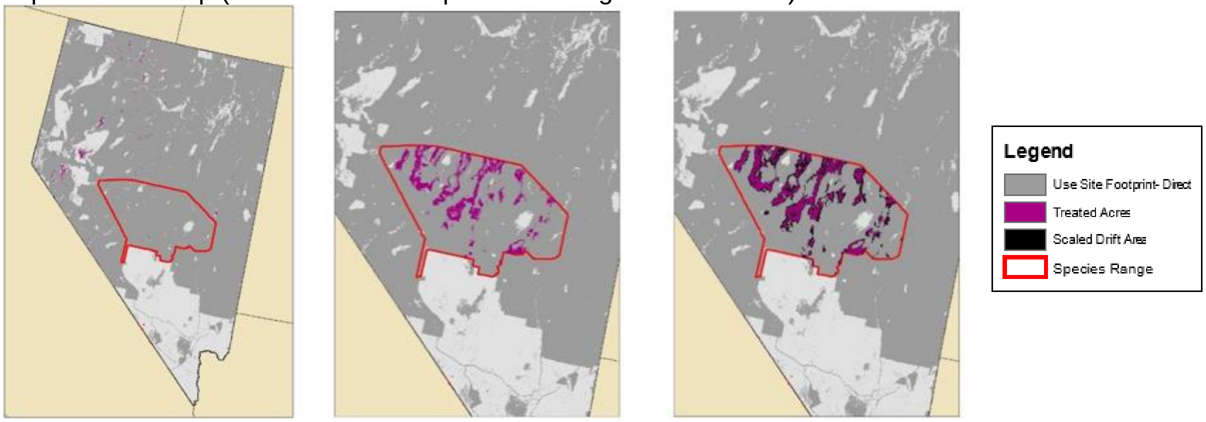

**Fig. S62.** Example of a species with low impact of pesticide usage

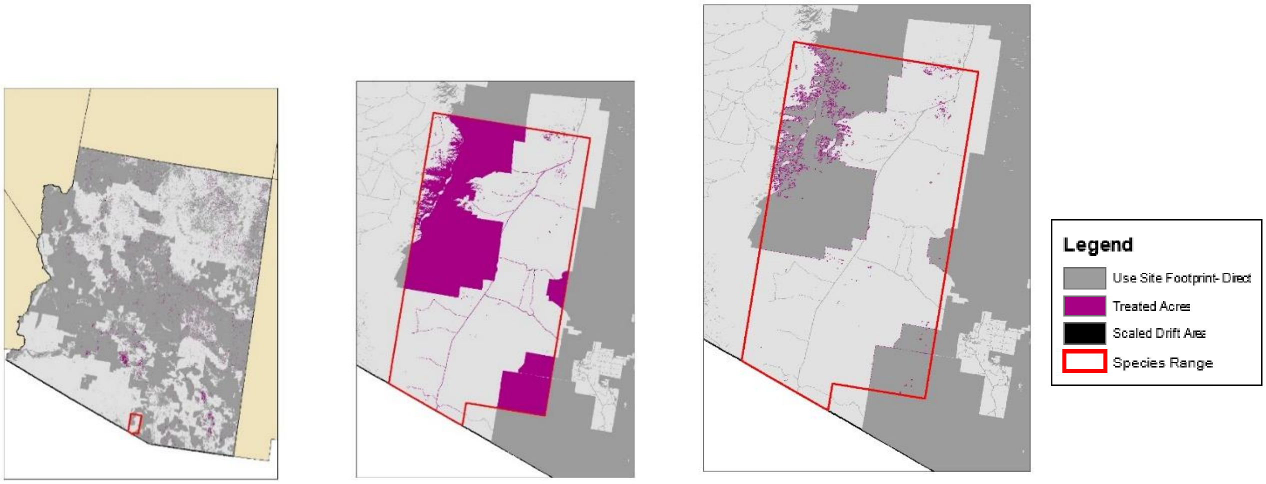

**Fig. S62.** Example of a species with low impact of pesticide usage

#### Data Sources and Citations

***S. E. Eikenberry, G. Iacona, E. Murphy, G. Watson, L. Gerber. Identifying opportunities for high resolution pesticide usage data to improve the efficiency of endangered species pesticide risk assessment: Data, notebooks, and results. Dryad, Dataset, https://datadryad.org/stash/share/GdNHpEf1xMa9ZRVA4Eo2xiTWlF7cXrzNeQzzhcmLNmU, (2022).**

****U.S. Environmental Protection Agency, 2021. Final national level listed species biological evaluation for carbaryl. Retrieved from https://www.epa.gov/endangered-species/final-national-levellisted-species-biological-evaluation-carbaryl**

*** Data and apps referenced in the SI**

**** Key reference that is summarized in Section 10**

Amos, J.J., C.M. Holmes, C.G. Hoogeweg, and S.A. Kay. 2010. Development of Datasets to Meet USEPA Threatened and Endangered Species Proximity to Potential Use Sites Data Requirements. Report Number: 437.01-Overview. Prepared by Waterborne Environmental, Inc. for the Generic Endangered Species Task Force.

Baker, N.T., and Capel, P.D., 2011. Environmental factors that influence the location of crop agriculture in the conterminous United States: U.S. Geological Survey Scientific Investigations Report 2011–5108, 72 p.

Bonneville Power Administration GIS, 2015. Bonneville Power Administration Right of Way (BPA ROW), 2015, https://bpagis.maps.arcgis.com/home/

Bureau of Land Management (BLM) Grazing Allotments BLM GIS, Grazing Allotment Boundaries, 20140112 https://data.doi.gov/dataset/blm-grazing-allotment-polygons

Dun & Bradstreet, Agriculture, US, 2012, Dun & Bradstreet, SEGS, Short Hills, NJ, 2013/04/08

ESRI StreetMap North America Railroads. ESRI, StreetMap North America, Redlands, CA 20100531. EPA Access ftp://cook.rtp.epa.gov/data/ESRI_DATA_AND_MAPS/

NAVTEQ Street Data. NAVTEQ 2013 Streets, Chicago, IL, 20131001EPA Access ftp://cook.rtp.epa.gov/data/NAVTEQ/2013/

Homer, C.G., Dewitz, J.A., Yang, L., Jin, S., Danielson, P., Xian, G., Coulston, J., Herold, N.D., Wickham, J.D., and Megown, K., 2015. Completion of the 2011 National Land Cover Database for the conterminous United States-Representing a decade of land cover change information. Photogrammetric Engineering and Remote Sensing, v. 81, no. 5, p. 345-354 (National Land Cover Dataset (NLCD) 2011)

National Oceanic and Atmospheric Administration, Coastal Services Center. 1995-present. The Coastal Change Analysis Program (C-CAP) Regional Land Cover. Charleston, SC: NOAA Coastal Services Center.

Accessed at https://coast.noaa.gov/digitalcoast/data/ (National Oceanic and Atmospheric Administration (NOAA) Coastal Change Analysis Program (CCAP))

United States Census Bureau's Topologically Integrated Geographic Encoding and Referencing database (TIGER) 2015 TIGER/Line Shapefiles (machine readable data files) / prepared by the U.S. Census Bureau, 2015, https://www.census.gov/geographies/mapping-files/time-series/geo/tiger-geodatabase-file.html

United States Department of Agriculture (USDA), National Agricultural Statistics Service (NASS), Research and Development Division (RDD), Geospatial Information Branch (GIB), Spatial Analysis Research Section (SARS), Cropland Data Layer for the United States, https://www.nass.usda.gov/Research_and_Science/Cropland/SARS1a.php (United States Department of Agriculture Cropland Data Layer (CDL) 2013-2017)

USDA Forest Service, Administrative Forest Boundaries, "S_USA.AdministrativeForest", 20151027, http://data.fs.usda.gov/geodata/edw/datasets.php (United States Forest Service Administrative Boundaries)

USFS Range Allotment Boundaries, NationalAllotmentFeatureClassAlbers, Rangeland Management Unit, 20140916. Provided by Gene O'Donnell, USFS Geospatial Interface Account Manager, (United States Forest Service Grazing Allotments)

US Geological Survey, Gap Analysis Program (GAP). May 2011. National Land Cover, Version 2 (United States Geological Survey GAP Land Cover Data (USGS GAP))

US Geological Survey, Gap Analysis Program (GAP). November 2012. Protected Areas Database of the United States (PADUS), version 1.3 Combined Feature Class. (United States Geological Survey GAP Protected Areas Database (USGS GAP PAD-US))

LANDFIRE, 2012, Existing Vegetation Type Layer, LANDFIRE 1.3.0, U.S. Department of the Interior, Geological Survey. Accessed 15 July 2015 at https://www.landfire.gov/version_comparison.php (United States Geological Survey LandFire Existing Vegetation Type (USGS LandFire EVT))

LANDFIRE, 2012, Public Events GeoDatabase, LANDFIRE 1.3.0, U.S. Department of the Interior, Geological Survey. Accessed 15 July 2015 at https://www.landfire.gov/version_comparison.php (United States Geological Survey LandFire Public Events GeoDatabase (USGS LandFire Events))

Query used to extract species from TESS: https://ecos.fws.gov/services/TessQuery?request=query&xquery=/SPECIES_DETAIL

Website for designated critical habitat: (http://ecos.fws.gov/crithab)

State of Hawaii, Plant Industry Division, 2016. Restricted Use Pesticides (RUP) Sales, http://hdoa.hawaii.gov/pi/ruplist/

U.S. Geological Survey Gap Analysis Program, 20160513, GAP/LANDFIRE National Terrestrial Ecosystems 2011: U.S. Geological Survey: Boise, ID, http://gapanalysis.usgs.gov/gaplandcover/. doi:10.5066/F7ZS2TM0.

USFWS, personal communication, November 2019.
